# The *ORGAN SIZE* (*ORG*) locus contributes to isometric gigantism in domesticated tomato

**DOI:** 10.1101/2021.04.08.439112

**Authors:** Mateus Henrique Vicente, Kyle MacLeod, Cassia Regina Fernandes Figueiredo, Antonio Vargas de Oliveira Figueira, Fady Mohareb, Zoltán Kevei, Andrew J. Thompson, Agustin Zsögön, Lázaro Eustáquio Pereira Peres

## Abstract

Gigantism is a key component of the domestication syndrome, a suite of traits that differentiates crops from their wild relatives. Allometric gigantism is strongly marked in horticultural crops, causing disproportionate increases in the size of edible parts such as stems, leaves or fruits. Tomato (*Solanum lycopersicum*) has attracted attention as a model for fruit gigantism, and many genes have been described controlling this trait. However, the genetic basis of a corresponding increase in size of vegetative organs contributing to isometric gigantism, has remained relatively unexplored. Here, we identified a 0.4 Mbp region on chromosome 7 in introgression lines (ILs) from the wild species *Solanum pennellii* in two different tomato genetic backgrounds (cv. M82 and cv. Micro-Tom) that controls vegetative and reproductive organ size in tomato. The locus, named *ORGAN SIZE* (*ORG*), was fine-mapped using genotype-by-sequencing. A survey of literature revealed that *ORG* overlaps with previously mapped QTLs controlling tomato fruit weight during domestication. Alleles from the wild species led to reduced cell number in different organs, which was partially compensated by greater cell expansion in leaves but not in fruits. The result was a proportional reduction in leaf, flower and fruit size in the ILs harbouring the wild alleles. These findings suggest that selection for large fruit during domestication also tends to select for increases in leaf size by influencing cell division. Since leaf size is relevant for both source-sink balance and crop adaptation to different environments, the discovery of *ORG* could allow fine-tuning of these parameters.

**One sentence summary:** A locus that controls isometric size increase in vegetative and reproductive organs of tomato through changes in cell division

## Introduction

The domestication syndrome is the suite of phenotypic changes that occurred through artificial selection to transform wild species into crops (Evans 1996). Some of the most commonly found traits in crops are increased apical dominance, determinate growth and loss of natural seed dispersal (Meyer et al. 2012; Denham et al. 2020). An increase in the size of certain organs, or gigantism, is also widespread, particularly in horticultural crops (Schwanitz 1957). Gigantism can be isometric, *i*.*e*. a proportional increase in all body parts, but most generally occurs through allometric alterations in the relative size of certain plant structures (Niklas 2004). A prime example is the species *Brassica oleracea*, where multiple cultivated strains were produced through artificial selection on the differential growth of edible organs such as stems (kohlrabi), buds (cabbage, Brussels sprouts), leaves (kale) and flowers (broccoli, cauliflower) (Prakash et al. 2011). Although increased organ size can be explained by alterations in cell division and expansion (Krizek 2009), it also requires developmental alterations to transform larger organs into stronger photosynthetic sources or sinks (Gifford et al 1984). Given that photosynthesis as a biochemical process has not been improved by crop domestication or breeding (Orr et al. 2017; Batista-Silva et al. 2020), most of the genetic gains in productivity have occurred indirectly through changes in plant development (Greenland et al. 1997; Zsögön and Peres 2018).

In tomato (*Solanum lycopersicum* L.), gigantism is evidenced in the phenomenal increase in fruit size when compared to its wild progenitor *S. pimpinellifolium* (Tanksley 2004). The genetic basis of fruit size control has attracted considerable attention (reviewed in Azzi *et al*., 2015). Increased fruit size in tomato involves mutations in multiple loci, some of which have been characterized at the molecular level, for instance *fruit weight 2*.*2* (*fw2*.*2*), *fw3*.*2, fw11*.*3, fasciated* (*fas*), *locule number* (*lc*) and *EXCESSIVE NUMBER OF FLORAL ORGANS* (*ENO*). All of them are involved in fundamental processes of plant developmental such as cell division, expansion and differentiation. The *FW2*.*2* gene is a negative regulator of cell division responsible for up to 30% of the increase in fruit size when comparing lines harbouring small- and big-fruit alleles (Frary et al. 2000). *FW3*.*2* and *FW11*.*3* were identified as a P450 enzyme of the CYP78A subfamily (*SlKLUH*) and a *Cell Size Regulator* (*CSR*), controlling cell division and expansion, respectively (Chakrabarti et al. 2013; Mu et al. 2017). Unlike *fw2*.*2, fw3*.*2* and *fw11*.*3*, which mostly affect fruit size, *fas* and *lc* also control fruit shape. The big-fruit *fas* and *lc* alleles increase the number of carpels, altering cell diferentiation through the CLAVATA3-WUSCHEL module (Schoof et al. 2000). The increase in the number of carpels often results in larger and wider fruits with many locules and pronounced ribbing (Lippman and Tanksley 2001; van der Knaap and Tanksley 2003). The *lc* mutant phenotype is caused by two single-nucleotide polymorphisms (SNPs) downstream of the coding region of the *WUSCHEL* (*WUS*) gene (Muños et al., 2011). The *fas* mutation is a partial loss of expression caused by a chromosome inversion with a break point in the vicinity of the *CLAVATA3* (*CLV3*) gene (Xu et al 2015), a negative regulator of *WUS* (Schoof et al. 2000). Lastly, *ENO* is an AP2/ERF transcription factor that interacts synergistically with *lc* and *fas*, causing a substantial increase of the *WUS* expression domain, which results in enlarged floral meristems (Fernández-Lozano et al., 2015; Yuste-Lisbona et al., 2020). Thus, the *ENO* domestication allele (a promoter deletion that knocks down its expression) also affects stem cell fate, giving rise to multilocular fruits that derive from the larger floral meristem.

Compared with the genetic regulation of fruit growth, relatively little is known about the control of vegetative organ size. In many crops, including tomato (Supp Fig. S1) but also peppers (Jarret et al., 2019), sunflower (Warburton et al., 2017), soybeans (Kofsky et al., 2018) and common beans (Herron et al., 2020), domestication entailed the selection of plants with bigger shoots and leaves. In tomato, the proportional increase in the size of vegetative parts is likely to be a component of isometric gigantism during domestication. Herein, we hypothesized that if vegetative gigantism is under genetic control, the wild species’ alleles leading to reduced organ size could be found through wide crosses between cultivated tomato and its wild relative species. We selected *S. pennellii* as a wild parental, due to its annotated genome sequence (Bolger et al. 2014) and its rich repertoire of genomic tools, such as fully sequenced introgression lines (Alseekh et al. 2013; Chitwood et al. 2014). We crossed it to the cultivated tomato cv. Micro-Tom (MT) and after successive backcrosses and phenotypic selection, we isolated an introgression line with reduced vegetative and reproductive organs compared to the recurrent parental MT. We mapped this introgression to chromosome *7* and named the locus *ORGAN SIZE* (*ORG*). We show that *ORG* leads to reduced organ size through changes in cell division, and that it segregates as a monogenic, semi-dominant Mendelian locus. Our fine mapping results show that the *ORG* candidate genes overlap a previously described domestication sweep (Lin et al. 2014). We speculate on the impact of this locus in the tomato domestication syndrome and discuss its potential exploitation for crop breeding.

## Results

### Natural genetic variation for leaf size in tomato

Compared to domesticated tomato cultivars, most wild relatives of the tomato have small leaves (Supplemental Figure S1). Thus, we decided to look for a genetic determinant of leaf size in the wild species. We crossed *S. pennellii* to the cultivated tomato cv. Micro-Tom (MT). Upon self-fertilization of the F_1_ population, we selected F_2_ plants with small leaves, from which we collected pollen to backcross (BC) to MT. After six rounds of backcrossing to the recurrent parental (MT), self-fertilization (BC_6_F_2_), phenotypic screening, and further self-fertilization (BC_6_F_n_), we produced an introgression line (IL) with reduced leaf size in the MT background, which we called *ORGAN SIZE* (*ORG*) (Figure 1). *ORG* plants show a very conspicuous phenotype for leaf size: the difference in leaf size between MT and *ORG* was consistent across all leaves and developmental stages (Figure 1). Monogenic segregation of *ORG* was verified on a segregating population of MT and *ORG*. We determined leaf size in F_1_ hybrids between MT and *ORG*, and the intermediate phenotype suggested that *ORG* behaves as a semi-dominant gene (Supplemental Figure S2).

**Figure 1.**
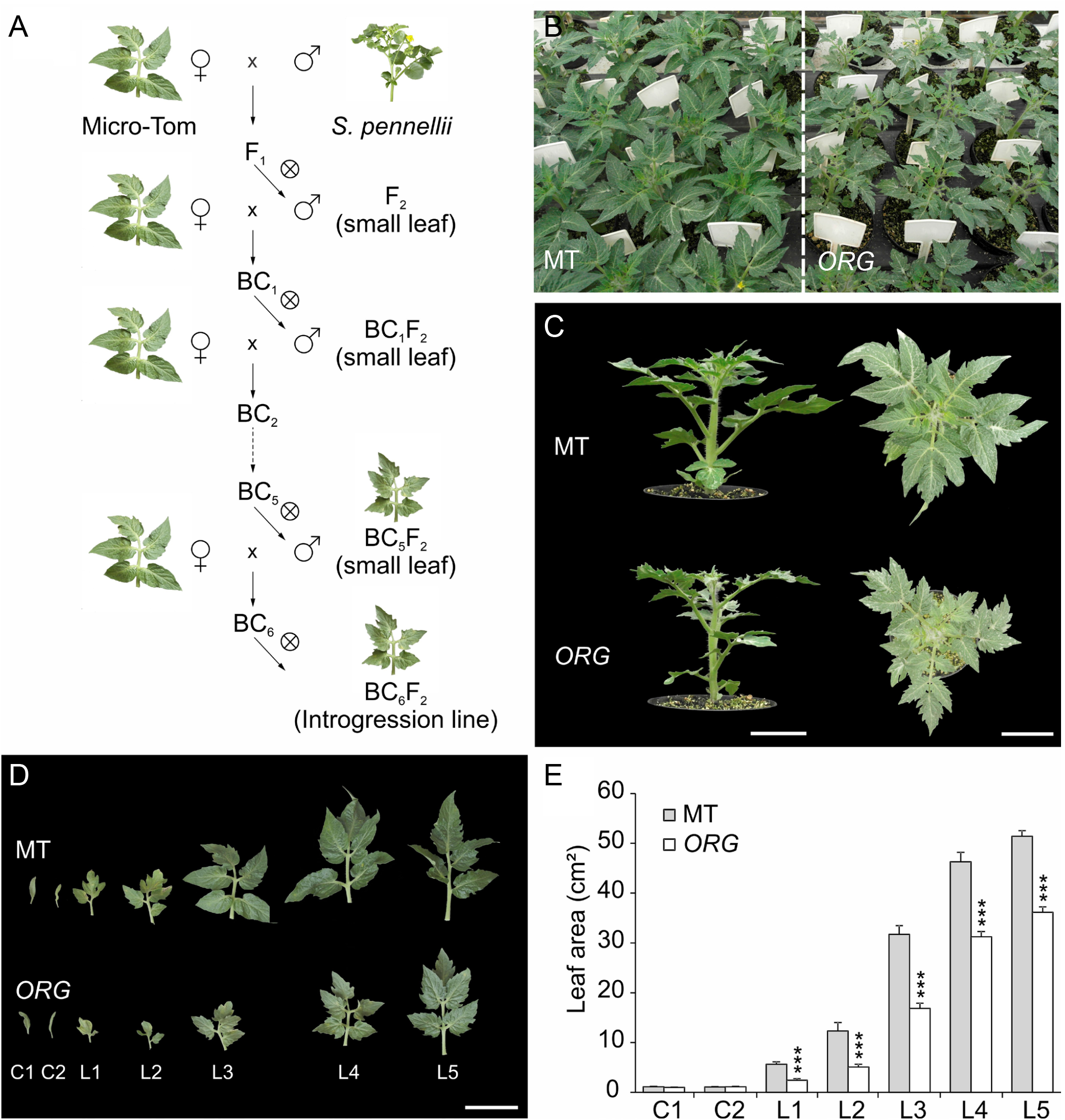
A tomato introgression line (IL) from *S. pennellii* with reduced vegetative organs (*ORG*) size. **(a)** Crossing scheme to create an introgression line with smaller leaves in the tomato cv Micro-Tom (MT) background **(b)** Representative population of MT (left) and *ORG* (right) plants, 25 days after germination (dag). **(c)** Side and top view of MT (top) and *ORG* (bottom) plants. **(d)** Leaf series of MT (top) and *ORG* (bottom) genotypes from cotyledons (C1) to fifth leaf (L5). Scale bar=5 cm. **(e)** Leaf area of the leaf series of MT (gray bar) and *ORG* (white bar) plants, 40 dag. Data are mean ± s.e.m. (n=14 leaves). Statistical significance was tested by Student’s *t-*test (\*\*\**p*<*0*.*001*).

### The smaller leaf size in ORG is caused by reduced cell division

Change in organ size is due to either altered cell proliferation or expansion, or a combination of both (Krizek 2009). We analysed *ORG* leaves and found enlarged epidermal and mesophyll cells compared to MT (Supplemental Figure S3). This suggests that the smaller leaves of *ORG* are caused by reduced cell proliferation as evidenced by cell number and density of *ORG* compared to MT (Supplemental Figure S3). The greater palisade parenchyma cell size promoted an increase in leaf thickness in *ORG*. We next performed a time course analysis of reproductive growth starting eight days before anthesis and until 16 days after anthesis and verified a decrease in the size of styles, ovaries and fruits in *ORG* (Figure 2). As in the case of leaves, the reduction was caused by lower cell numbers, which we verified as a reduced number of cell layers in the pericarp. The ovary cells of *ORG* were also smaller than MT cells at anthesis and post-anthesis. Other floral organs, namely, petals and sepals, were also reduced in *ORG* flowers compared to MT (Supplemental Figure S4). The reduced size of floral organs may have strong consequences on fruit development, given their impact on ovary size (Supplemental Figure S4e-h).

**Figure 2.**
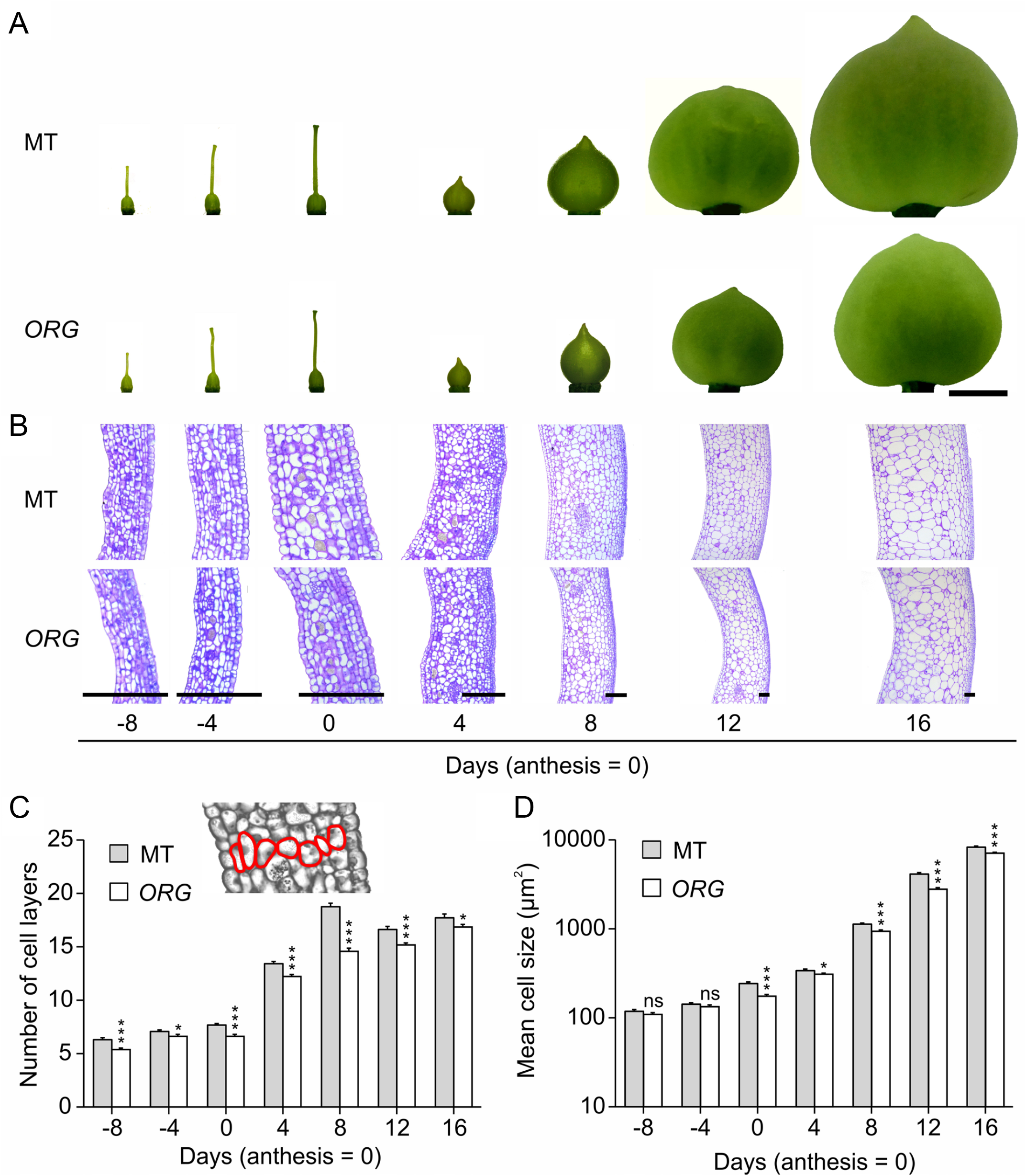
*ORG* affects cell number and size during fruit development. **(a)** Developing ovary/fruit at −12, −8, −4, 0, 4, 8, 12 and 16 days (anthesis = 0). MT (top) and *ORG* (botton). Scale bar=5mm. **(b)** Longitudinal sections of MT (top) and *ORG* (bottom) pericarp at −12, −8, − 4, 0, 4, 8, 12 and 16 days (anthesis = 0). Scale bar = 150µm. **(c)** Time course of the number of cell layers in the longitudinal sections of MT (gray bar) and *ORG* (white bar) ovary/fruit pericarp. Insert in top of this figure represents how the counting of the cells was performed and red lines delimited cell perimeter (n=30). **(d)** Time course of cell area in the cell layers of MT (gray bar) and *ORG* (white bar) (n=30). Data are mean ± s.e.m. Statistical significance was tested by Student *t-*test (\**p*<*0*.*05*, \*\*\**p*<*0*.*001, ns* indicates non-significant differences).

### Fruit weight and yield are reduced in ORG

The size and shape of the ovary before anthesis is strongly correlated with the final size and shape of the fruit (Grandillo et al., 1999; Azzi et al., 2015). Thus, we next analysed the potential impact of *ORG* on fruit development. Fruit set was reduced in heterostylic *ORG* flowers, so we hand-pollinated emasculated MT and *ORG* flowers in a reciprocal cross. Several ovaries per plant were pollinated, but after fruit set confirmation (five days after pollination), we performed selective fruit removal to allow only five fruits to set on each plant. The presence of *ORG* ovaries had a substantial impact on the final fruit size regardless of pollen origin (Figure 3). Fruit weight was 31-37% lower in *ORG* than in MT (*P* < 0.0001, Supplemental Table S2). *ORG* fruits have higher total soluble solids content (°Brix) compared to MT (Supplemental Figure S5). We further observed that *ORG* had a similar frequency of locule number per fruit and reduced seed number (Supplemental Figure S5). Reciprocal crosses indicated that the reduction in seed number is determined by *ORG* ovaries rather than pollen (Supplemental Figure 5c).

**Figure 3.**
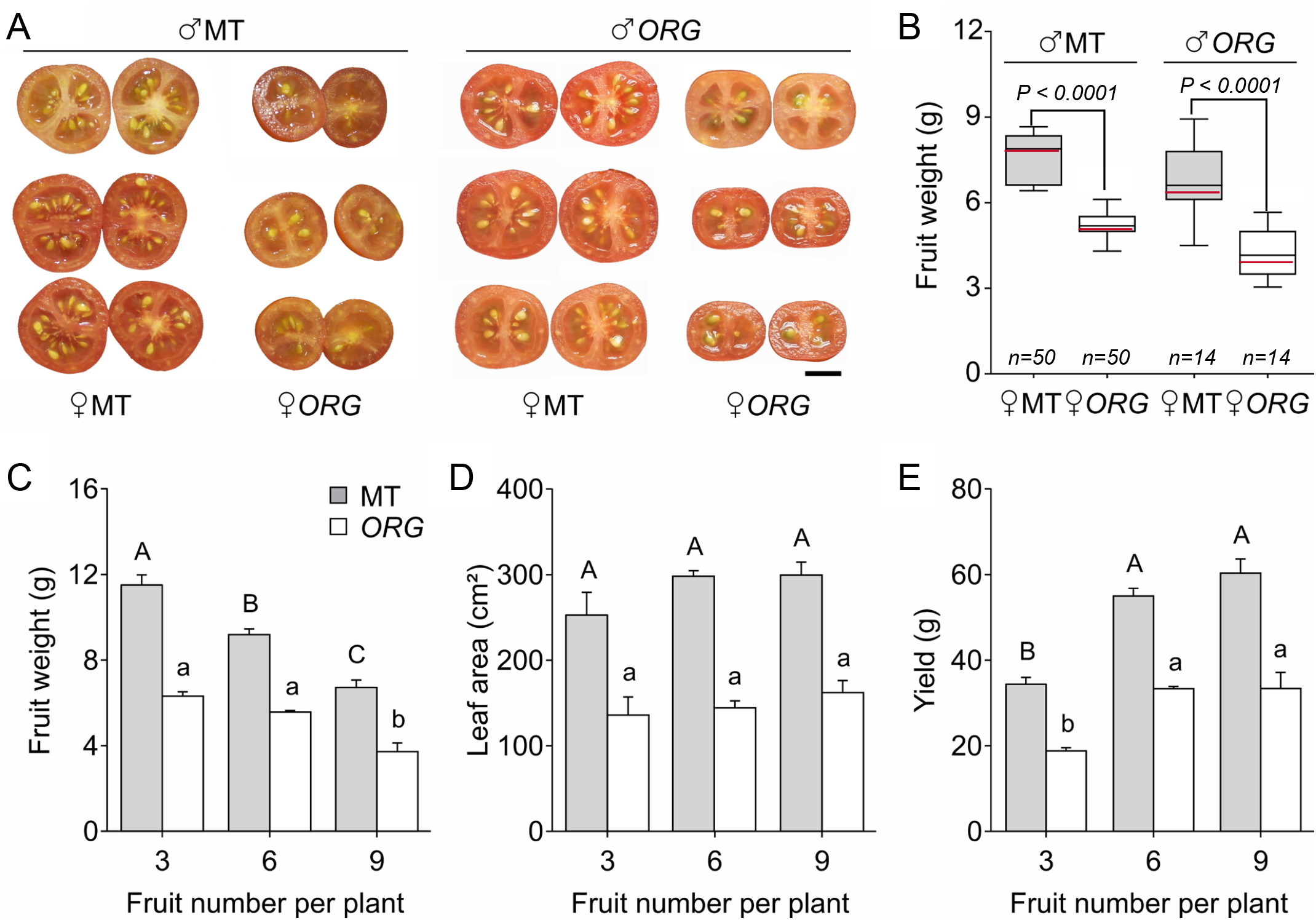
Fruit growth and source-sink relationships are altered in *ORG*. **(a)** Representative MT (♀, left) and *ORG* (♀, right) ripe fruits pollinated with MT (♂, left) and *ORG* (♂, right) pollen. Scale bar=1 cm. **(b)** Mean (red) and median (black) values of fruit weight of MT (gray box) and *ORG* (white box) ripe fruits pollinated with MT (n=10) and *ORG* (n=14) pollen. **(c)** Frequency of locule number per fruit in MT and *ORG* fruits (n=125). **(d)** Seeds per fruit of MT and *ORG* pollinated with MT (n=11) and *ORG* (n=15) pollen. Data are mean±s.e.m. Statistical significance was tested by Student’s *t*-test (\*\*\**p*<*0*.*001*). **(e-g)** Average values of fruits weight (e), leaf area (f) and yield (g) from MT (gray bar) and *ORG* (white bar) plants pruned to three, six and nine fruits (n=6 plants per treatment). Data are mean±s.e.m. Different capital and lowercase letters on the symbols indicate significant differences by Tukey’s test (*p*<*0*.*001*) between the treatments in MT and *ORG* genotypes, respectively.

We next addressed the possibility that reduced fruit size could be the consequence of altered photosynthetic source-sink relationships due to reduced leaf area. We thus manipulated the plants to maintain the availability of sources (leaves) constant and altered the source:sink ratio by changing the number of sinks (fruits). Three treatments were performed: either three, six or nine fruits were allowed to set on each plant. To ensure that additional sinks did not interfere in the results, we also pruned all the plants to remove side shoots. The results are summarized on Figure 3c-e. *ORG* plants produced consistently smaller fruits than MT in all treatments (Figure 3). The increase in fruit number, from three to six, promoted a reduction in fruit weight only in MT plants, suggesting that leaf area was a limiting factor to the final fruit weight in MT, since the leaf area was similar in both experimental conditions (Figure 3). On the other hand, when the number of fruits was increased from six to nine, there was a reduction in the final fruit weight for both genotypes. These results suggest that the smaller leaf size of *ORG* could also account for its reduced fruit size, but only under full fruit load. Therefore, the primary cause of the reduced fruit size in *ORG* is likely a direct effect of this organ development since the pre-anthesis (Fig. 2c). In addition, the presence of the ORG introgression reduced the yield in all treatments.

### Expression patterns are altered in genes related to cell division and expansion in ORG

The results described so far suggest that the transcriptional activity of genes involved in the control of cell division and expansion could be altered in *ORG*. To assess this, we extracted mRNA from ovaries/fruits at −8, −4, 0, 4 and 8 days pre/post anthesis, and fruit pericarps at 12 and 16 days to analyse the transcriptional profile of a set of genes related to the control of cell division: *CYCLIN B2;1* (Solyc02g082820), *FW2*.*2* (Solyc02g090730), *FW3*.*2/SlKLUH* (Solyc03g114940) and *EXPANSIN PRECURSOR 5* (Solyc02g088100).

In ovary/fruit tissues, we verified that the mRNA levels of the cell-division genes *CYCB2;1* and *FW3*.*2* showed greatest expression in both genotypes at 4 days pre-anthesis (Figure 4). *CYCB2;1* was higher in MT than *ORG* especially in pre-anthesis and anthesis stages (at −4, −8, and 0 days), while *FW3*.*2* was higher in anthesis and post-anthesis stages (at 0, 4 and 12 days). On the other hand, *FW2*.*2*, another cell-division gene, but a negative regulator, was highly expressed at 4 and 8 days post-anthesis in both genotypes. Quantitative variation in *FW2*.*2* expression was observed pre- and post-anthesis between genotypes (at −4 and 8 days, respectively), whereas *ORG* ovaries showed significant increased levels of this transcript compared than MT (Figure 4). After 4 days post-anthesis, the expression of the cell-expansion gene *EXPA5*, a member of the α-expansin gene family, increased in in both genotypes (Figure 4). However, ovaries of *ORG* plants displayed a significant decrease in the expression of this gene at anthesis compared to MT. Similar behavior was observed 16 DPA.

**Figure 4.**
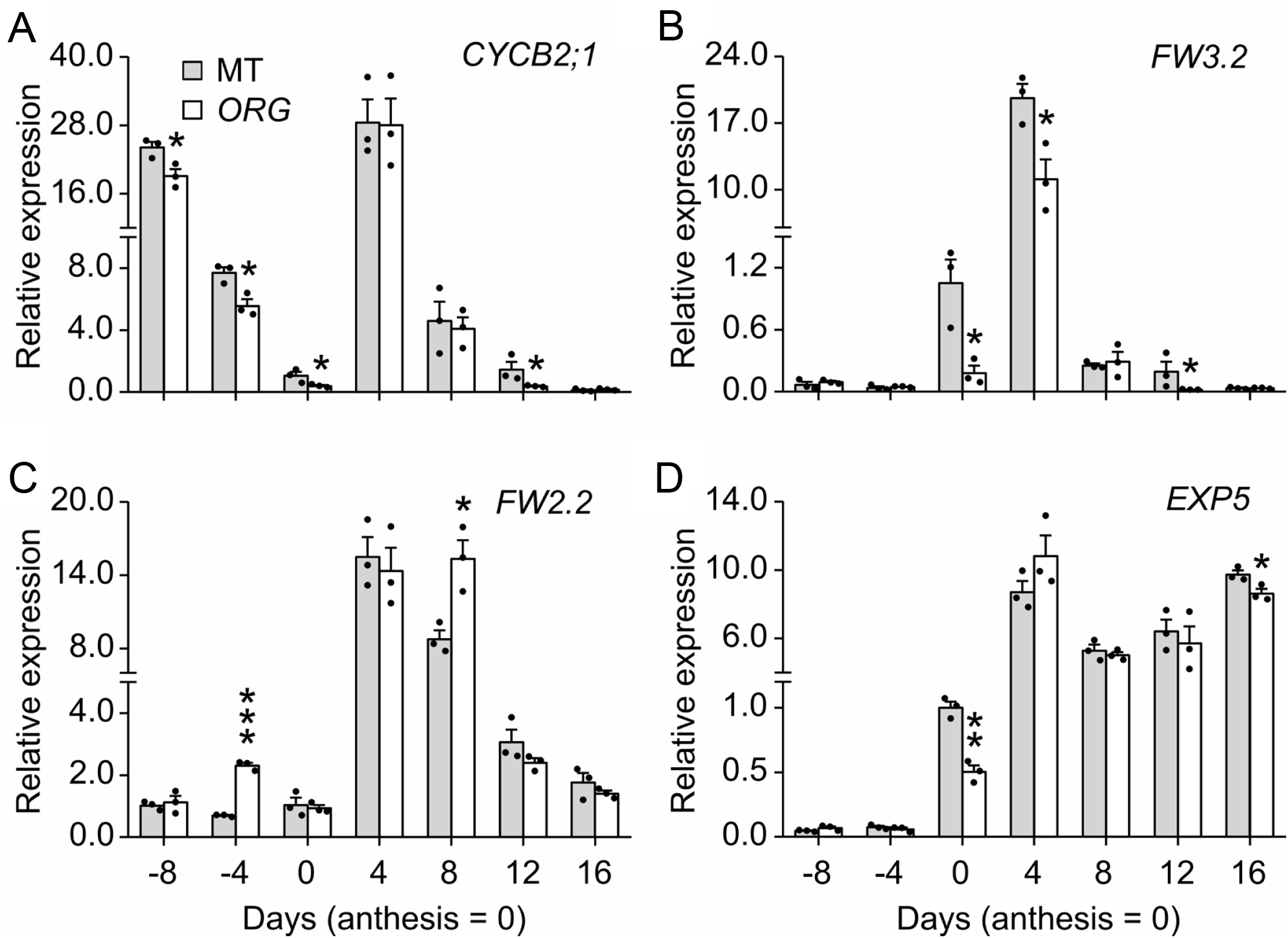
Altered patterns of gene expression in *ORG*. Time course of transcript levels of cell division- and expansion-related genes in ovaries/fruits of MT (gray bar) and *ORG* (white bar) genotypes. Relative (to actin control) transcript levels of *CYCB2;1* **(a)**, *FW2*.*2* **(b)**, *FW3*.*2* **(c)** and *EXP5* **(d)** in ovaries/fruit at −8, −4, 0, 4, 8 days and fruit pericarp at 12 and 16 days (anthesis = 0). Data are mean±s.e.m (n=3 biological replicates indicated with black dots). Statistical significance was tested by Student’s *t-*test (\**p*<*0*.*05*, \*\**p*<*0*.*01*, \*\*\**p*<*0*.*001*).

### The ORG locus is located on chromosome 7

We next conducted a genotyping by sequencing (GBS) analysis to determine the size and location of the *S. pennellii* introgression in *ORG*. The results show a discrete region in the terminal end of the long arm of chromosome *7* encompassing ∼11 Mb (Figure 5). No further segments of *S. pennellii* genome were found on other chromosomes. Based on the SL2.50 tomato genome annotation, the introgression region contains 1169 genes. A closer look at the introgressed region revealed a small double recombination, from position 64,826,717 to 65,444,176, encompassing 78 tomato genes which score as *S. lycopersicum* (Figure 5b).

**Figure 5.**
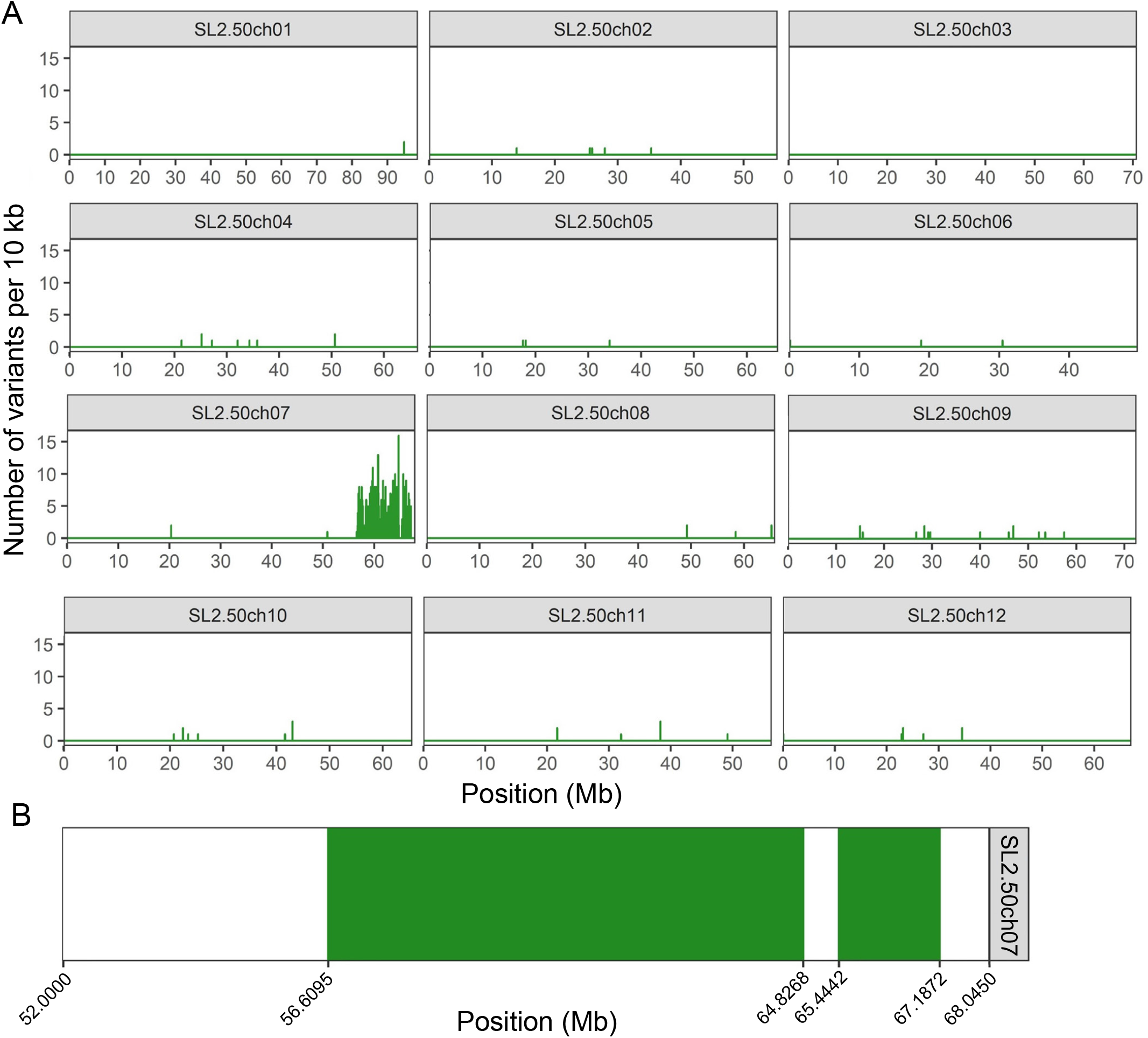
GBS defines the span of the introgression in the *ORG* introgression line (IL). **(a)** Genome-wide density of unique variants shared between *ORG* and *S. pennellii* LA716 in the genetic background of tomato cv Micro-Tom. **(b)** Close up view of the introgression on chromosome 7.

### Fine-mapping of OS using introgression lines

To reduce the list of candidate genes for *ORG*, we next analysed two other introgression lines (ILs) of *S. pennellii* in the MT background previously generated in our laboratory: *Brilliant corolla* (*Bco)* and *Regeneration 7H* (*Rg7H)*, both of which partially overlap either end of the *ORG* introgression (Figure 6). We used the span of the introgressions in *Bco* and *Rg7H* and the extent of their overlap with *ORG* (Supplemental Figure 6 for *Bco* and Pinto *et al*., 2017 for *Rg7H*) to narrow down the candidate region for *ORG*. Given that neither of these ILs show the reduced organ phenotype of *ORG*, the resulting candidate region is located between positions 65,444,176 and 66,373,175 (Figure 6a).

**Figure 6.**
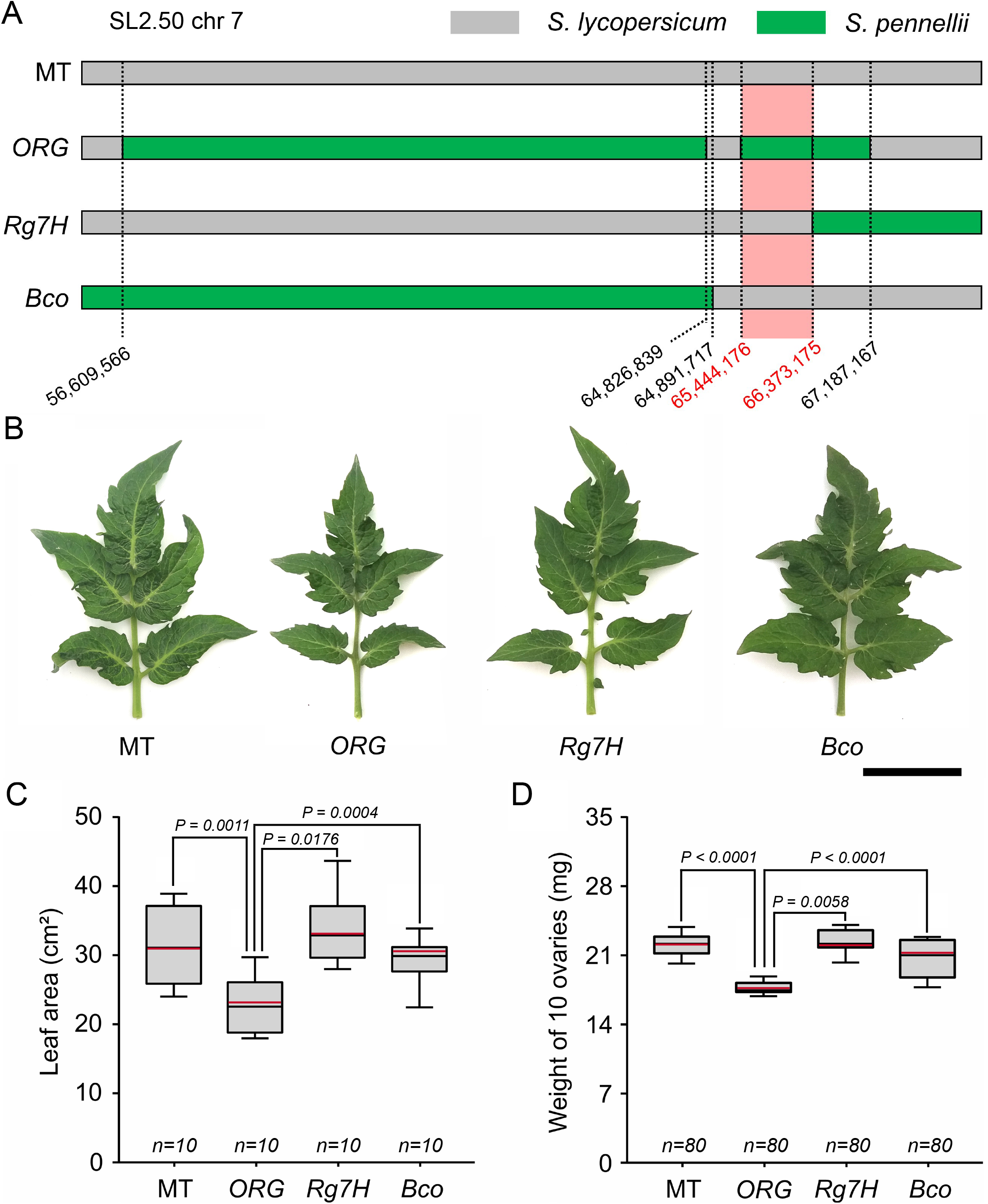
Mapping refines the candidate region for *ORG*. **(a)** Two introgression lines (ILs) in the tomato cv Micro-Tom (MT) background that contain different segments from *S. pennellii* on chromosome 7 (*Bco* and *Rg7H*) were mapped to refine the candidate region harboring the *ORG* locus (red segment). **(b)** Representative leaf of MT, *ORG, Rg7H* and *Bco* genotypes. Scale bar = 5 cm. **(c-d)** Leaf area (c) and ovary weight (d) of MT, *ORG, Rg7H* and *Bco* (n=10). Statistical significance was tested by Tukey’s test (*p*<*0*.*05*). Different letters indicate significant difference between genotypes.

We took advantage of the existing collection of ILs from *S. pennellii* in tomato cv. M82 as a tool to further refine the above chromosome location (Zamir and Eshed 1994; 1995). The introgressions were precisely delimited by sequencing by Chitwood et al. (2014), who also characterized terminal and lateral leaflet size in the ILs. Their results revealed the existence of a QTL for reduced leaflet size on both IL7-2 and IL7-3 (Figure 7a-b). We also cultivated ILs harbouring *S. pennellii* genomic segments on chromosome 7 (IL7-1; IL7-2, IL7-3; IL7-4 and IL7-5) and determined their leaf and ovary size. We found a reduction in the ovaries of both IL7-2 and IL7-3, compared to M82, but under our growth conditions only IL7-2 showed consistently smaller leaves than the parental line (Supplemental Figure S7). We found a discrepancy between the Chitwood et al. dataset and ours for leaf size on IL7-1, but the consistently smaller pistils in IL7-2 and IL7-3 helped us delimit the right border of the candidate region to 65,865,655, narrowing the interval to 421,479 bp (Figure 7c).

**Figure 7.**
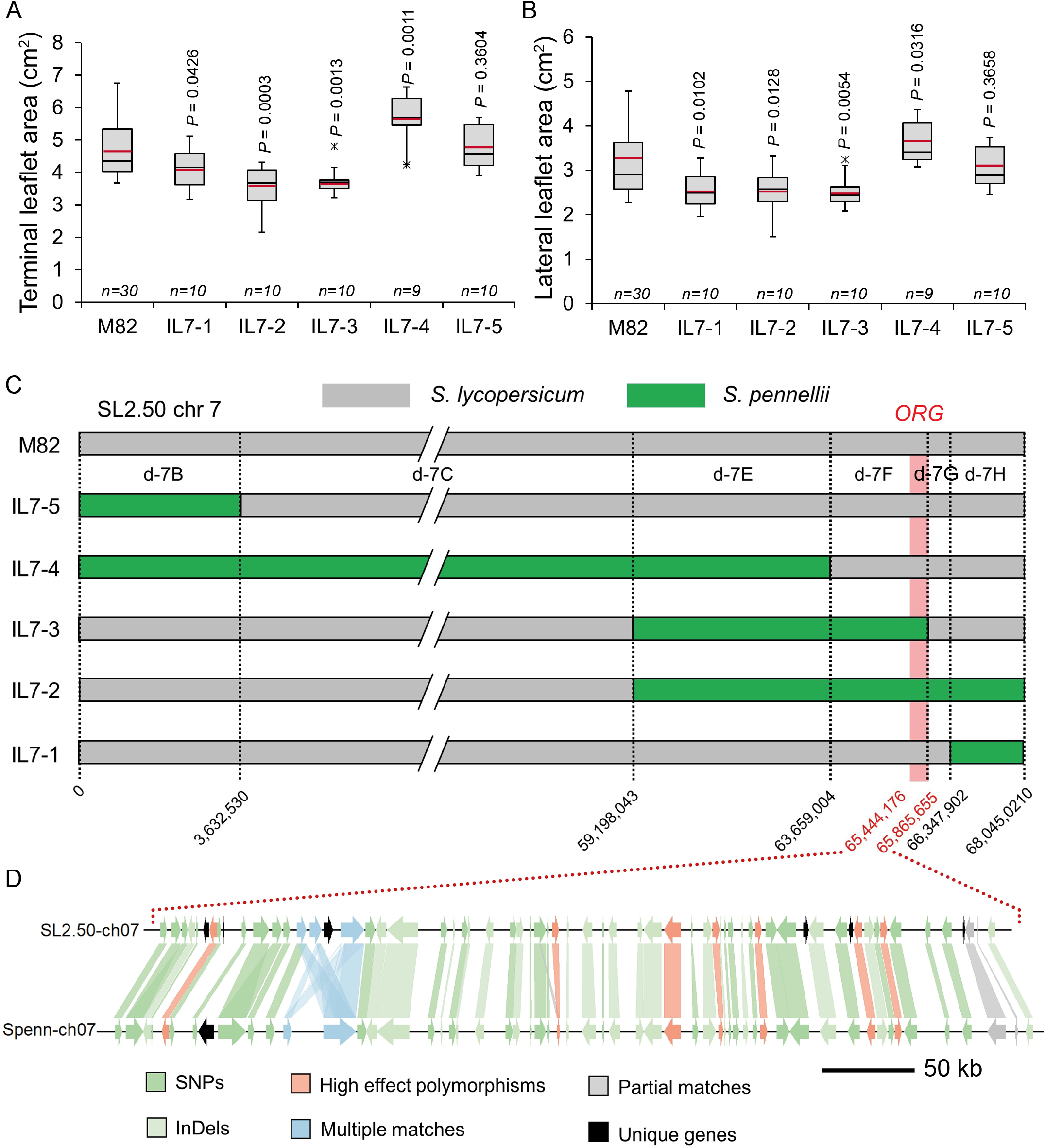
Analysis of the genomic region containing *ORG*. Terminal **(a)** and lateral **(b)** leaflet area of M82 and chromosome *7* introgression lines (ILs) from *S. pennellii*. Statistical significance was tested by ANOVA followed by Tukey’s HSD test. Redrawn from Chitwood et al. (2014). **(c)** Chromosomal position of *S. pennellii* genomes segments in tomato cv. M82 background in chromosome *7*. The location of the *ORG* candidate region is shown in red. **(d)** Synteny plot of the coding sequences (CDS) within the *ORG* region between *S. lycopersicum* and *S. pennellii* genomes. The similarity between the CDS of *S. lycopersicum* (SL2.50) and *S. pennellii* (Spenn) were tested with BLAST+ and variant effect prediction was obtained from the resequenced dataset (Aflitos *et al*. 2014). Key: Dark green, CDS that match with a high level of similarity, but *S. pennellii* alleles contain single nucleotide polymorphisms (SNPs). Light green, *S. pennellii* alleles contain insertions and deletions (InDels). Red, *S. pennellii* alleles contain variants predicted to cause loss of function. Blue, complex relationship between *S. lycopersicum* and *S. pennellii* alleles, *i*.*e*. multiple matches between different genes. Grey, partial matches between *S. lycopersicum* and *S. pennellii* alleles, *i*.*e*. CDS with conserved regions but otherwise dissimilar. Black, genes present in *S. lycopersicum* or *S. pennellii* only.

### Genomic analysis of ORG and identification of candidate genes

The resequenced dataset of tomato and wild relative accessions (Aflitos et al., 2014) was used to identify the polymorphisms of *S. pennellii* when aligned with *S. lycopersicum* (SL2.50) in the *ORG* region. We found 58 CDS within the *ORG* region in the *S. pennellii* genome and 65 CDS within *S. lycopersicum*, with considerable synteny (Figure 7d). Within the *ORG* region, an alignment of the *S. pennellii* genome sequence (Spenn-ch07:76,477,056-76,940,423) with *S. lycopersicum* (SL2.50ch07:65,444,176-65,865,655) showed that the two genomes were structurally similar (Supplemental Figure S8). We therefore investigated the similarities and differences in the coding sequences (CDS) between the two genomes with BLAST (Supplemental Table S3). We found a total of 6,009 polymorphisms, 5,093 of which were single-nucleotide polymorphisms (SNPs) and 916 were insertions-deletions (InDels). Additionally, there were 304 moderate effect missense variants affecting 58 genes (Supplemental Table S4) and 18 high effect polymorphisms (*e*.*g*. frameshift variants, stop gained) (Supplemental Table S5). There was one *S. pennellii* CDS without a corresponding match in *S. lycopersicum, i*.*e*. a new gene within the *ORG* region, namely Sopen07g031050 (hypothetical protein). Additionally, there were six presence-absence variants (PAVs) in *S. lycopersicum* without a corresponding match in *S. pennellii* (Supplemental Table S6), *i*.*e*. six genes lost in the *ORG* region, namely, a Yippee family protein (Solyc07g062900), a nucleolar GTP-binding protein 2 (Solyc07g063280), a Tir 2C resistance protein (Solyc07g063360) and three CDS annotated as ‘unknown protein’. The genes Sopen07g031090 and Sopen07g031100, both being putative Yippee family zinc-binding proteins, produced multiple significant matches with Solyc07g062880, Solyc07g062890 and Solyc07g062910. Additionally, Sopen07g031530 (beta glucosidase 46) and Sopen07g031540 (hypothetical protein) produced only partial matches with Solyc07g063370 (beta glucosidase) and Solyc07g063380 (unknown protein), respectively; indicating that the gene pairs share conserved regions but are otherwise dissimilar (Figure 7d).

## Discussion

The genetic basis of fruit gigantism has been extensively explored in tomato and a number of major genes controlling that trait have been identified (Nesbitt and Tanksley 2001; Causse et al. 2004; Muños et al. 2011; Chakrabarti et al. 2013; Mu et al. 2017). However, the genetic mechanisms behind isometric gigantism between vegetative and reproductive organs are unknown. Are they driven pleiotropically by genes for fruit gigantism that operate on the meristem simultaneously controlling vegetative and reproductive development, or are they the product of indirect selection on independent loci necessitated by the altered source-sink relationships between vegetative or reproductive organs? As a starting point to address this question, we set out to discover genetic determinants for changes in the size of vegetative organs in the tomato. We thus identified *ORGAN SIZE* (*ORG*), an introgression with reduced leaf size but which also showed smaller reproductive organs, namely flowers and fruits.

Instead of the conventional approach of QTL mapping, which sometimes is followed by fine-mapping and gene cloning, we revisited the alternative, forward genetics strategy, of wide cross followed by controlled introgression (Rick, 1969). We crossed *S. pennellii* to the tomato cv. Micro-Tom (MT) and conducted multiple rounds of crosses and backcrosses to the recurrent domesticated parental, selecting plants with smaller leaves in each generation. Our results, which identified the *ORG* locus, tie up previous, independent studies of the genetic control of leaf (Holtan and Hake, 2003; Chitwood *et al*., 2014) and fruit (Grandillo et al. 1999; van der Knaap and Tanksley 2003; Causse et al. 2004; Barrantes et al. 2016) size in tomato using QTL analysis. Hence, a survey of previous studies that identified putative QTLs for increased fruit weight during tomato domestication and breeding revel a chromosomal region overlapping *ORG* (Supplemental Figure 9). However, none of these studies reported alterations in vegetative development associated to fruit weight QTLs. This indicates that controlled introgression guided by phenotypic selection is a powerful tool that, unlike QTL mapping, allows the detection of genes (or closely linked genes) that control more than one trait simultaneously. Either QTL mapping, or its more up-to-date variant, genome-wide sequencing analysis (GWAS), are useful to detect multiple genes spread out in the genome controlling one trait, but on the other hand, are prone to miss pleiotropic or tightly linked genes controlling multiple traits, because generally only one phenotype is analysed at a time (Korte and Farlow, 2013).

Genotyping-by-sequencing showed *ORG* to harbour 1169 genes in approximately 11 Mb of *S. pennellii* genome. This represents 1.15% of the tomato genome, which is a good fit with the theoretically expected proportion of donor genome after six rounds of back-crossing (Stam and Zeven 1981). Although the segregation data indicate that *ORG* behaves as a Mendelian, semi-dominant gene, we cannot at this stage exclude the possibility that the IL harbours two or more genes controlling similar traits on chromosome 7. However, we showed that the common denominator for the reduced size of vegetative and reproductive organs in *ORG* is a reduction in the number of cells, possibly through alteration of cell division rate, as suggested by our gene expression analyses for *CYCB2;1, FW2*.*2* and *FW3*.*2*. This trait could be under pleiotropic control of a single gene. In fact, our analysis of the genes contained in the candidate region shows variation between *S. pennellii* and *S. lycopersicum* for genes predicted to be involved in the control of cell division, as well as regulatory genes that could control the size of organs (Supplemental Tables S4 and S5). An interval containing 19 putative domestication genes was also identified on chromosome 7 by Lin *et al*. (2014) by analyzing the genome sequence of 360 tomato accessions. All 19 genes are contained within the list of 58 candidates for the *ORG* region. This paves the way for the future identification and validation of, potentially, a single gene with a unique underlying variant (*e*.*g*. SNP, InDel, PAV) controlling organ size.

Increased organ size, or gigantism, is a recurrent domestication trait observed in many crops. Selection for increased size of edible parts led to allometric increases in reproductive organs. However, domesticated plants also tend to present gigantism in vegetative parts, *e*.*g*. larger leaves and thicker stems in *Phaseolus vulgaris* (Donald and Hamblin, 1983), larger leaves in eggplant (Page *et al*., 2019) and soybean (Kofsky *et al*., 2018). The tomato shows striking increases in fruit size (Tanksley 2004), but also leaf area, and stem thickness compared to its wild relatives (Milla and Matesanz, 2017). This isometric size change could lead to a better balance between photosynthetic sources and fruit sinks. When we altered the relative strength of the sinks by allowing only three, six or nine fruits to develop in either MT or *ORG* plants, we found an inverse correlation between fruit number and size in MT but not in *ORG*. In addition, the reduction in fruit size of MT has no penalty in its final yield. These results suggest two things. First, that the reduced size of *ORG* fruits is an intrinsic trait, possibly a developmental result of smaller ovaries, and not an indirect consequence of reduced leaf area (photosynthetic source). The second is that leaf area is not always directly limiting fruit (sink) size and/or yield. In agreement with this, both experimental and modelling work have shown that defoliation does not have a negative effect on crop yield, implying that source strength is not limiting (provided water and nutrient availability are sufficient and that photosynthesis is not light limited) (Heuvelink *et al*., 2005). An extreme situation is found in garden peas (*Pisum sativum*), where leaf area reduction has been a breeding goal to reduce interplant competition and increase yield (Cousin, 1997). Mutants of the ‘leafless’ and ‘semi-leafless’ type show 40% lower leaf area with up to 20% higher yield and better standing ability, which in turn facilitates mechanical harvesting (Checa *et al*., 2020). The increased popularity and growing market niche for ‘gourmet’ cherry tomatoes opens up the perspective of breeding varieties with smaller leaves to improve agronomic management (*e*.*g*. reduced fertilizer, water use) (Sarlikioti et al., 2011).

## Conclusions

Based on the analysis of natural genetic variation, we have described a potential genetic determinant for increased leaf size in cultivated tomato. Our results could unveil a novel link in the genetic control of isometric fruit and leaf gigantism in tomato. Further research to determine the molecular identity of the gene(s) underlying the *ORG* phenotype is underway. This knowledge would be a valuable addition in the repertoire of gene targets that can be manipulated with ideotype breeding (Donald, 1968; Zsögön et al. 2017) or *de novo* domestication platforms (Gasparini et al., 2021).

## Materials and methods

### Plant material

The wild relatives of tomato used in this work were *S. pennellii* (LA0716), *S. chilense* (LA1969), *S. peruvianum* (LA1537), *S. neorickii* (LA1322), *S. chmieslewskii* (LA1028), *S. habrochaites* f. *glabratum* (PI134417), *S. habrochaites* f. *hirsutum* (LA1777), *S. galapagense* (LA1401), *S. pimpinellifolium* (CNPH384), and *S. lycopersicum* var. *cerasiforme* (LA1320). Domesticated tomatoes of the cultivars Micro-Tom (MT) (LA3911), M82 (LA3475), Moneymaker (LA2706) and Santa Clara (Brazilian local cultivar) were also used. The *S. pennellii* chromosome 7 introgression lines (ILs) harboring alleles of *ORGAN SIZE* (*ORG*), *BRILLIANT COROLLA* (*Bco*) (Chetelat 1998) and *Rg7H* (Pinto et al. 2017) were obtained through repeated backcrossing between cultivated MT as a pollen receptor and *S. pennellii*, as described in Carvalho et al. (2011). Seeds of the tomato wild relatives were obtained from the UC Davis/C.M. Rick Tomato Genetics Resource Center, maintained by the Department of Plant Sciences, University of California, Davis, CA 95616. Seeds of MT were kindly donated by Prof. Avram Levy (Weizmann Institute of Science, Israel) in 1998 and kept as a true-to-type cultivar through self-pollination.

### Growth conditions

Plants were grown in a greenhouse at the Laboratory of Plant Developmental Genetics, ESALQ-USP, (543 m a.s.l., 22° 42’ 36” S; 47° 37’ 50” W), Piracicaba, SP, Brazil. Automatic irrigation took place four times a day. Growth conditions were: mean temperature of 28°C, 11.5 h/13 h (winter/summer) photoperiod, 250–350 µmol photons m^−2^ s^−1^ PAR irradiance, attained by a reflecting mesh (Aluminet, Polysack Indústrias Ltda, Leme, Brazil). Seeds were germinated in 350 mL pots with a 1:1 mixture of commercial potting mix Basaplant® (Base Agro, Artur Nogueira, SP, Brazil) and expanded vermiculite supplemented with 1 g L^−1^ 10:10:10 NPK and 4 g L^−1^ dolomite limestone (MgCO_3_ + CaCO_3_). Upon the appearance of the first true leaf, seedlings were transplanted to pots containing the soil mix described above, except for NPK supplementation, which was increased to 8 g L^−1^. In addition, MT and *OS* plants received a supplementary fertilization of 0.5g of NPK formulation 10:10:10 after flowering. Cultivated and wild tomato plants were supplemented with 2g of NPK formulation 10:10:10 per plant.

### Phenotypic characterization

We scanned all leaves of the MT and *ORG* plants 40 days after germination (dag) and determined the leaf area using ImageJ software (http://rsbweb.nih.gov/ij/).

For the characterization of floral whorls, we evaluated: length of petals and sepals; corolla area; and ovary weight, height and diameter. To measure ovary length and height we used a magnifying glass (Leica S8AP0, Wetzlar, Germany), coupled to a camera (Leica DFC295 Wetzlar, Germany). To determine ovary weight we determined the weight of 1.5 mL Eppendorf microtubes with 1 mL of distilled water, before and after collection of 10 ovaries. Ovary weight was then determined as the difference between initial and final tube weight. We also evaluated the leaf area and ovary weight of M82 plants and introgression lines (ILs) from chromosome 7, using the same methodology as for MT and *ORG*.

### MT and ORG productivity traits

We hand-pollinated MT and *ORG* plants with pollen from MT and *ORG* plants, because the *ORG* genotype displayed low fruit set. Various ovaries were pollinated, but after fruit set confirmation (five days after pollination), we performed selective fruit removal to allow only five fruits to set on each plant.

Productive performance of plants was assessed 90 days after germination. The following parameters were determined: mean weight per fruit; total soluble solids content in fruits (Brix); locule number and number of seeds per fruit; and weight of 10 seeds. Total soluble solids content of fruits was assessed using a digital refractometer (PR-101, Atago, Tokyo, Japan).

### Source-sink ratio in MT and ORG plants

To determine whether leaf area of *ORG* plants is a limiting factor for fruit development (since leaves and fruits are the major sources and sinks of photoassimilates, respectively), we manipulated plants creating three categories based on different source-to-sink ratios. Thus, we kept the same amount of source tissue (leaves) in all plants of each genotype and altered the sink strength by changing fruit number (either three, six or nine per plant, to produce high, medium or low source-to-sink ratios, respectively). We removed side branches to prevent them from acting as alternative sinks. The following parameters were then determined: total fruit weight per plant (yield); average fruit weight and whole-plant leaf area.

### Mapping and PCR amplification of DNA markers

We designed molecular markers to discover polymorphisms between tomato and *S. pennellii* in the region comprising the IL-7-2 and part of the IL 7-4 (Chitwood et al., 2014). The sequences and types of molecular makers are shown on Supplemental Table S1. Two further genotypes harbouring genome segments of *S. pennellii* for chromosome 7, *Brilliant corolla* (*Bco*) and *Regeneration 7h* (*Rg7H*), both in cv. MT, were characterized molecularly and phenotypically. Cross-referencing information from these genotypes and the ILs in the M82 background we constructed a map with the putative location of the *ORG* locus.

Genomic DNA extraction from young leaves was performed as described by Fulton et al. (1995). PCR was performed using the following program: a denaturation step at 95°C for 2 min, 35 cycles of 30 s at 95°C, 60s at 56°C, 90 s at 72°C, and a final cycle at 72°C for 7 min. When required, restriction enzyme analysis (Supplemental Table S1) was performed following the manufacturer’s recommendations (NEB, Bethesda, USA). The final PCR products were analyzed via 1.5% (m/v) agarose gel electrophoresis, stained with SYBR Gold (Invitrogen).

### Histological and microscopic analyses

Samples of MT and *ORG* ovaries/fruits at −8, −4, 0, 4 and 8 days, and fruit pericarps at 12 and 16 days (anthesis=0), were collected and fixed in Karnovsky solution (Karnovsky 1965), and vacuum-infiltrated for 15 min. The times referred to as −8 and −4 days correspond to 8 and 4 days before anthesis, respectively. We based these on the length of the closed flower buds (Faria 2014).

Samples were next dehydrated in an increasing ethanol series (10–100%), and infiltrated into synthetic resin, using a HistoResin embedding kit (Leica, www.leica-microsystems.com), according to the manufacturer’s instructions. The tissues were sliced using a rotary microtome (Leica RM 2045, Wetzlar, Germany), stained with toluidine blue 0.05% (Sakai 1973), and photographed in a microscope (Leica DMLB, Heidelberg, Germany), coupled to a Leica DFC310 camera (Wetzlar, Germany). Histological analysis of ovaries was performed in the central region of the outer pericarp of the fruits, and the area and number of cells were determined using ImageJ software (http://rsbweb.nih.gov/ij/). This histological analysis also was performed in the mature leaves of these genotypes adopting the procedures described above. The area and number of cells in the adaxial leaf epidermis of the MT and *ORG* genotypes was also evaluated using the leaf dental resin imprinting technique (Weyers and Johansen 1985).

### Quantitative real-time reverse transcription PCR

Total RNA was extracted from ovaries/fruits at −8, −4, 0, 4 and 8 days, and fruit pericarps at 12 and 16 days (anthesis = 0), using Trizol reagent (Invitrogen), as indicated by the manufacturer, and treated with RQ1 RNAse-Free DNAse (Promega). Fruit pericarps were carefully collected from the central region of the outer pericarp of the fruits, at 12 and 16 days. After DNase treatment, a single-strand cDNA was synthesized from total RNA (1µg) by reverse-transcription, using RevertAid RT Reverse Transcription Kit (Thermo Fisher Scientific).

Gene expression analyses were performed on a Rotor-Gene Q real-time PCR cycler (Qiagen), using Kapa Sybr Fast qPCR Master Mix (Kapa Biosystems) and specific primers for *CYCB2;1* (Solyc02g082820), *FW2*.*2* (Solyc02g090730), *FW3*.*2* (Solyc03g114940) and *EXP5* (Solyc02g088100) genes. The reactions were amplified for 2 min at 95 °C, followed by 40 cycles of 95 °C for 15 s and 60 °C for 30 s. The threshold cycle (C_T_) was determined. Melting curve analysis was performed with each primer set to confirm the presence of only a single peak before the gene expression analyses. Two technical replicates were analyzed for each of three or four biological samples. The relative transcript accumulation was normalized to an *ACTIN* (Solyc04g011500) gene. The fold changes for each gene were calculated using the equation 2^-ΔΔCT^ (Livak and Schmittgen 2001). Primer sequences used to qRT-PCR are shown in Supplemental Table S1.

### In silico *analysis of probable* ORG *region*

The genomes of *S. lycopersicum* cv. Heinz 1706, SL2.50 (https://solgenomics.net/) and *S. pennellii* LA716 (Bolger et al., 2014b) were aligned and plotted with Mummer v4.0.0 (Marcais et al., 2018). Variants of *S. pennellii* LA716 versus SL2.50 within the *OS* region were obtained through the Wageningen resequencing project (Aflitos et al., 2014). The coding sequences of the genes within the region were obtained from Solanaceae Genomics Network (https://solgenomics.net/) and similarities between Heinz 1706 and LA716 were tested with BLAST v2.10.0 (Camacho et al., 2009). The Circos plot was created with Circos v0.69.9 (Krzywinski et al., 2009) on Windows 10. The synteny plot was created with the genoPlotR package (Guy et al., 2011) within R (Team, 2017).

### Genotyping by sequencing (GBS)

DNA was extracted from young leaf samples (∼10 mm length) that were freeze-dried (CoolSafe™ 55-9; Scanvac, Lynge, Denmark) overnight. Leaf samples were powdered in a Star-Beater (VWR, Lutterworth, UK) at 30 Hz for 30s in 2 mL microcentrifuge tubes containing two 5 mm acid-rinsed soda-glass balls. DNA was extracted from ∼50 mg samples with an E.Z.N.A® Plant DNA Kit (VWR, Lutterworth, UK). DNA fragment size was assessed on a 1% agarose gel in Tris/Borate/EDTA to confirm that all samples had the majority of DNA fragments >10 kilobases.

The GBS library was prepared using the restriction enzyme *Msl*I and sequenced on an Illumina NextSeq 500 V2 by LGC Genomics (Berlin, Germany). The 150 base-pair paired-end reads were aligned to the *Solanum lycopersicum* Heinz 1706 reference genome (SL2.50) with BWA v0.7.15 (Li and Durbin, 2009). The SAM files were processed with Samtools Fixmate v1.3.1 (Li et al., 2009). InDels were realigned with GATK’s IndelRealigner v3.8-0 (Mckenna et al., 2010; Depristo et al., 2011) before variant calling with Samtools Mpileup v1.3.1 and Bcftools Call v1.3 (Li, 2011).

The raw VCF files of the GBS sample, a 40× resequenced Micro-Tom (Cranfield University, unpublished data) and the resequencing of *S. pennellii* LA716 (Aflitos et al., 2014), were combined into an index using Tersect (Kurowski and Mohareb, 2020). Tersect was used to determine which variants were shared between the *ORG* IL and *S. pennellii* LA716, excluding the variants shared with Micro-Tom. The variants output from Tersect were then filtered as follows: all variants with a quality score less than 20, a mapping quality score below 40 and a raw read depth either below 10 and above 200 were removed. In addition, heterozygous variants were removed. The variant density of the filtered variants over a 10 kb window (5 kb sliding) were plotted across all 12 chromosomes with ggplot2 (Wickham, 2016) within R.

### Statistical analysis

Statistical analysis was performed using SAS software (SAS Institute Inc., Cary, NC, USA). The variables data were submitted to analysis of variance (ANOVA) and the means compared by the Student’s t*-* or Tukey’s test. When the data did not meet the assumptions of ANOVA, we performed to non-parametric analysis, using Wilcoxon rank sum or Dunn’s test to compare the means.

## Supplemental Data

Supplemental Figure S1. Leaf size increases during tomato domestication and improvement.

Supplemental Figure S2. Heterozygous *ORG* plants (*ORG*/+) show an intermediate leaf area compared to MT and *ORG* plants.

Supplemental Figure S3. Smaller leaf size in *ORG* is caused by reduced cell division. Supplemental Figure S4. *ORG* reduces organ size in all floral whorls

Supplemental Figure 5. Fruit traits are altered in *ORG* plants.

Supplemental Figure 6. GBS defines the span of the introgression in the *Brilliant corolla (Bco)* introgression line.

Supplemental Figure 7. Characterization of *S. pennellii* introgression lines (IL) in chromosome *7*.

Supplemental Figure 8. Alignment plot of the *S. pennellii* and *S. lycopersicum* genomes within the *ORG* region.

Supplemental Figure 9. Colocalization of ORG and previously mapped fruit size QTLs. Supplemental Table S1. Oligonucleotide sequences used for genotyping and quantitative PCR analyses in this work.

Supplemental Table S2. Fruit weight of MT and ORG plants.

Supplemental Table S3. Similarities and discrepancies between coding sequences of S. pennellii v. S. lycopersicum candidate genes.

Supplemental Table S4. Polymorphisms with a moderate effect on gene function for S. pennellii v S. lycopersicum within the ORG region.

Supplemental Table S5. Polymorphisms with a high effect on gene function for S. pennellii v S. lycopersicum within the ORG region.

Supplemental Table S6. Coding sequences of S. lycopersicum not producing a match on the S. pennellii genome assembly.

## Acknowledgements

AZ was partly funded by a grant (RED□00053□16) from the Foundation for Research Assistance of the Minas Gerais State (FAPEMIG, Brazil). AT, FM, ZK were partially funded by BBSRC BB/S007970/1; AT and AZ were partially funded by the Royal Society’s Newton Mobility Grant NMG\R2\170027 and the Global Challenges Research Fund (GCRF, UK Research and Innovation). LEPP was supported by grant 306518/2018-0 from CNPq (Brazil) and 2018/050003-1 from FAPESP (Brazil). MHV gratefully acknowledges a PhD scholarship from FAPESP (16/05566-0). We thank Diego S. Reartes for valuable technical assistance.

