## Supplemental Figures for "The *ORGAN SIZE* (*ORG*) locus contributes to isometric gigantism in domesticated tomato"

**Supplemental Data**

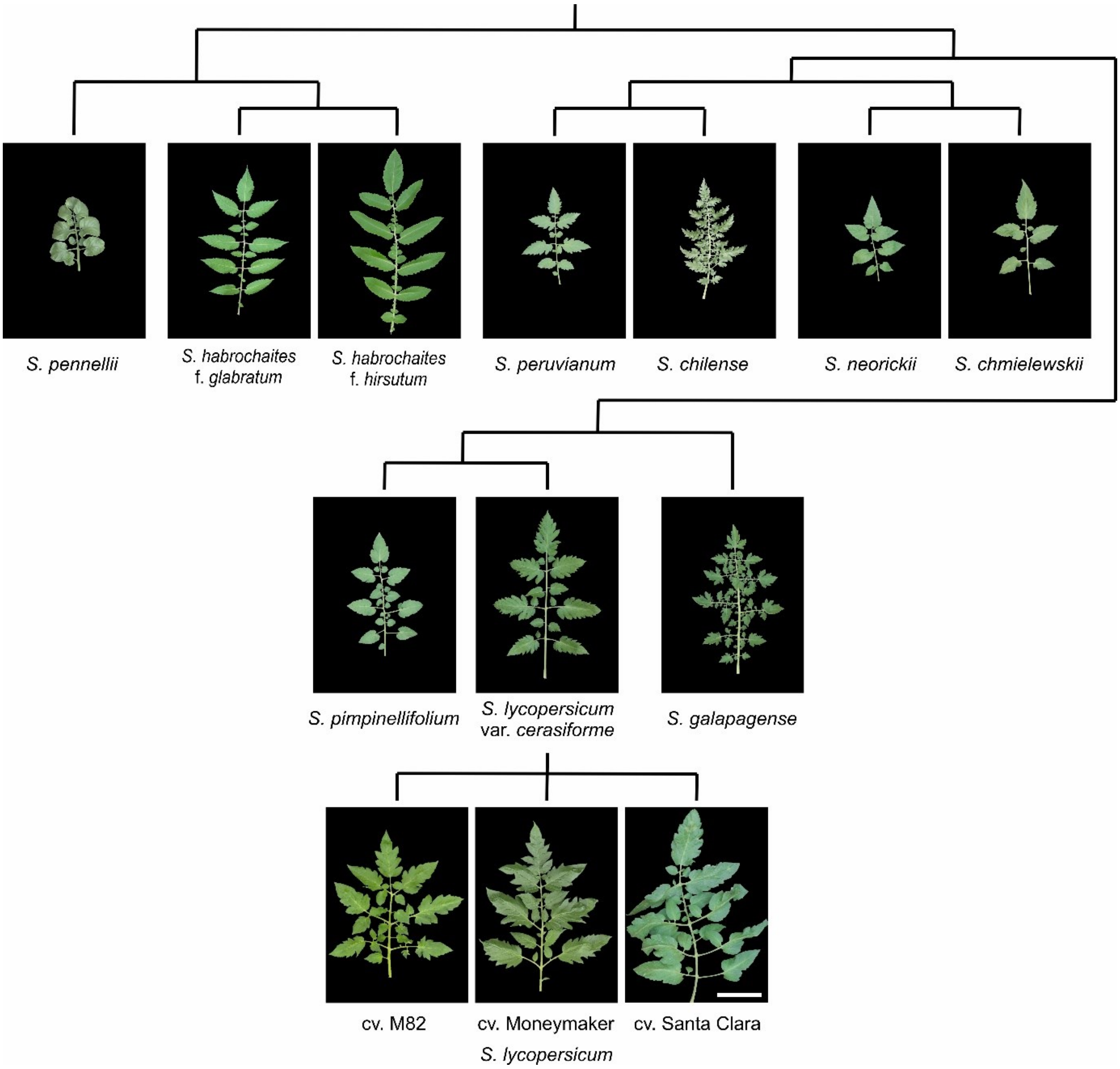


**Supplemental Figure S1. Leaf size increases during tomato domestication and improvement.** Representative accessions of *Solanum pennelli* (LA0716); *S. habrochaites* f. *glabratum* (PI134417); *S. habrochaites* f. *hirsutum* (LA1777); *S. peruvianum* (LA1537); *S. chilense* (LA1969); *S. neorickii* (LA1322); *S. chmieslewskii* (LA1028); *S. pimpinellifolium* (CNPH384); *S. lycopersicum* var. *cerasiforme* (LA1320); *S. galapagense* (LA1401); *S. lycopersicum* cv. M82 (LA3475); *S. lycopersicum* cv. Ailsa Craig (LA2838A); *S. lycopersicum* cv. Moneymaker (LA2706); *S. lycopersicum* cv. Santa Clara (local cultivar). Scale bar = 10 cm. Phylogenetic hypotheses from Aflitos et al. (2014).

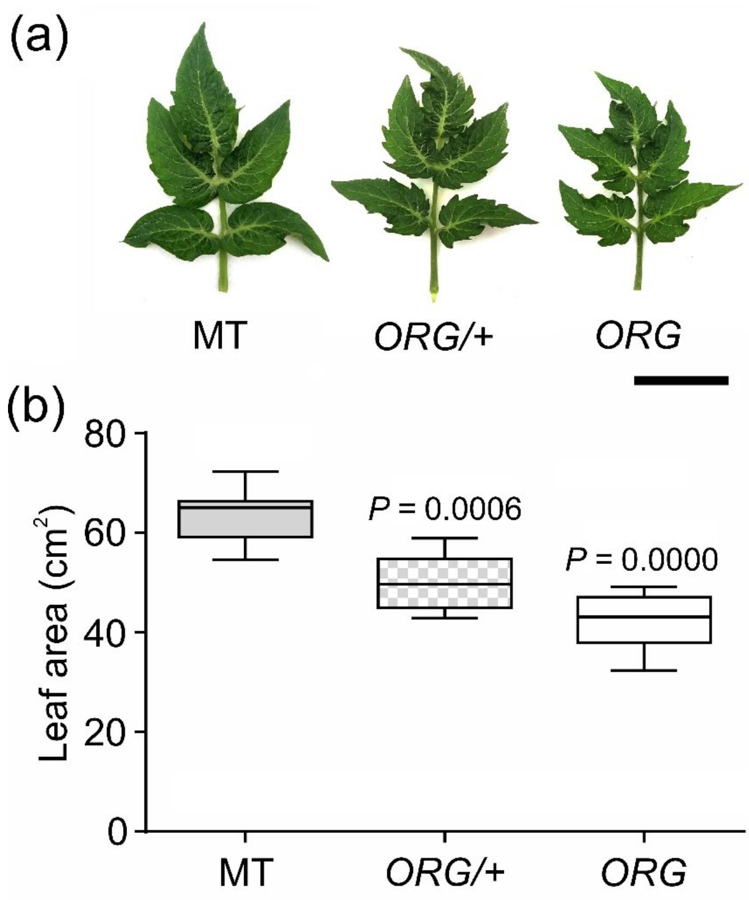


**Supplemental Figure S2. Heterozygous *ORG* plants (*ORG*/+) show an intermediate leaf area compared to MT and *ORG* plants. (a**) Representative leaf from MT, *ORG*/+, and *ORG* plants. Scale bar=5 cm. **(b)** Area of representative leaves of the MT, *ORG/+*, and *ORG* genotypes, 50 days after germination (*n* at the bottom of the graphic represents the leaf number on each evaluation). Statistical significance was tested by Tukey's test ($p$<0.001).

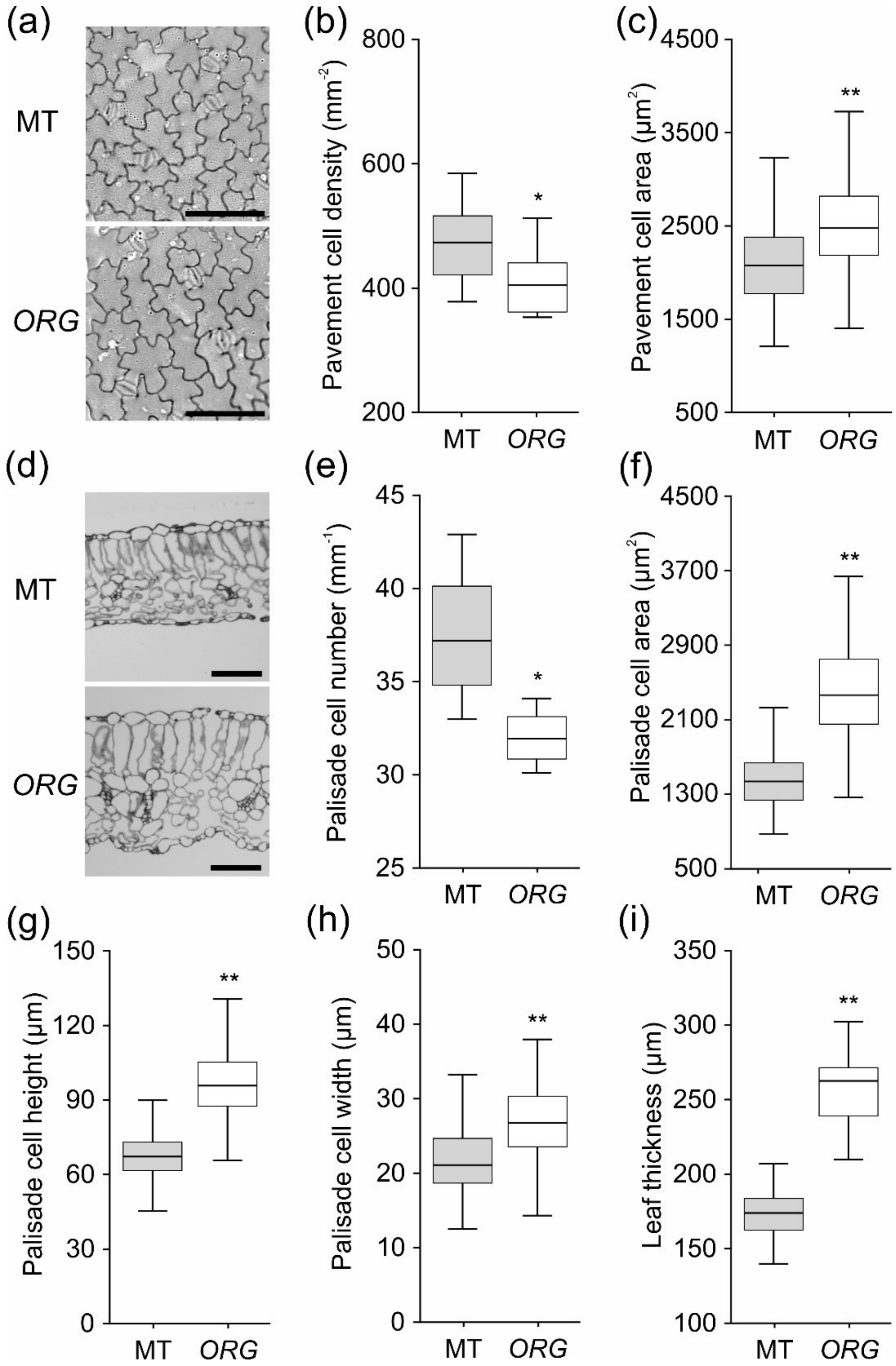


**Supplemental Figure S3. Smaller leaf size in *ORG* is caused by reduced cell division. (a)** Representative imprints of adaxial side from MT (top) and *ORG* (bottom) leaves. Scale bar=100µm. **(b)** Number of pavement cells per $mm^2$ in adaxial side of MT (gray box) and *ORG* (white box) leaves (n=14 sections). **(c)** Cell area in adaxial side of MT and *ORG* leaves (n=320 cells). **(d)** Representative cross-sections of MT (top) and *ORG* (bottom) leaves. Scale bar=100µm. **(e-h)** Characterization of palisade parenchyma. Palisade cell number (e), area (f), height (g) and width (h) from MT and *ORG* leaves (n=5). **(i)** Leaf thickness (n=5). Data are means±s.e.m. Statistical significance was tested by Student's *t*-test (*$p$<0.05, **$p$<0.01).

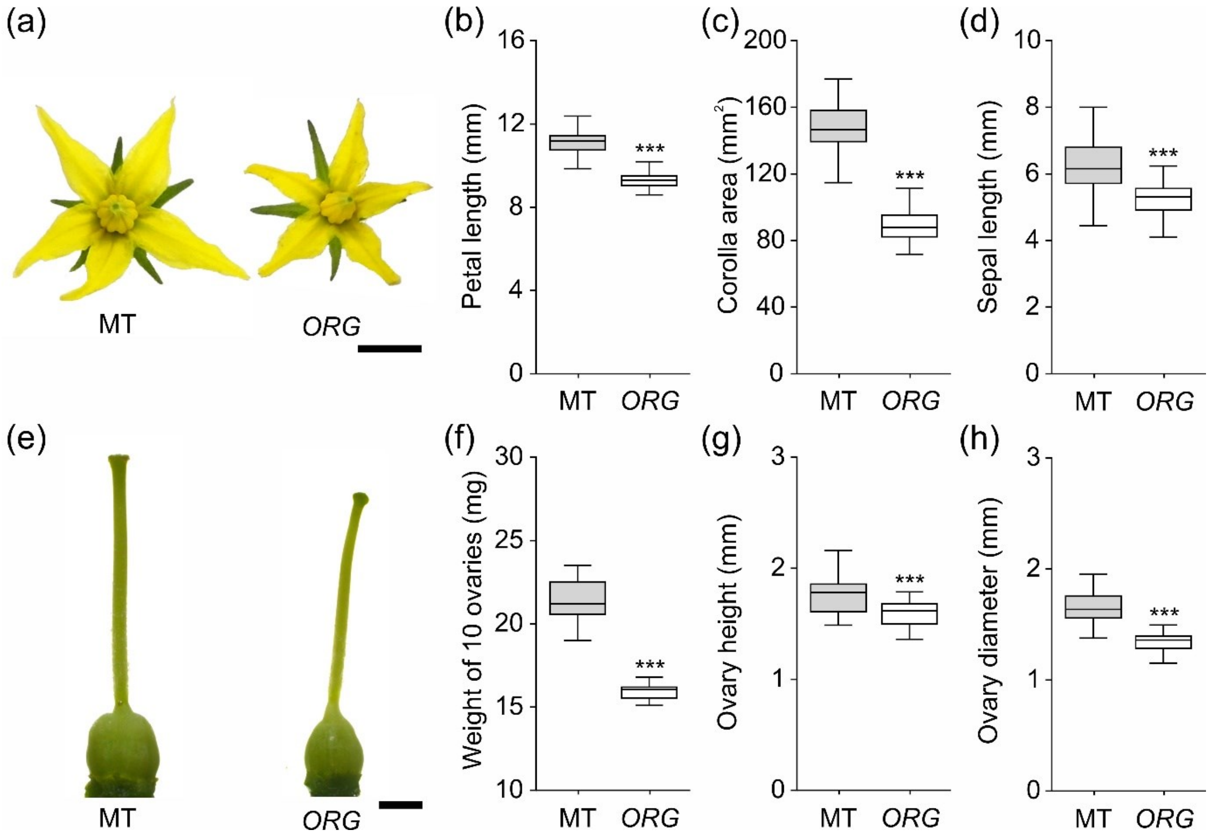


**Supplemental Figure S4.** ***ORG*** **reduces organ size in all floral whorls (a)** Representative MT (left) and *ORG* (right) flower at anthesis. Scale bar=0.5 cm. **(b-d)** Petal length (b), corolla area and sepal length (d) of MT (gray box) and *ORG* (white box) flowers (n=45 flowers). **(e)** Representative MT (left) and *ORG* (right) ovary at anthesis. Scale bar=1 mm. **(f)** Fresh weight of 10 ovaries at anthesis (n=13). **(g** and **h)** Ovary height (g) and diameter (h) at anthesis (n=25 flowers). Data are mean±s.e.m. *** indicate significant differences by Wilcoxon rank sum test ($p$<0.001).

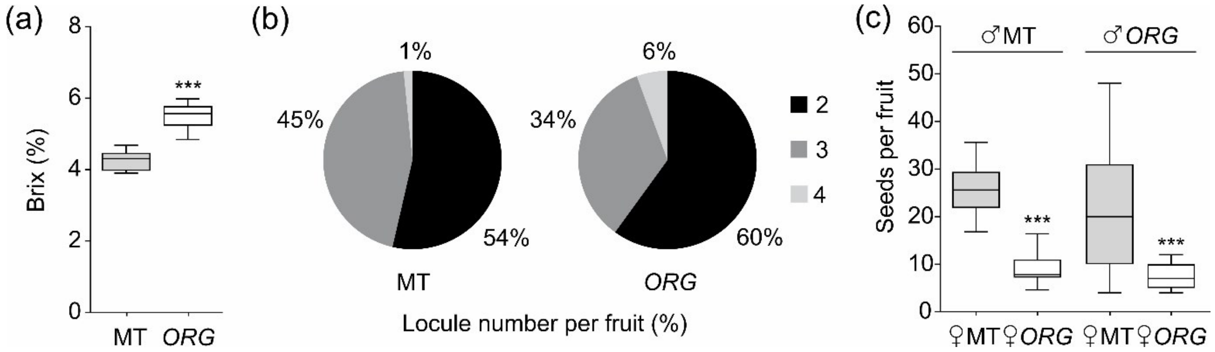


**Supplemental Figure 5. Fruit traits are altered in *ORG* plants. (a)** The average total soluble solids content in MT and *ORG* fruits (Brix) (n = 10 plants with 5 fruits on each). **(b)** Frequency of locule number per fruit in MT and *ORG* fruits (n=125 fruits). (c) Seeds per fruit of MT and *ORG* pollinated with MT (n=11 plants) and *ORG* (n= 15 fruits) pollen. Data are mean±s.e.m. Statistical significance was tested by Student's t test (***p<0.001).

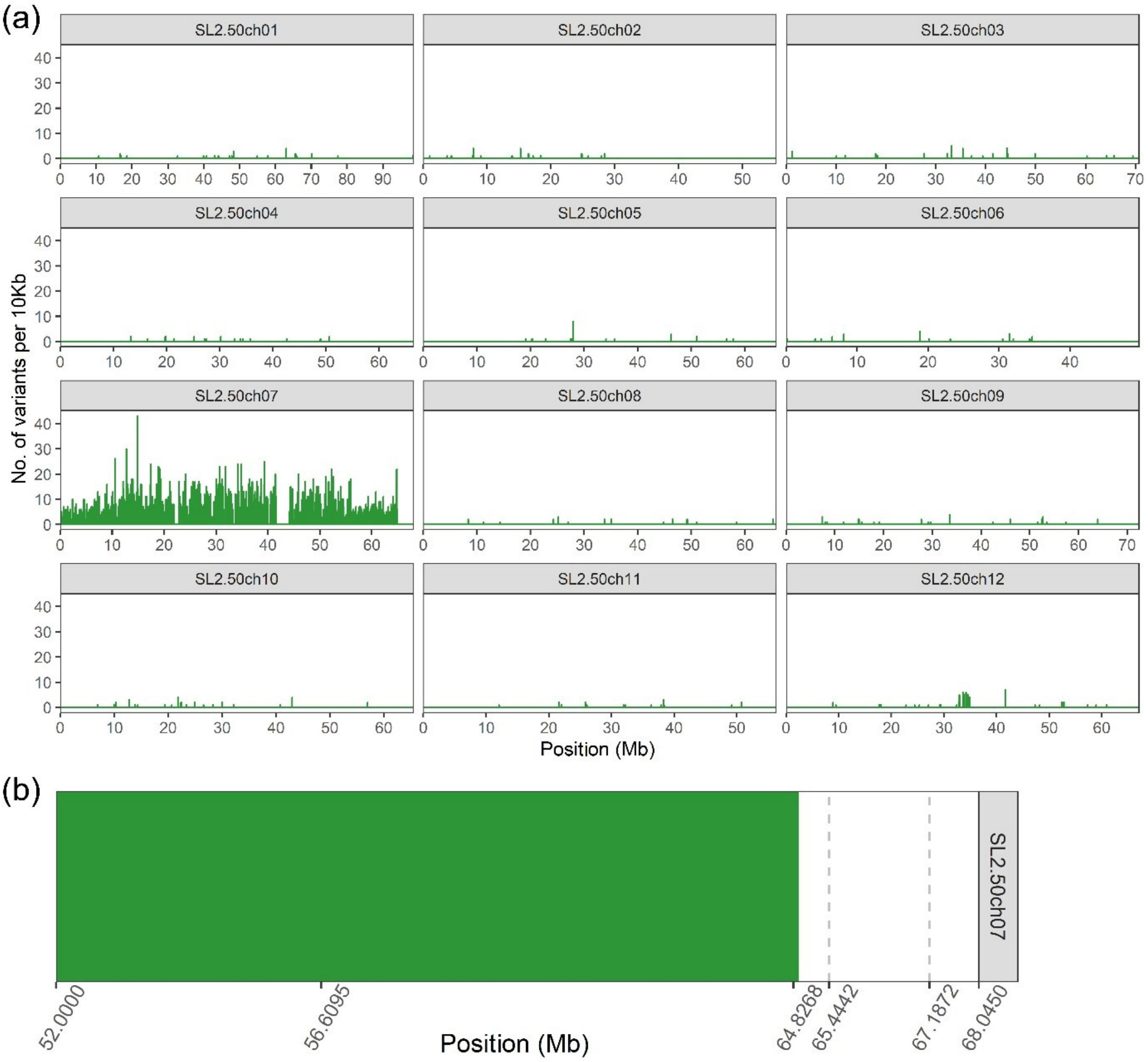


**Supplemental Figure 6. GBS defines the span of the introgression in the *Brilliant corolla (Bco)* introgression line. (a)** Genome-wide density of unique variants shared between *Bco* IL and *S. pennellii* LA716 in the genetic background of tomato cv Micro-Tom. **(b)** Close up view of the introgression on chromosome 7.

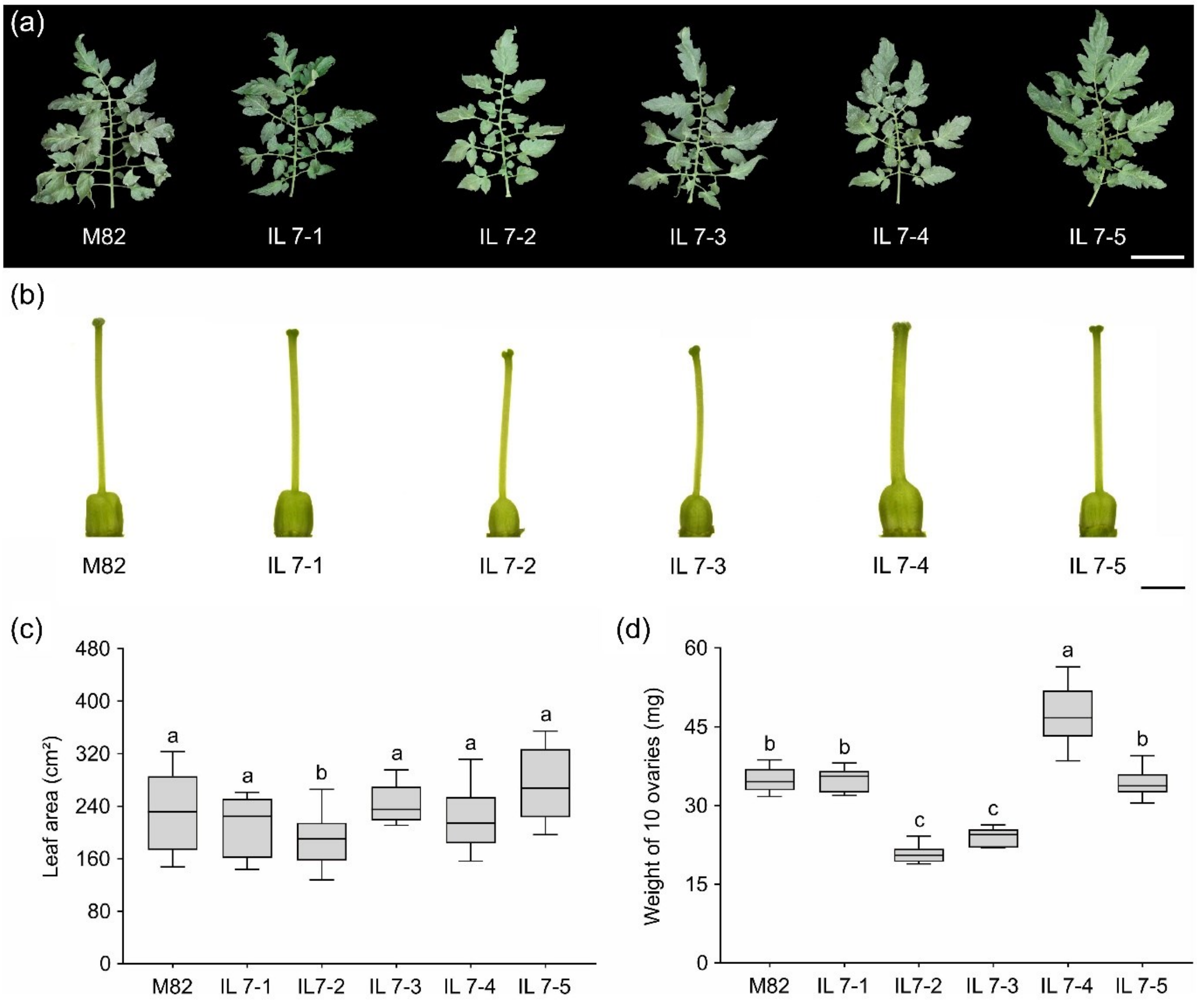


**Supplemental Figure 7. Characterization of *S. pennellii* introgression lines (IL) in chromosome *7*. (a-b)** Representative fully-expanded fifth leaf (a) and ovary at anthesis (b) of M82, IL7-1, IL7-2, IL7-3, IL7-4 and IL7-5 plants. Scale bars =10 cm and 2 mm, respectively. **(c-d)** Leaf area (c) and weight of 10 ovaries (d) of M82 and chr 7 ILs (n=14). Statistical significance was tested by ANOVA followed by Tukey's test. Different letters indicate significant differences ($p$<0.05) between genotypes.

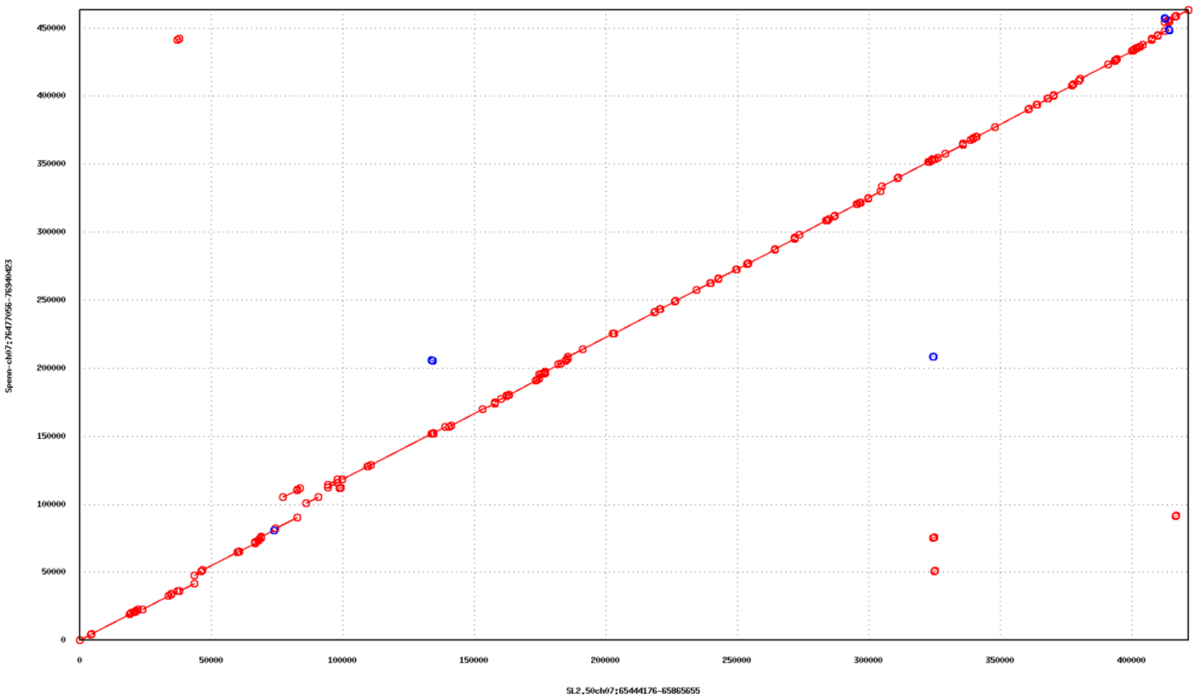


**Supplemental Figure 8. Alignment plot of the *S. pennellii* and *S. lycopersicum* genomes within the *ORG* region.** The *S. pennellii* genome sequence was aligned to *S. lycopersicum* with the nucmer script in Mummer v4.0 and the alignment was subsequently potted with the mummerplot script using GNU plot. Aligned sequences are depicted by a line between two dots. Alignments in red depict *S. pennellii* sequences aligning on the same (*i.e.* forward) strand of *S. lycopersicum*. Alignments in blue depict *S. pennellii* sequences aligning on the opposite (*i.e.* reverse) strand of *S. lycopersicum*.

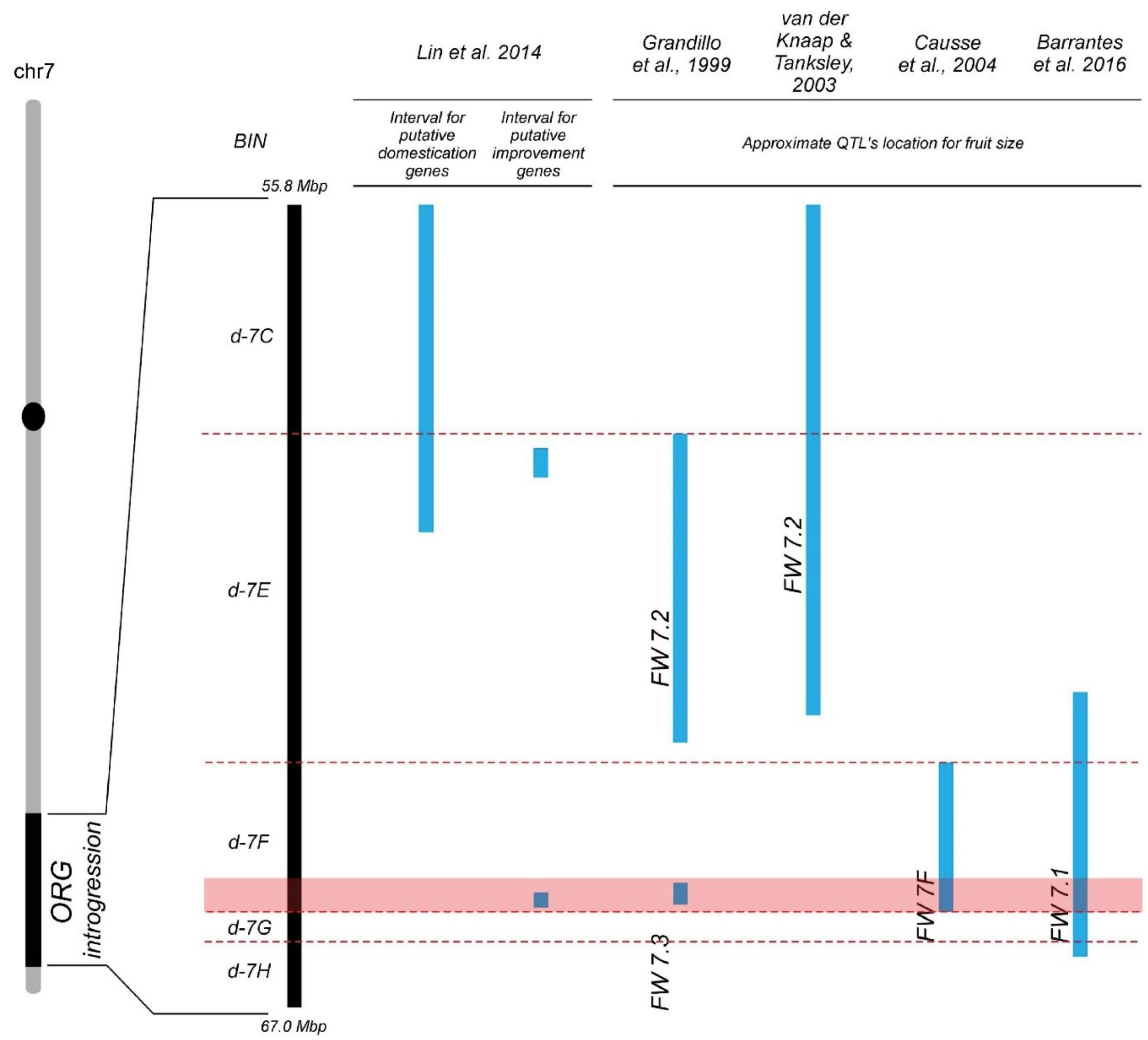


**Supplemental Figure 9. Colocalization of ORG and previously mapped fruit size QTLs.** The *ORG* region coincided with previously mapped QTLs (Quantitative Trait Loci) affecting fruit weight on chromosome *7*. See text for details. Bars indicate the approximate QTL location.
