## Supplemental Tables for "The *ORGAN SIZE* (*ORG*) locus contributes to isometric gigantism in domesticated tomato"

**Supplemental Table S1.** Oligonucleotide sequences used for genotyping and quantitative PCR analyses in this work.

| Locus ID[1] | Forward | Reverse | Utilization | Enzyme | Reference |
|---|---|---|---|---|---|
| Solyc02g082820_*CYCB2;1* | GAAGGCAGCAACAGGGAAACTAA | CCCTCTTCTCCTCTATTCTTGTGTC | qPCR | | This work |
| Solyc02g090730_*FW2.2* | GTTTTGATGACCCTGCTAACTG | CAAAGAGCACAAGGTTCACAG | qPCR | | This work |
| Solyc03g114940_*FW3.2* | CAGGTGGTCTTGCTTGGTC | CCAACTGAAAACGCCATTAACG | qPCR | | This work |
| Solyc02g088100_EXP5 | ACCATCGCCTGTAGTGACCTTAAAG | AAGGGTTCAAGAACTCAATGGCAAC | qPCR | | Chusreeaeom *et al* . (2014)[3] |
| Solyc04g011500_*ACTIN* | GGTCCCTCTATTGTCCACAG | TGCATCTCTGGTCCAGTAGGA | qPCR | | Silva *et al* . (2018)[4] |
| Solyc07g009130 | TCTTACTCTCCGTCATTAGCCT | ATGTCCTGTTTCCTTCGCAA | Genotyping | XhoI | This work |
| Solyc07g017650 | GTAGGTTACCAAGTAGTAGAGGG | AGAGCTACATACCTTATCCAACA | Genotyping | EcoRV | This work |
| Solyc07g020960 | CATTATCTTATTTCCACCCACC | TGAACCTCATCTAACCCTACC | Genotyping | HincII | This work |
| Solyc07g026680 | GTGGATATGGGAAGGAAGT | CGAGGAATATGATGTATGGT | Genotyping | RsaI | This work |
| Solyc07g039570 | TGGCAATACTTTCAACCTTCC | CAATGTCTGAGGACGAGCA | Genotyping | PsiI | This work |
| Solyc07g041190 | GTGGACTTGACCTGGAGATG | AAGAGAGAAGGATTGGGGACT | Genotyping | RsaI | This work |
| Solyc07g041980 | GTGGACTTGACCTGGAGATG | AAGAGAGAAGGATTGGGGACT | Genotyping | RsaI | This work |
| Solyc07g042560 | CTGTTAATACCGAATAATCTTGTAAT | GGGACTAAACAAAACAATGAATGG | Genotyping | BstNI | This work |
| Solyc07g043450 | CCATTACATCTTCAAAGCCATCTC | GGCATAGTGACTACAATACATATCG | Genotyping | PagI | This work |
| Solyc07g045180 | GCTTCCAGGGTGTCAATG | ATGTCTAAGTCTGCCAACC | Genotyping | HpyCH4V | This work |
| Solyc07g047950 | GCCATACCTAGACGGGATTT | TATTTCCAGGCACAAGAAGC | Genotyping | DpnII | This work |
| Solyc07g051830[2] | CAATGAGGACAGACTGATAGC | CACCTGATCTTTGACAACGAG | Genotyping | | This work |
| Solyc07g052970[2] | TGGATGTCAATGGTGAACAAG | GCCCCACACATGAACAATG | Genotyping | | This work |
| Solyc07g054090 | TTTGTCAACCTTCATCAGCCTT | AGAGAAAAAGCAGGTAAGGAGAG | Genotyping | Hpy188I | This work |
| Solyc07g055020 | CCGCCATCTTGACCCATTTATA | CTTTCTTCCGCAATCCACTCAT | Genotyping | HincII | This work |
| Solyc07g055400 | GCAACCCAAAATCCAAATCAAC | ATCTCCCTGTAGACACCATG | Genotyping | Hpy188I | This work |
| Solyc07g056060 | GCAAGAAGGCAATCAACTAC | CACTTGTAAAGACTTCCTGC | Genotyping | PsiI | This work |
| Solyc07g056580 | CCACGACTGTTATGCTTTATG | GGGTATCAGTTCACTTTCTTG | Genotyping | TaqI | This work |
| Solyc07g062600 | AAGCCATTCACCTTATGTCAG | CCACCAGGAGAACTTGATTCA | Genotyping | EcoRI | This work |
| Solyc07g062750 | GGGTTTGTTGTGTCCTATGTG | TGGGTTGGTGTGATGCTTC | Genotyping | ApaLI | This work |
| Solyc07g062930 | CCTTTCTCCTACCTCCCAGAT | GAATTGTGGTCCTGAGTGTAGA | Genotyping | MspI and HaeIII | This work |
| Solyc07g063010[2] | GTGATGATTTGAAGGAGGATTGA | TTATCAGTCACCTCTCTGCCAG | Genotyping | | This work |
| Solyc07g063080 | GTGTCCAGGGAGAAGGTA | GACGTTCTCGACTGTTGA | Genotyping | RsaI | This work |
| Solyc07g063130 | GCAGAAAAGACAAGGTCAATCC | CAAACTCCCTCAACTCACCCTT | Genotyping | HpyCH4V | This work |
| Solyc07g063300 | CCATTGTAGGTGTTACGGTGC | CAGAACCAATAGATGCGAGTAGT | Genotyping | DraI | This work |
| Solyc07g063390 | GTCTTACTGGCTTAAAACTTGG | CTTGGGAACTCGATCTAATGTA | Genotyping | EcoRV | This work |
| Solyc07g063480 | GCCTTTATTTCTATCCTCTCC | GCTAGACCTTGTTCAGATCTC | Genotyping | PsiI | This work |
| Solyc07g063590 | GGTGCTATGAAGGTGAAAGGA | GTTGTGAAGAACAGATGGCTC | Genotyping | HinfI | This work |
| Solyc07g064080 | TCTAAGTTGAGAGTTATGGATGATGT | TGGAACGAATCATGAAGGAA | Genotyping | EcoRV | This work |
| Solyc07g064300 | AGCATCTTAATATAGGGGGCTTA | CTCTTATTCGGGCAAGCAAG | Genotyping | DraI | This work |
| Solyc07g064550 | TCGTCTCCTTGTTGATCCTGT | TCCAAACACCATCCAGCATA | Genotyping | PstI | This work |
| Solyc07g064830 | AATGGGCTCCAGGTTCAAAT | TTGTTGACCACCCATTGAAG | Genotyping | EcoRV | This work |
| Solyc07g065050 | TGTGGGCATTTTGTTGACTG | CAATAGCAAAACAAAAGGCATC | Genotyping | PstI | This work |
| Solyc07g065270 | GTCGGGTAACAGTTCGTGCT | CCCTGAGCAATCTGGAAATC | Genotyping | MspI | This work |
| Solyc07g065440 | CTCAGAAGAAGAACGAGCACAG | TGGTGCATCCTGTGTAACATC | Genotyping | HinfI | This work |
| Solyc07g066630 | TGATAGCTTAAATGTTGTGGGAAG | GCAACCAAGCAACTAACCAAA | Genotyping | HindII | This work |

[1] Locus according to the Sol Genomics Network database (http://solgenomics.net/).

[2] SCAR (sequence characterized amplified regions) markers

[3] Silva *et al* . 2018. *SELF-PRUNING* Acts Synergistically with *DIAGEOTROPICA* to guide auxin responses and proper growth form. *Plant Physiology* , 176, 2904–2916.

[4] Chusreeaeom *et al* . 2014. Regulatory change in cell division activity and genetic mapping of a tomato (*Solanum lycopersicum* L.) elongated-fruit mutant. *Plant Biotechnology* , 31: 149–158.

**Supplemental Table S2**. Fruit weight (g)

| Pollinated with MT pollen | | |
|---|---|---|
| Plant | MT | ORG |
| 1 | 7.86 | 6.14 |
| 1 | 7.54 | 5.96 |
| 1 | 7.46 | 4.91 |
| 1 | 6.51 | 4.7 |
| 1 | 4.9 | 3.84 |
| 2 | 7.16 | 4.7 |
| 2 | 7.16 | 4.43 |
| 2 | 6.55 | 4.4 |
| 2 | 5.96 | 4 |
| 2 | 5.71 | 3.97 |
| 3 | 11.63 | 7.58 |
| 3 | 8.21 | 6.24 |
| 3 | 7.94 | 6.24 |
| 3 | 7.8 | 6 |
| 3 | 4.46 | 4.5 |
| 4 | 9.82 | 5.52 |
| 4 | 8.4 | 5.46 |
| 4 | 8.1 | 5.1 |
| 4 | 7.8 | 4.91 |
| 4 | 7.3 | 4.6 |
| 5 | 8.14 | 6.1 |
| 5 | 7.85 | 5.33 |
| 5 | 7.67 | 4.8 |
| 5 | 5.2 | 4.8 |
| 5 | 3.22 | 3.58 |
| 6 | 10.2 | 6.32 |
| 6 | 9.58 | 5.93 |
| 6 | 8.74 | 5.48 |
| 6 | 6.14 | 4.99 |
| 6 | 5.62 | 4 |
| 7 | 12.3 | 6.71 |
| 7 | 11.2 | 5.35 |
| 7 | 10.85 | 4.99 |
| 7 | 4.69 | 4.58 |
| 7 | 4.25 | 3.27 |
| 8 | 9.36 | 6.8 |
| 8 | 6.6 | 6.12 |
| 8 | 6.42 | 5 |
| 8 | 5.6 | 4.98 |
| 8 | 5.1 | 3.36 |
| 9 | 9.13 | 6.55 |
| 9 | 8.74 | 6.54 |
| 9 | 8.41 | 6.29 |
| 9 | 7.46 | 5.4 |
| 9 | 5.07 | 5.3 |
| 10 | 10.2 | 6.13 |
| 10 | 9.46 | 5.69 |
| 10 | 9.38 | 5.25 |
| 10 | 8.2 | 5.13 |
| 10 | 5.78 | 4.7 |
| **Mean** | **7.58** | **5.25** |
| SD | 2.0255 | 0.9497 |
| SEM | 0.2864 | 0.1343 |
| *P* -value | 0.0001 | |

| Pollinated with ORG pollen | | |
|---|---|---|
| Fruit | MT | ORG |
| **1** | 5.83 | 3.5 |
| **2** | 6.5 | 5.38 |
| **3** | 7.85 | 4.53 |
| **4** | 5 | 3.04 |
| **5** | 6.87 | 3.46 |
| **6** | 4.5 | 5.08 |
| **7** | 6.71 | 3.73 |
| **8** | 7.8 | 3.47 |
| **9** | 6.35 | 4.98 |
| **10** | 7.18 | 3.79 |
| **11** | 6.17 | 3.36 |
| **12** | 8.93 | 4.99 |
| **13** | 8.87 | 5.66 |
| **14** | 6.22 | 4.71 |
| **Mean** | **6.77** | **4.26** |
| SD | 1.288 | 0.8707 |
| SEM | 0.3442 | 0.2327 |
| *P* -value | 0.0001 | |

| Legend | |
|---|---|
| Colour | Description |
| | Matches with high similarity between SPENN and SLYC, however, SPENN genes contain SNPs |
| | Matches with high similarity between SPENN and SLYC, however, SPENN genes contain InDels |
| | Matches with high similarity between SPENN and SLYC, however, SPENN genes contain HIGH effect polymorphisms |
| | Matches with high similarity between SPENN and SLYC, however, SPENN genes contain HIGH effect polymorphisms. Genes are annotated as HYPOTHETICAL/UNKNOWN. |
| | SPENN genes without a corresponding match in SLYC |
| | SPENN genes with only a partial match to SLYC genes |
| | SPENN genes matching with multiple SLYC genes |

**Supplemental Table S3. SIMILARITIES AND DISCREPANCIES BETWEEN THE CODING SEQUENCES OF S.PENNELLII VS S.LYCOPERSICUM (SL2.50) AS DETERMINED BY BLAST.** The coding sequences (CDS) of S.pennellii (Bolger et al. 2014) were queried against the CDS of S.lycopersicum SL2.50 using BLAST+ and subsequently grouped according to the legend above. The variant information was obtained from the S.pennellii resequencing dataset (Aflitos et al. 2014).

| Number | S.pennellii query | Matching S.lyc gene (SL2.50) | Spenn annotation | SL2.50 Annotation | Comments | Variants vs SL2.50: Variant type | Variants vs SL2.50: Description |
|---|---|---|---|---|---|---|---|
| 1 | Query= Sopen07g030980.1<br>Length=954 | >Solyc07g062750.2.1<br>Length=954<br>Score = 1668 bits (903), Expect = 0.0<br>Identities = 937/954 (98%), Gaps = 0/954 (0%) | Note=Polynucleotidyl transferase, ribonuclease H-like superfamily protein \| FUNCTIONS IN: exonuclease activity \| LOCATED IN: intracellular, chloroplast \| EXPRESSED IN: 14 plant structures \| EXPRESSED DURING: 9 growth stages \| CONTAINS InterPro DOMAIN/s: Exonuclease, RNase T/DNA polymerase III | Note=ERI1 exoribonuclease 3 (AHRD V1 **-- ERI3_BOVIN)%3B contains Interpro domain(s) IPR006055 Exonuclease ;Ontology_term=GO:0005515;from_BOGAS=1;interpro2go_term=GO:0005622;length=3928;nb_exon=6;sifter_term=GO:0005515 | | missense_variant | Qty: 4 |
| 2 | Query= Sopen07g030990.1<br>Length=5496 | >Solyc07g062760.1.1<br>Length=3315<br>Score = 5925 bits (3208), Expect = 0.0<br>Identities = 3268/3298 (99%), Gaps = 0/3298 (0%) | Note=P-loop containing nucleoside triphosphate hydrolases superfamily protein \| FUNCTIONS IN: nucleoside-triphosphatase activity, ATPase activity, nucleotide binding, ATP binding \| EXPRESSED IN: 22 plant structures \| EXPRESSED DURING: 13 growth stages \| CONTAINS InterPro DOMAIN/s: ATPase, AAA-type, core , ATPase, AAA+ type, core , ATPase, AAA-type, conserved site \| BEST Arabidopsis thaliana protein match is: cell division cycle protein 48-related / CDC48-related | Note=26S protease regulatory subunit 4 homolog (AHRD V1 *--- PRS4_ORYSJ)%3B contains Interpro domain(s) IPR003959 ATPase%2C AAA-type%2C core ;Ontology_term=GO:0005524,GO:0017111;eugene_evidence_code=10F0H1E1IEG;from_BOGAS=1;interpro2go_term=GO:0005524,GO:0017111;length=4458;nb_exon=4 | In SL4.0 760 and 770 have been fused into one gene | frameshift_variant<br>missense_variant | Phe1101fs<br>Qty: 12 |
| | | >Solyc07g062770.2.1<br>Length=2211<br>Score = 3917 bits (2121), Expect = 0.0<br>Identities = 2173/2199 (99%), Gaps = 0/2199 (0%) | | Note=Unknown Protein (AHRD V1);from_BOGAS=1;length=2962;nb_exon=4 | | missense_variant | Qty: 12 |
| 3 | Query= Sopen07g031000.1<br>Length=774 | >Solyc07g062780.2.1<br>Length=768<br>Score = 1362 bits (737), Expect = 0.0<br>Identities = 755/764 (99%), Gaps = 0/764 (0%) | Note=Encodes a SUMO E3 ligase that regulates endocycle onset and meristem maintenance. \| HIGH PLOIDY2 (HPY2) | Note=E3 SUMO-protein ligase NSE2 (AHRD V1 **** C3KJP6_ANOFI)%3B contains Interpro domain(s) IPR004181 Zinc finger%2C MIZ-type ;Ontology_term=GO:0019789;from_BOGAS=1;interpro2go_term=GO:0008270;length=3441;nb_exon=7;sifter_term=GO:0019789 | | missense_variant | Qty: 3 |
| 4 | Query= Sopen07g031010.1<br>Length=525 | >Solyc07g062790.1.1<br>Length=525<br>Score = 942 bits (510), Expect = 0.0<br>Identities = 520/525 (99%), Gaps = 0/525 (0%) | Note=Encodes a homeodomain leucine zipper class I (HD-Zip I) protein. \| homeobox protein 52 (HB52) \| FUNCTIONS IN: sequence-specific DNA binding transcription factor activity \| INVOLVED IN: response to blue light, regulation of transcription, DNA-dependent, response to absence of light \| LOCATED IN: nucleus \| EXPRESSED IN: root, flower \| EXPRESSED DURING: seedling growth \| CONTAINS InterPro DOMAIN/s: Homeobox , Homeobox, conserved site , Homeodomain-like , Homeodomain-related \| BEST Arabidopsis thaliana protein match is: homeobox 3 | Note=Homeobox leucine zipper protein (AHRD V1 ***- Q9XH73_PRUAR)%3B contains Interpro domain(s) IPR001356 Homeobox ;Ontology_term=GO:0005634,GO:0006355,GO:0045449;eugene_evidence_code=10F0H1E0IEG;from_BOGAS=1;interpro2go_term=GO:0005634,GO:0006355,GO:0045449;length=525;nb_exon=1 | | missense_variant | Qty: 2 |
| 5 | Query= Sopen07g031020.1<br>Length=891 | >Solyc07g062810.2.1<br>Length=1050<br>Score = 1624 bits (879), Expect = 0.0<br>Identities = 887/891 (99%), Gaps = 0/891 (0%) | Note=D-aminoacid aminotransferase-like PLP-dependent enzymes superfamily protein \| FUNCTIONS IN: 4-amino-4-deoxychorismate lyase activity, catalytic activity \| INVOLVED IN: tetrahydrofolate biosynthetic process \| LOCATED IN: chloroplast \| EXPRESSED IN: 24 plant structures \| EXPRESSED DURING: 15 growth stages \| CONTAINS InterPro DOMAIN/s: Aminotransferase, class IV \| BEST Arabidopsis thaliana protein match is: D-aminoacid aminotransferase-like PLP-dependent enzymes superfamily protein | Note=Branched-chain amino acid aminotransferase-like (AHRD V1 **** Q5NAM3_ORYSJ)%3B contains Interpro domain(s) IPR001544 Aminotransferase%2C class IV ;Ontology_term=GO:0008696;from_BOGAS=1;interpro2go_term=GO:0008152;length=3568;nb_exon=6;sifter_term=GO:0008696 | | splice_acceptor_variant&intron_variant<br>missense_variant | <br>Qty: 4 |
| 6 | Query= Sopen07g031030.1<br>Length=1854 | >Solyc07g062820.1.1<br>Length=1854<br>Score = 3230 bits (1749), Expect = 0.0<br>Identities = 1819/1854 (98%), Gaps = 0/1854 (0%) | Note=Pentatricopeptide repeat (PPR) superfamily protein \| CONTAINS InterPro DOMAIN/s: Pentatricopeptide repeat \| BEST Arabidopsis thaliana protein match is: Tetratricopeptide repeat (TPR)-like superfamily protein | Note=Pentatricopeptide repeat protein PPR1106-17 (AHRD V1 ***- B6SWY6_MAIZE)%3B contains Interpro domain(s) IPR002885 Pentatricopeptide repeat ;Ontology_term=GO:0008116;eugene_evidence_code=10F0H1E1IEG;from_BOGAS=1;length=1854;nb_exon=1;sifter_term=GO:0008116 | | missense_variant | Qty: 6 |
| 7 | Query= Sopen07g031040.1<br>Length=1059 | >Solyc07g062840.2.1<br>Length=1059<br>Score = 1851 bits (1002), Expect = 0.0<br>Identities = 1040/1059 (98%), Gaps = 0/1059 (0%) | Note=Transcriptional activator of the NAC gene family, with CUC1 redundantly required for embryonic apical meristem formation, cotyledon separation and expression of STM. Proper timing of CUC2 expression is required to maintain the phyllotactic pattern initiated in the meristem. CUC2 expression in leaf sinus region is required for serration and the extent of serration is modulated by mir164A mediated repression of CUC2. \| CUP-SHAPED COTYLEDON 2 (CUC2) \| CONTAINS InterPro DOMAIN/s: No apical meristem (NAM) protein \| BEST Arabidopsis thaliana protein match is: NAC (No Apical Meristem) domain transcriptional regulator superfamily protein | Note=NAC domain protein IPR003441 (AHRD V1 ***- B9N1D6_POPTR)%3B contains Interpro domain(s) IPR003441 No apical meristem (NAM) protein ;Ontology_term=GO:0003700;from_BOGAS=1;interpro2go_term=GO:0045449;length=2145;nb_exon=3;sifter_term=GO:0003700 | | missense_variant | Qty: 5 |
| | | >Solyc07g063410.2.1<br>Length=1071<br>Score = 108 bits (58), Expect = 5e-25<br>Identities = 198/265 (75%), Gaps = 12/265 (5%) | | Note=NAC domain protein IPR003441 (AHRD V1 ***- B9HZY3_POPTR)%3B contains Interpro domain(s) IPR003441 No apical meristem (NAM) protein ;Ontology_term=GO:0016563,GO:0003700;from_BOGAS=1;interpro2go_term=GO:0045449;length=1578;nb_exon=3;sifter_term=GO:0016563,GO:0003700 | | | |
| 8 | Query= Sopen07g031050.1<br>Length=411 | | Note=hypothetical protein | | | | |
| 9 | Query= Sopen07g031060.1<br>Length=1059 | >Solyc07g062850.2.1<br>Length=1086<br>Score = 1860 bits (1007), Expect = 0.0<br>Identities = 1042/1059 (98%), Gaps = 1/1059 (0%) | Note=ARM repeat superfamily protein \| FUNCTIONS IN: binding \| INVOLVED IN: biological_process unknown \| EXPRESSED IN: 22 plant structures \| EXPRESSED DURING: 11 growth stages \| CONTAINS InterPro DOMAIN/s: Armadillo-type fold , 26S proteasome non-ATPase regulatory subunit 5 ., 26S proteasome non-ATPase regulatory subunit 5 | Note=Genomic DNA chromosome 3 BAC clone F4B12 (AHRD V1 ***- Q9LIL7_ARATH)%3B contains Interpro domain(s) IPR016024 Armadillo-type fold ;Ontology_term=GO:0005488;from_BOGAS=1;interpro2go_term=GO:0005488;length=7917;nb_exon=11 | | frameshift_variant<br>stop_lost<br>missense_variant | Lys349fs<br>Ter362Glnext*?<br>Qty: 8 |
| | Query= Sopen07g031060.2<br>Length=1572 | >Solyc07g062850.2.1<br>Length=1086<br>Score = 1753 bits (949), Expect = 0.0<br>Identities = 973/985 (99%), Gaps = 0/985 (0%) | | Note=Genomic DNA chromosome 3 BAC clone F4B12 (AHRD V1 ***- Q9LIL7_ARATH)%3B contains Interpro domain(s) IPR016024 Armadillo-type fold ;Ontology_term=GO:0005488;from_BOGAS=1;interpro2go_term=GO:0005488;length=7917;nb_exon=11 | In Sl4.0 850 and 860 have been fused into one gene | frameshift_variant<br>stop_lost<br>missense_variant | Lys349fs<br>Ter362Glnext*?<br>Qty: 8 |
| | | >Solyc07g062860.2.1<br>Length=492<br>Score = 857 bits (464), Expect = 0.0<br>Identities = 482/491 (98%), Gaps = 0/491 (0%) | | Note=Genomic DNA chromosome 3 BAC clone F4B12 (AHRD V1 ***- Q9LIL7_ARATH);from_BOGAS=1;length=4534;nb_exon=6 | | missense_variant | Qty: 1 |
| | Query= Sopen07g031060.3<br>Length=822 | >Solyc07g062850.2.1<br>Length=1086<br>Score = 1417 bits (767), Expect = 0.0<br>Identities = 800/816 (98%), Gaps = 1/816 (0%) | | Note=Genomic DNA chromosome 3 BAC clone F4B12 (AHRD V1 ***- Q9LIL7_ARATH)%3B contains Interpro domain(s) IPR016024 Armadillo-type fold ;Ontology_term=GO:0005488;from_BOGAS=1;interpro2go_term=GO:0005488;length=7917;nb_exon=11 | | frameshift_variant<br>stop_lost<br>missense_variant | Lys349fs<br>Ter362Glnext*?<br>Qty: 8 |
| | Query= Sopen07g031060.4<br>Length=276 | >Solyc07g062860.2.1<br>Length=492<br>Score = 466 bits (252), Expect = 2e-133<br>Identities = 268/276 (97%), Gaps = 0/276 (0%) | | Note=Genomic DNA chromosome 3 BAC clone F4B12 (AHRD V1 ***- Q9LIL7_ARATH);from_BOGAS=1;length=4534;nb_exon=6 | | missense_variant | Qty: 1 |
| 10 | Query= Sopen07g031070.1<br>Length=555 | >Solyc07g062870.2.1<br>Length=555<br>Score = 1014 bits (549), Expect = 0.0<br>Identities = 553/555 (99%), Gaps = 0/555 (0%) | Note=chloroplast 30S ribosomal protein S20, putative \| FUNCTIONS IN: structural constituent of ribosome, RNA binding \| INVOLVED IN: translation, ribosome biogenesis \| LOCATED IN: ribosome, chloroplast stroma, chloroplast, membrane \| EXPRESSED IN: 22 plant structures \| EXPRESSED DURING: 13 growth stages \| CONTAINS InterPro DOMAIN/s: Ribosomal protein S20 | Note=30S ribosomal protein S20 (AHRD V1 *-*- RS20_THEEB)%3B contains Interpro domain(s) IPR002583 Ribosomal protein S20 ;Ontology_term=GO:0005840;from_BOGAS=1;interpro2go_term=GO:0005840;length=3283;nb_exon=3 | | | |
| 11 | Query= Sopen07g031080.1<br>Length=390 | >Solyc07g062880.2.1<br>Length=390<br>Score = 721 bits (390), Expect = 0.0<br>Identities = 390/390 (100%), Gaps = 0/390 (0%) | Note=Yippee family putative zinc-binding protein \| CONTAINS InterPro DOMAIN/s: Yippee-like protein \| BEST Arabidopsis thaliana protein match is: Yippee family putative zinc-binding protein | Note=Yippee zinc-binding-like protein (AHRD V1 **** C1FDK9_9CHLO)%3B contains Interpro domain(s) IPR004910 Yippee-like protein ;Ontology_term=GO:0005515,GO:0008270;from_BOGAS=1;length=4567;nb_exon=5;sifter_term=GO:0005515,GO:0008270 | | | |
| 12 | Query= Sopen07g031090.1<br>Length=396 | >Solyc07g062890.2.1<br>Length=372<br>Score = 549 bits (297), Expect = 2e-158<br>Identities = 302/304 (99%), Gaps = 1/304 (0%) | Note=Yippee family putative zinc-binding protein \| CONTAINS InterPro DOMAIN/s: Yippee-like protein \| BEST Arabidopsis thaliana protein match is: Yippee family putative zinc-binding protein | Note=Yippee zinc-binding-like protein (AHRD V1 *-** C1FDK9_9CHLO)%3B contains Interpro domain(s) IPR004910 Yippee-like protein ;Ontology_term=GO:0005515,GO:0008270;from_BOGAS=1;length=5871;nb_exon=6;sifter_term=GO:0005515,GO:0008270 | | missense_variant | Qty: 2 |
| | | >Solyc07g062910.2.1<br>Length=663<br>Score = 388 bits (210), Expect = 5e-110<br>Identities = 306/352 (87%), Gaps = 8/352 (2%)<br>Score = 130 bits (70), Expect = 4e-32<br>Identities = 88/97 (91%), Gaps = 0/97 (0%)<br>Score = 128 bits (69), Expect = 1e-31<br>Identities = 87/96 (91%), Gaps = 0/96 (0%) | | Note=Yippee family protein (AHRD V1 *-*- D7MTM3_ARALY)%3B contains Interpro domain(s) IPR004910 Yippee-like protein ;from_BOGAS=1;length=11892;nb_exon=8 | | frameshift_variant<br>missense_variant | Tyr215fs<br>Qty: 5 |
| 13 | Query= Sopen07g031100.1<br>Length=393 | >Solyc07g062910.2.1<br>Length=663<br>Score = 501 bits (271), Expect = 6e-144<br>Identities = 324/349 (93%), Gaps = 5/349 (1%)<br>Score = 130 bits (70), Expect = 4e-32<br>Identities = 88/97 (91%), Gaps = 0/97 (0%)<br>Score = 128 bits (69), Expect = 1e-31<br>Identities = 87/96 (91%), Gaps = 0/96 (0%) | Note=Yippee family putative zinc-binding protein \| CONTAINS InterPro DOMAIN/s: Yippee-like protein \| BEST Arabidopsis thaliana protein match is: Yippee family putative zinc-binding protein | Note=Yippee family protein (AHRD V1 *-*- D7MTM3_ARALY)%3B contains Interpro domain(s) IPR004910 Yippee-like protein ;from_BOGAS=1;length=11892;nb_exon=8 | | frameshift_variant<br>missense_variant | Tyr215fs<br>Qty: 5 |
| | | >Solyc07g062890.2.1<br>Length=372<br>Score = 381 bits (206), Expect = 9e-108<br>Identities = 273/305 (90%), Gaps = 6/305 (2%) | | Note=Yippee zinc-binding-like protein (AHRD V1 *-** C1FDK9_9CHLO)%3B contains Interpro domain(s) IPR004910 Yippee-like protein ;Ontology_term=GO:0005515,GO:0008270;from_BOGAS=1;length=5871;nb_exon=6;sifter_term=GO:0005515,GO:0008270 | | missense_variant | Qty: 2 |
| | | >Solyc07g062880.2.1<br>Length=390<br>Score = 379 bits (205), Expect = 3e-107<br>Identities = 323/381 (85%), Gaps = 3/381 (1%) | | Note=Yippee zinc-binding-like protein (AHRD V1 **** C1FDK9_9CHLO)%3B contains Interpro domain(s) IPR004910 Yippee-like protein ;Ontology_term=GO:0005515,GO:0008270;from_BOGAS=1;length=4567;nb_exon=5;sifter_term=GO:0005515,GO:0008270 | | | |
| | Query= Sopen07g031100.2<br>Length=873 | >Solyc07g062890.2.1<br>Length=372<br>Score = 538 bits (291), Expect = 1e-154<br>Identities = 298/301 (99%), Gaps = 1/301 (0%)<br>Score = 379 bits (205), Expect = 7e-107<br>Identities = 274/307 (89%), Gaps = 6/307 (2%) | Note=Yippee family putative zinc-binding protein \| CONTAINS InterPro DOMAIN/s: Yippee-like protein \| BEST Arabidopsis thaliana protein match is: Yippee family putative zinc-binding protein | Note=Yippee zinc-binding-like protein (AHRD V1 *-** C1FDK9_9CHLO)%3B contains Interpro domain(s) IPR004910 Yippee-like protein ;Ontology_term=GO:0005515,GO:0008270;from_BOGAS=1;length=5871;nb_exon=6;sifter_term=GO:0005515,GO:0008270 | | missense_variant | Qty: 2 |
| | | >Solyc07g062910.2.1<br>Length=663<br>Score = 459 bits (248), Expect = 9e-131<br>Identities = 267/276 (97%), Gaps = 1/276 (0%)<br>Score = 383 bits (207), Expect = 6e-108<br>Identities = 305/352 (87%), Gaps = 8/352 (2%)<br>Score = 130 bits (70), Expect = 8e-32<br>Identities = 88/97 (91%), Gaps = 0/97 (0%)<br>Score = 128 bits (69), Expect = 3e-31<br>Identities = 87/96 (91%), Gaps = 0/96 (0%) | | Note=Yippee family protein (AHRD V1 *-*- D7MTM3_ARALY)%3B contains Interpro domain(s) IPR004910 Yippee-like protein ;from_BOGAS=1;length=11892;nb_exon=8 | | frameshift_variant<br>missense_variant | Tyr215fs<br>Qty: 5 |
| | | >Solyc07g062880.2.1<br>Length=390<br>Score = 313 bits (169), Expect = 7e-87<br>Identities = 251/291 (86%), Gaps = 3/291 (1%) | | Note=Yippee zinc-binding-like protein (AHRD V1 **** C1FDK9_9CHLO)%3B contains Interpro domain(s) IPR004910 Yippee-like protein ;Ontology_term=GO:0005515,GO:0008270;from_BOGAS=1;length=4567;nb_exon=5;sifter_term=GO:0005515,GO:0008270 | | | |
| | | >Solyc07g062900.2.1<br>Length=192<br>Score = 124 bits (67), Expect = 4e-30<br>Identities = 69/70 (99%), Gaps = 0/70 (0%) | | Note=Yippee family protein (AHRD V1 ***- D7MTM3_ARALY)%3B contains Interpro domain(s) IPR004910 Yippee-like protein ;from_BOGAS=1;length=4569;nb_exon=3 | | | |
| 14 | Query= Sopen07g031110.1<br>Length=1818 | >Solyc07g062920.2.1<br>Length=1818<br>Score = 3197 bits (1731), Expect = 0.0<br>Identities = 1789/1818 (98%), Gaps = 0/1818 (0%) | Note=hypothetical protein | Note=Genomic DNA chromosome 5 TAC clone K19P17 (AHRD V1 *-*- Q9FN33_ARATH);from_BOGAS=1;length=4642;nb_exon=3 | | missense_variant | Qty: 16 |
| 15 | Query= Sopen07g031120.1<br>Length=1170 | >Solyc07g062930.2.1<br>Length=1161<br>Score = 2013 bits (1090), Expect = 0.0<br>Identities = 1145/1170 (98%), Gaps = 9/1170 (1%) | Note=ribosomal protein L11 methyltransferase-related \| FUNCTIONS IN: protein methyltransferase activity \| INVOLVED IN: protein amino acid methylation \| LOCATED IN: cytoplasm \| CONTAINS InterPro DOMAIN/s: Ribosomal protein L11 methyltransferase , Ribosomal L11 methyltransferase, PrmA | Note=Ribosomal protein L11 methyltransferase-like protein (AHRD V1 **** Q9FN34_ARATH)%3B contains Interpro domain(s) IPR010456 Ribosomal L11 methyltransferase ;Ontology_term=GO:0008757;from_BOGAS=1;interpro2go_term=GO:0005737;length=4977;nb_exon=7;sifter_term=GO:0008757 | | missense_variant | Qty: 8 |
| 16 | Query= Sopen07g031130.1<br>Length=2118 | >Solyc07g062940.2.1<br>Length=2151<br>Score = 3749 bits (2030), Expect = 0.0<br>Identities = 2090/2118 (99%), Gaps = 8/2118 (0%) | Note=Protein kinase superfamily protein \| FUNCTIONS IN: protein serine/threonine kinase activity, protein kinase activity, kinase activity, ATP binding \| INVOLVED IN: protein amino acid phosphorylation \| LOCATED IN: cytoplasm, spindle \| EXPRESSED IN: 21 plant structures \| EXPRESSED DURING: 11 growth stages \| CONTAINS InterPro DOMAIN/s: Protein kinase, ATP binding site , Protein kinase, catalytic domain , Serine/threonine-protein kinase domain , Serine/threonine-protein kinase-like domain , Protein kinase-like domain \| BEST Arabidopsis thaliana protein match is: Protein kinase superfamily protein | Note=Serine/threonine kinase 25 (AHRD V1 *-*- Q6V9V7_CHICK)%3B contains Interpro domain(s) IPR002290 Serine/threonine protein kinase ;Ontology_term=GO:0006468;from_BOGAS=1;interpro2go_term=GO:0006468;length=15792;nb_exon=22 | | missense_variant | Qty: 7 |
| 17 | Query= Sopen07g031140.1<br>Length=1263 | >Solyc07g062950.2.1<br>Length=1263<br>Score = 2300 bits (1245), Expect = 0.0<br>Identities = 1257/1263 (99%), Gaps = 0/1263 (0%) | Note=Serine/threonine-protein kinase WNK (With No Lysine)-related \| BEST Arabidopsis thaliana protein match is: Serine/threonine-protein kinase WNK (With No Lysine)-related | Note=BHLH transcription factor (AHRD V1 *-*- Q9XIN0_ARATH);from_BOGAS=1;length=3036;nb_exon=7 | | missense_variant | Qty: 1 |
| | Query= Sopen07g031140.2<br>Length=948 | >Solyc07g062950.2.1<br>Length=1263<br>Score = 1729 bits (936), Expect = 0.0<br>Identities = 944/948 (99%), Gaps = 0/948 (0%) | | | | | |
| 18 | Query= Sopen07g031150.1<br>Length=645 | >Solyc07g062960.1.1<br>Length=669<br>Score = 1127 bits (610), Expect = 0.0<br>Identities = 632/643 (98%), Gaps = 0/643 (0%) | Note=hypothetical protein | Note=Unknown Protein (AHRD V1);eugene_evidence_code=10F0H0E0IEG;from_BOGAS=1;length=669;nb_exon=1 | | missense_variant | Qty: 3 |
| 19 | Query= Sopen07g031160.1<br>Length=849 | >Solyc07g062970.2.1<br>Length=849<br>Score = 1524 bits (825), Expect = 0.0<br>Identities = 841/849 (99%), Gaps = 0/849 (0%) | Note=Protein phosphatase 2C family protein \| FUNCTIONS IN: protein serine/threonine phosphatase activity, catalytic activity \| EXPRESSED IN: 23 plant structures \| EXPRESSED DURING: 14 growth stages \| CONTAINS InterPro DOMAIN/s: Protein phosphatase 2C-related , Protein phosphatase 2C , Protein phosphatase 2C, N-terminal \| BEST Arabidopsis thaliana protein match is: Protein phosphatase 2C family protein | Note=Serine/threonine phosphatase family protein (AHRD V1 **** C1E0J1_9CHLO)%3B contains Interpro domain(s) IPR015655 Protein phosphatase 2C ;Ontology_term=GO:0004722;from_BOGAS=1;interpro2go_term=GO:0003824;length=3117;nb_exon=5;sifter_term=GO:0004722 | | missense_variant | Qty: 1 |
| 20 | Query= Sopen07g031170.1<br>Length=396 | >Solyc07g062980.2.1<br>Length=399<br>Score = 706 bits (382), Expect = 0.0<br>Identities = 394/399 (99%), Gaps = 3/399 (1%) | Note=member of SPL gene family, encodes DNA binding proteins and putative transcription factors. All have the SBP-box, which encodes the SBP-domain, required for and sufficient for interaction with DNA. \| squamosa promoter binding protein-like 1 (SPL1) \| FUNCTIONS IN: DNA binding, sequence-specific DNA binding transcription factor activity \| INVOLVED IN: regulation of transcription \| LOCATED IN: nucleus \| EXPRESSED IN: 22 plant structures \| EXPRESSED DURING: 13 growth stages \| CONTAINS InterPro DOMAIN/s: Transcription factor, SBP-box , Ankyrin repeat-containing domain , Ankyrin repeat \| BEST Arabidopsis thaliana protein match is: squamosa promoter-binding protein-like 12 | Note=Squamosa promoter binding-like protein (AHRD V1 ***- Q52UU1_9ROSI)%3B contains Interpro domain(s) IPR004333 Transcription factor%2C SBP-box ;Ontology_term=GO:0003700;from_BOGAS=1;interpro2go_term=GO:0005634;length=1170;nb_exon=2;sifter_term=GO:0003700 | | missense_variant | Qty: 2 |
| 21 | Query= Sopen07g031180.1<br>Length=444 | >Solyc07g062990.1.1<br>Length=465<br>Score = 385 bits (208), Expect = 8e-109<br>Identities = 227/236 (96%), Gaps = 1/236 (0%)<br>Score = 207 bits (112), Expect = 2e-55<br>Identities = 118/121 (98%), Gaps = 0/121 (0%) | Note=hypothetical protein | Note=Unknown Protein (AHRD V1);eugene_evidence_code=10F0H0E0IEG;from_BOGAS=1;length=1881;nb_exon=2 | | frameshift_variant<br>splice_acceptor_variant&intron_variant<br>missense_variant | Lys151fs<br><br>Qty: 4 |
| 22 | Query= Sopen07g031190.1<br>Length=3111 | >Solyc07g063000.2.1<br>Length=3054<br>Score = 4595 bits (2488), Expect = 0.0<br>Identities = 2538/2562 (99%), Gaps = 4/2562 (0%)<br>Score = 874 bits (473), Expect = 0.0<br>Identities = 482/486 (99%), Gaps = 1/486 (0%) | Note=Encodes a leucine-rich repeat receptor kinase (LRR-RK) involved in the perception of phytosulfokine (PSK), which is a 5-aa tyrosine-sulfated peptide that primarily promotes cellular proliferation. \| phytosylfokine-alpha receptor 2 (PSKR2) \| FUNCTIONS IN: peptide receptor activity, protein serine/threonine kinase activity, ATP binding \| INVOLVED IN: protein amino acid phosphorylation, transmembrane receptor protein tyrosine kinase signaling pathway, response to wounding \| LOCATED IN: chloroplast, plasma membrane \| EXPRESSED IN: 22 plant structures \| EXPRESSED DURING: 13 growth stages \| CONTAINS InterPro DOMAIN/s: Protein kinase, ATP binding site , Serine/threonine-protein kinase domain , Leucine-rich repeat-containing N-terminal domain, type 2 , Leucine-rich repeat , Serine/threonine-protein kinase-like domain , Serine/threonine-protein kinase, active site , Protein kinase-like domain , Protein kinase, catalytic domain , Leucine-rich repeat, typical subtype , Tyrosine-protein kinase, catalytic domain \| BEST Arabidopsis thaliana protein match is: phytosulfokin receptor 1 | Note=Receptor like kinase%2C RLK;Ontology_term=GO:0001653;from_BOGAS=1;interpro2go_term=GO:0005515,GO:0006468;length=3636;nb_exon=2;sifter_term=GO:0001653 | | missense_variant | Qty: 6 |
| 23 | Query= Sopen07g031200.1<br>Length=891 | >Solyc07g063010.2.1<br>Length=891<br>Score = 1485 bits (804), Expect = 0.0<br>Identities = 862/891 (97%), Gaps = 0/891 (0%) | Note=3-hydroxyacyl-CoA dehydrogenase family protein \| FUNCTIONS IN: in 7 functions \| INVOLVED IN: fatty acid metabolic process, metabolic process \| LOCATED IN: peroxisome \| EXPRESSED IN: 24 plant structures \| EXPRESSED DURING: 15 growth stages \| CONTAINS InterPro DOMAIN/s: Dehydrogenase, multihelical , 6-phosphogluconate dehydrogenase, C-terminal-like , 3-hydroxyacyl-CoA dehydrogenase, NAD binding , 3-hydroxyacyl-CoA dehydrogenase, C-terminal , 3-hydroxyacyl-CoA dehydrogenase, conserved site , NAD(P)-binding domain , 3-hydroxyacyl-CoA dehydrogenase \| BEST Arabidopsis thaliana protein match is: multifunctional protein 2 | Note=Fatty acid oxidation complex subunit alpha (AHRD V1 **-- FADB_ACIBC)%3B contains Interpro domain(s) IPR006176 3-hydroxyacyl-CoA dehydrogenase%2C NAD binding ;Ontology_term=GO:0042803;from_BOGAS=1;interpro2go_term=GO:0050662,GO:0008152,GO:0006631,GO:0016491;length=3127;nb_exon=4;sifter_term=GO:0042803 | | missense_variant | Qty: 3 |
| 24 | Query= Sopen07g031210.1<br>Length=3111 | >Solyc07g063020.1.1<br>Length=2952<br>Score = 3727 bits (2018), Expect = 0.0<br>Identities = 2160/2230 (97%), Gaps = 3/2230 (0%)<br>Score = 2508 bits (1358), Expect = 0.0<br>Identities = 1632/1761 (93%), Gaps = 31/1761 (2%) | Note=early nodulin-like protein 3 (ENODL3) \| FUNCTIONS IN: electron carrier activity, copper ion binding \| LOCATED IN: anchored to membrane \| CONTAINS InterPro DOMAIN/s: Plastocyanin-like , Cupredoxin \| BEST Arabidopsis thaliana protein match is: early nodulin-like protein 4 | Note=CT099 (Fragment) (AHRD V1 *-*- Q4KR07_SOLPE)%3B contains Interpro domain(s) IPR008972 Cupredoxin ;Ontology_term=GO:0009055;eugene_evidence_code=10F0H1E0IEG;from_BOGAS=1;interpro2go_term=GO:0009055;length=3261;nb_exon=3 | | missense_variant | Qty: 28 |

**Supplemental Table S3 (cont.). SIMILARITIES AND DISCREPANCIES BETWEEN THE CODING SEQUENCES OF S.PENNELLII VS S.LYCOPERSICUM (SL2.50) AS DETERMINED BY BLAST.**

| | | | | | | | |
|---|---|---|---|---|---|---|---|
| 25 | Query=Sopen07g031220.1<br>Length=387 | >Solyc07g063030.1.1<br>Length=387<br>Score = 671 bits (363), Expect = 0.0<br>Identities = 379/387 (98%), Gaps = 0/387 (0%) | Note=hypothetical protein | Note=Rapid alkalinization factor 2 (AHRD V1 **-- Q6TF28_SOLCH)%3B contains Interpro domain(s) IPR008801 Rapid ALkalinization Factor ;Ontology_term=GO:0004871;eugene_evidence_code=10F0H0E0IEG;from_BOGAS=1;length=387;nb_exon=1;sifter_term=GO:0004871 | | missense_variant | Qty: 4 |
| 26 | Query=Sopen07g031230.1<br>Length=1527 | >Solyc07g063040.2.1<br>Length=1527<br>Score = 2760 bits (1494), Expect = 0.0<br>Identities = 1516/1527 (99%), Gaps = 0/1527 (0%) | Note=haloacid dehalogenase-like hydrolase family protein \| FUNCTIONS IN: hydrolase activity, ion binding, methylthioribulose 1-phosphate dehydratase activity, metal ion binding \| INVOLVED IN: L-methionine salvage \| LOCATED IN: chloroplast \| EXPRESSED IN: 23 plant structures \| EXPRESSED DURING: 13 growth stages \| CONTAINS InterPro DOMAIN/s: Class II aldolase/adducin, N-terminal , Methylthioribulose-1-phosphate dehydratase | Note=Enolase-phosphatase E-1 (AHRD V1 *-*- A8UQN2_9AQUI)%3B contains Interpro domain(s) IPR017714 Methylthioribulose-1-phosphate dehydratase ;Ontology_term=GO:0046872,GO:0008152,GO:0005737,GO:0019509;from_BOGAS=1;interpro2go_term=GO:0046872,GO:0008152,GO:0005737,GO:0019509;length=5720;nb_exon=12 | | missense_variant | Qty: 3 |
| 27 | Query=Sopen07g031240.1<br>Length=834 | >Solyc07g063050.1.1<br>Length=834<br>Score = 1513 bits (819), Expect = 0.0<br>Identities = 829/834 (99%), Gaps = 0/834 (0%) | Note=unknown protein \| BEST Arabidopsis thaliana protein match is: unknown protein | Note=Unknown Protein (AHRD V1);eugene_evidence_code=10F0H0E1IEG;from_BOGAS=1;length=834;nb_exon=1 | | missense_variant | Qty: 3 |
| 28 | Query=Sopen07g031250.1<br>Length=282 | >Solyc07g063060.1.1<br>Length=384<br>Score = 255 bits (138), Expect = 4e-70<br>Identities = 146/150 (97%), Gaps = 0/150 (0%)<br>>Solyc07g063040.2.1<br>Length=1527<br>Score = 217 bits (117), Expect = 2e-58<br>Identities = 139/150 (93%), Gaps = 0/150 (0%) | Note=haloacid dehalogenase-like hydrolase family protein \| FUNCTIONS IN: hydrolase activity, ion binding, methylthioribulose 1-phosphate dehydratase activity, metal ion binding \| INVOLVED IN: L-methionine salvage \| LOCATED IN: chloroplast \| EXPRESSED IN: 23 plant structures \| EXPRESSED DURING: 13 growth stages \| CONTAINS InterPro DOMAIN/s: Class II aldolase/adducin, N-terminal , Methylthioribulose-1-phosphate dehydratase | Note=Os01g0101200 protein (Fragment) (AHRD V1 *-*- Q0JRH3_ORYSJ);eugene_evidence_code=10F0H1E0IEG;from_BOGAS=1;length=3270;nb_exon=5<br>Note=Enolase-phosphatase E-1 (AHRD V1 *-*- A8UQN2_9AQUI)%3B contains Interpro domain(s) IPR017714 Methylthioribulose-1-phosphate dehydratase ;Ontology_term=GO:0046872,GO:0008152,GO:0005737,GO:0019509;from_BOGAS=1;interpro2go_term=GO:0046872,GO:0008152,GO:0005737,GO:0019509;length=5720;nb_exon=12 | | start_lost<br>stop_gained<br>stop_lost&splice_region_variant<br>missense_variant<br>missense_variant | Met1?<br>Gln89*<br>Ter128Trpext*?<br>Qty: 5<br>Qty: 3 |
| 29 | Query=Sopen07g031260.1<br>Length=696 | >Solyc07g063070.1.1<br>Length=702<br>Score = 1214 bits (657), Expect = 0.0<br>Identities = 688/702 (98%), Gaps = 6/702 (1%) | Note=VQ motif-containing protein \| FUNCTIONS IN: molecular_function unknown \| INVOLVED IN: biological_process unknown \| EXPRESSED IN: 22 plant structures \| EXPRESSED DURING: 13 growth stages \| CONTAINS InterPro DOMAIN/s: VQ \| BEST Arabidopsis thaliana protein match is: VQ motif-containing protein | Note=DNA-binding WRKY VQ (AHRD V1 ***- Q2HTS9_MEDTR)%3B contains Interpro domain(s) IPR008889 VQ ;eugene_evidence_code=10F1H1E1IEG;from_BOGAS=1;length=702;nb_exon=1 | | missense_variant | Qty: 2 |
| 30 | Query=Sopen07g031270.1<br>Length=2478 | >Solyc07g063080.2.1<br>Length=2490<br>Score = 4394 bits (2379), Expect = 0.0<br>Identities = 2455/2490 (99%), Gaps = 12/2490 (0%) | Note=Encodes DRP5A, a dynamin protein involved in cytokinesis in Arabidopsis. \| Dynamin related protein 5A (DRP5A) \| FUNCTIONS IN: GTP binding, GTPase activity \| INVOLVED IN: cytokinesis \| EXPRESSED IN: 18 plant structures \| EXPRESSED DURING: 6 growth stages \| CONTAINS InterPro DOMAIN/s: Dynamin, GTPase domain \| BEST Arabidopsis thaliana protein match is: P-loop containing nucleoside triphosphate hydrolases superfamily protein | Note=Dynamin family protein (AHRD V1 ***- D7KKE2_ARALY)%3B contains Interpro domain(s) IPR001401 Dynamin%2C GTPase region ;Ontology_term=GO:0005525;from_BOGAS=1;interpro2go_term=GO:0005525;length=4658;nb_exon=8 | | missense_variant | Qty: 6 |
| 31 | Query=Sopen07g031280.1<br>Length=441 | >Solyc07g063090.1.1<br>Length=441<br>Score = 809 bits (438), Expect = 0.0<br>Identities = 440/441 (99%), Gaps = 0/441 (0%) | Note=Encodes GRIM REAPER (GRI), involved in the regulation of cell death induced by extracellular ROS (reactive oxygen species). Secreted into the extracellular space. \| GRIM REAPER (GRI) \| CONTAINS InterPro DOMAIN/s: Stigma-specific protein Stig1 \| BEST Arabidopsis thaliana protein match is: Stigma-specific Stig1 family protein | Note=Stig1 (Fragment) (AHRD V1 *-*- Q9SBR0_PETHY)%3B contains Interpro domain(s) IPR006969 Stigma-specific protein Stig1 ;eugene_evidence_code=10F0H1E0IEG;from_BOGAS=1;length=441;nb_exon=1 | | | |
| 32 | Query=Sopen07g031290.1<br>Length=1890 | >Solyc07g063100.2.1<br>Length=1887<br>Score = 3410 bits (1846), Expect = 0.0<br>Identities = 1876/1890 (99%), Gaps = 3/1890 (0%) | Note=interacts with H+-ATPase, and regulates its activity \| proton pump interactor 1 (PPI1) \| FUNCTIONS IN: protein binding \| INVOLVED IN: regulation of proton transport \| LOCATED IN: endoplasmic reticulum, plasma membrane \| EXPRESSED IN: 25 plant structures \| EXPRESSED DURING: 15 growth stages \| BEST Arabidopsis thaliana protein match is: unknown protein | Note=Proton pump interactor 1 (AHRD V1 **-- D5L6G0_SOLTU);from_BOGAS=1;length=3849;nb_exon=7 | | missense_variant | Qty: 4 |
| 33 | Query=Sopen07g031300.1<br>Length=1170 | >Solyc07g063110.2.1<br>Length=777<br>Score = 1214 bits (657), Expect = 0.0<br>Identities = 678/687 (99%), Gaps = 5/687 (1%) | Note=Core-2/I-branching beta-1,6-N-acetylglucosaminyltransferase family protein \| FUNCTIONS IN: transferase activity, transferring glycosyl groups, acetylglucosaminyltransferase activity \| INVOLVED IN: carbohydrate biosynthetic process \| LOCATED IN: endomembrane system, membrane \| EXPRESSED IN: 12 plant structures \| EXPRESSED DURING: 8 growth stages \| CONTAINS InterPro DOMAIN/s: Glycosyl transferase, family 14 , Core-2/I-Branching enzyme \| BEST Arabidopsis thaliana protein match is: Core-2/I-branching beta-1,6-N-acetylglucosaminyltransferase family protein | Note=Beta-1%2C3-galactosyl-O-glycosyl-glycoprotein beta-1%2C6-N-acetylglucosaminyltransferase (AHRD V1 *--* GCNT1_BOVIN)%3B contains Interpro domain(s) IPR003406 Glycosyl transferase%2C family 14 ;Ontology_term=GO:0016757;from_BOGAS=1;interpro2go_term=GO:0016020;length=2326;nb_exon=5;sifter_term=GO:0016757 | | | |
| 34 | Query=Sopen07g031310.1<br>Length=2193<br><br><br><br><br>Query=Sopen07g031310.2<br>Length=2646 | >Solyc07g063120.2.1<br>Length=2493<br>Score = 3334 bits (1805), Expect = 0.0<br>Identities = 1892/1934 (98%), Gaps = 6/1934 (0%)<br>Score = 110 bits (59), Expect = 3e-25<br>Identities = 61/62 (98%), Gaps = 0/62 (0%)<br>>Solyc07g063120.2.1<br>Length=2493<br>Score = 4346 bits (2353), Expect = 0.0<br>Identities = 2448/2494 (98%), Gaps = 6/2494 (0%) | Note=Encodes a member of the SPA (suppressor of phyA-105) protein family (SPA1-SPA4). SPA proteins contain an N-terminal serine/threonine kinase-like motif followed by a coiled-coil structure and a C-terminal WD-repeat domain. SPA proteins function redundantly in suppressing photomorphogenesis in dark- and light-grown seedlings. SPA3 (and SPA4) predominantly regulates elongation growth in adult plants. \| SPA1-related 3 (SPA3) \| FUNCTIONS IN: protein binding, signal transducer activity \| INVOLVED IN: protein amino acid phosphorylation \| LOCATED IN: CUL4 RING ubiquitin ligase complex, heterotrimeric G-protein complex \| CONTAINS InterPro DOMAIN/s: WD40 repeat 2 , WD40 repeat, conserved site , Serine/threonine-protein kinase domain , WD40 repeat , Serine/threonine-protein kinase-like domain , Protein kinase-like domain , WD40 repeat-like-containing domain , Protein kinase, catalytic domain , WD40-repeat-containing domain , WD40/YVTN repeat-like-containing domain , WD40 repeat, subgroup \| BEST Arabidopsis thaliana protein match is: SPA1-related 4 | Note=WD-40 repeat protein (AHRD V1 *--- B7KCU4_CYAP7)%3B contains Interpro domain(s) IPR020472 G-protein beta WD-40 repeat%2C region ;Ontology_term=GO:0042802;from_BOGAS=1;interpro2go_term=GO:0006468;length=8897;nb_exon=9;sifter_term=GO:0042802 | | missense_variant<br><br><br><br><br><br>missense_variant | Qty: 13<br><br><br><br><br><br>Qty: 13 |
| 35 | Query=Sopen07g031320.1<br>Length=2118 | >Solyc07g063130.2.1<br>Length=2169<br>Score = 3829 bits (2073), Expect = 0.0<br>Identities = 2103/2118 (99%), Gaps = 0/2118 (0%) | Note=Protein kinase superfamily protein \| FUNCTIONS IN: protein serine/threonine kinase activity, protein kinase activity, kinase activity, ATP binding \| INVOLVED IN: protein amino acid phosphorylation, N-terminal protein myristoylation \| LOCATED IN: plasma membrane \| EXPRESSED IN: 24 plant structures \| EXPRESSED DURING: 13 growth stages \| CONTAINS InterPro DOMAIN/s: Protein kinase, ATP binding site , Protein kinase, catalytic domain , Serine/threonine-protein kinase domain , Serine/threonine-protein kinase-like domain , Protein kinase-like domain , Serine/threonine-protein kinase, active site \| BEST Arabidopsis thaliana protein match is: Protein kinase superfamily protein | Note=Cell division protein kinase 10 (AHRD V1 *-*- D3TRH8_GLOMM)%3B contains Interpro domain(s) IPR002290 Serine/threonine protein kinase ;Ontology_term=GO:0006468;from_BOGAS=1;interpro2go_term=GO:0006468;length=8368;nb_exon=9 | | stop_gained&splice_region_variant<br>missense_variant | Ser17*<br>Qty: 7 |
| 36 | Query=Sopen07g031330.1<br>Length=423 | >Solyc07g063140.1.1<br>Length=423<br>Score = 737 bits (399), Expect = 0.0<br>Identities = 415/423 (98%), Gaps = 0/423 (0%) | Note=FUNCTIONS IN: molecular_function unknown \| INVOLVED IN: regulation of translational fidelity \| LOCATED IN: chloroplast \| EXPRESSED IN: 22 plant structures \| EXPRESSED DURING: 13 growth stages \| CONTAINS InterPro DOMAIN/s: Glu-tRNAGln amidotransferase, C subunit | Note=Aspartyl/glutamyl-tRNA(Asn/Gln) amidotransferase subunit C (AHRD V1 ***- A3IGR0_9CHRO)%3B contains Interpro domain(s) IPR003837 Glu-tRNAGln amidotransferase%2C C subunit ;Ontology_term=GO:0006450;eugene_evidence_code=10F1H1E1IEG;from_BOGAS=1;interpro2go_term=GO:0006450;length=423;nb_exon=1 | | missense_variant | Qty: 5 |
| 37 | Query=Sopen07g031340.1<br>Length=1725 | >Solyc07g063150.2.1<br>Length=1326<br>Score = 2405 bits (1302), Expect = 0.0<br>Identities = 1318/1326 (99%), Gaps = 0/1326 (0%) | Note=Protein of unknown function (DUF616) \| CONTAINS InterPro DOMAIN/s: Protein of unknown function DUF616 \| BEST Arabidopsis thaliana protein match is: Protein of unknown function (DUF616) | Note=Os11g0198100 protein (Fragment) (AHRD V1 *--- Q0IU20_ORYSJ)%3B contains Interpro domain(s) IPR006852 Protein of unknown function DUF616 ;from_BOGAS=1;length=4157;nb_exon=8 | | missense_variant | Qty: 5 |
| 38 | Query=Sopen07g031350.1<br>Length=267 | >Solyc07g063160.2.1<br>Length=408<br>Score = 455 bits (246), Expect = 3e-130<br>Identities = 260/267 (97%), Gaps = 0/267 (0%) | Note=Encodes protein similar to yeast COX17, a copper-binding protein that mediates the delivery of Cu to the mitochondria for the assembly of a functional cytochrome oxidase complex. \| cytochrome c oxidase 17 (COX17) \| FUNCTIONS IN: copper chaperone activity \| INVOLVED IN: response to bacterium, response to copper ion \| LOCATED IN: mitochondrial intermembrane space \| EXPRESSED IN: 22 plant structures \| EXPRESSED DURING: 13 growth stages \| CONTAINS InterPro DOMAIN/s: Cytochrome c oxidase copper chaperone \| BEST Arabidopsis thaliana protein match is: Cytochrome C oxidase copper chaperone (COX17) | Note=Cytochrome c oxidase copper chaperone family protein (AHRD V1 *-*- D7KKD3_ARALY)%3B contains Interpro domain(s) IPR007745 Cytochrome c oxidase copper chaperone ;Ontology_term=GO:0005758;from_BOGAS=1;interpro2go_term=GO:0005758;length=3650;nb_exon=3 | | frameshift_variant<br>stop_gained<br>missense_variant | Phe25fs<br>Leu42*<br>Qty: 3 |
| 39 | Query=Sopen07g031360.1<br>Length=519 | >Solyc07g063170.1.1<br>Length=507<br>Score = 804 bits (435), Expect = 0.0<br>Identities = 493/519 (95%), Gaps = 12/519 (2%) | Note=hypothetical protein | Note=Unknown Protein (AHRD V1);eugene_evidence_code=10F0H0E0IEG;from_BOGAS=1;length=507;nb_exon=1 | | missense_variant | Qty: 9 |
| 40 | Query=Sopen07g031370.1<br>Length=444 | >Solyc07g063180.2.1<br>Length=444<br>Score = 782 bits (423), Expect = 0.0<br>Identities = 437/444 (98%), Gaps = 0/444 (0%) | Note=Dynein light chain type 1 family protein \| FUNCTIONS IN: microtubule motor activity \| INVOLVED IN: microtubule-based process \| LOCATED IN: microtubule associated complex \| EXPRESSED IN: 23 plant structures \| EXPRESSED DURING: 13 growth stages \| CONTAINS InterPro DOMAIN/s: Dynein light chain, type 1/2 \| BEST Arabidopsis thaliana protein match is: Dynein light chain type 1 family protein | Note=Cytoplasmic dynein light chain (AHRD V1 *-*- D3BU56_POLPA)%3B contains Interpro domain(s) IPR001372 Dynein light chain%2C type 1 and 2 ;Ontology_term=GO:0042803;from_BOGAS=1;interpro2go_term=GO:0005875;length=775;nb_exon=2;sifter_term=GO:0042803 | | missense_variant | Qty: 1 |
| 41 | Query=Sopen07g031380.1<br>Length=549 | >Solyc07g063190.2.1<br>Length=549<br>Score = 970 bits (525), Expect = 0.0<br>Identities = 541/549 (99%), Gaps = 0/549 (0%) | Note=encodes a prokaryotic thioredoxin \| thioredoxin M-type 4 (TRX-M4) \| FUNCTIONS IN: enzyme activator activity \| INVOLVED IN: response to oxidative stress, positive regulation of catalytic activity \| LOCATED IN: thylakoid, chloroplast thylakoid membrane, cell wall, chloroplast, chloroplast envelope \| EXPRESSED IN: 24 plant structures \| EXPRESSED DURING: 13 growth stages \| CONTAINS InterPro DOMAIN/s: Thioredoxin fold , Thioredoxin, core , Thioredoxin domain , Thioredoxin, conserved site , Thioredoxin , Thioredoxin-like subdomain , Thioredoxin-like , Thioredoxin-like fold \| BEST Arabidopsis thaliana protein match is: thioredoxin M-type 1 | Note=Thioredoxin (AHRD V1 ***- A7LNX7_9CARY)%3B contains Interpro domain(s) IPR005746 Thioredoxin ;Ontology_term=GO:0004857,GO:0008047;from_BOGAS=1;interpro2go_term=GO:0045454;length=2507;nb_exon=2;sifter_term=GO:0004857,GO:0008047 | | missense_variant | Qty: 4 |
| 42 | Query=Sopen07g031390.1<br>Length=750 | >Solyc07g063200.1.1<br>Length=717<br>Score = 1286 bits (696), Expect = 0.0<br>Identities = 710/717 (99%), Gaps = 0/717 (0%) | Note=Encodes a RING finger E3 ubiquitin ligase. \| RING membrane-anchor 3 (RMA3) \| CONTAINS InterPro DOMAIN/s: Zinc finger, RING-type, conserved site , Zinc finger, RING-type , Zinc finger, C3HC4 RING-type \| BEST Arabidopsis thaliana protein match is: RING membrane-anchor 1 | Note=RING finger protein 5 (AHRD V1 *-*- B6TF06_MAIZE)%3B contains Interpro domain(s) IPR018957 Zinc finger%2C C3HC4 RING-type ;Ontology_term=GO:0004842;eugene_evidence_code=10F0H1E1IEG;from_BOGAS=1;interpro2go_term=GO:0008270;length=717;nb_exon=1;sifter_term=GO:0004842 | | missense_variant | Qty: 4 |
| 43 | Query=Sopen07g031400.1<br>Length=771 | >Solyc07g063210.2.1<br>Length=771<br>Score = 1369 bits (741), Expect = 0.0<br>Identities = 761/771 (99%), Gaps = 0/771 (0%) | Note=member of Alpha-Expansin Gene Family. Naming convention from the Expansin Working Group (Kende et al, 2004. Plant Mol Bio) \| expansin 12 (EXPA12) \| INVOLVED IN: plant-type cell wall organization, unidimensional cell growth, plant-type cell wall loosening \| LOCATED IN: endomembrane system, extracellular region \| EXPRESSED IN: root \| CONTAINS InterPro DOMAIN/s: Barwin-related endoglucanase , Pollen allergen, N-terminal , Expansin , Rare lipoprotein A , Expansin/Lol pI , Expansin 45, endoglucanase-like , Pollen allergen/expansin, C-terminal \| BEST Arabidopsis thaliana protein match is: expansin A17 | Note=Expansin S2 (AHRD V1 ***- Q39626_CUCSA)%3B contains Interpro domain(s) IPR002963 Expansin ;Ontology_term=GO:0005199;from_BOGAS=1;interpro2go_term=GO:0005576,GO:0009664;length=1951;nb_exon=3;sifter_term=GO:0005199 | | missense_variant | Qty: 1 |
| 44 | Query=Sopen07g031410.1<br>Length=1524 | >Solyc07g063220.2.1<br>Length=1668<br>Score = 2699 bits (1461), Expect = 0.0<br>Identities = 1503/1523 (99%), Gaps = 3/1523 (0%) | Note=Tetratricopeptide repeat (TPR)-like superfamily protein \| CONTAINS InterPro DOMAIN/s: Pentatricopeptide repeat \| BEST Arabidopsis thaliana protein match is: Tetratricopeptide repeat (TPR)-like superfamily protein | Note=Pentatricopeptide repeat-containing protein (AHRD V1 ***- D7MEQ2_ARALY)%3B contains Interpro domain(s) IPR002885 Pentatricopeptide repeat ;from_BOGAS=1;length=3362;nb_exon=3 | | stop_gained<br>missense_variant | Arg531*<br>Qty: 5 |
| 45 | Query=Sopen07g031420.1<br>Length=2100 | >Solyc07g063230.2.1<br>Length=2100<br>Score = 3762 bits (2037), Expect = 0.0<br>Identities = 2079/2100 (99%), Gaps = 0/2100 (0%) | Note=Plasma-membrane choline transporter family protein \| INVOLVED IN: biological_process unknown \| LOCATED IN: plasma membrane \| EXPRESSED IN: 22 plant structures \| EXPRESSED DURING: 13 growth stages \| CONTAINS InterPro DOMAIN/s: Protein of unknown function DUF580 | Note=Choline transporter like family (AHRD V1 ***- C1E2U6_9CHLO)%3B contains Interpro domain(s) IPR007603 Protein of unknown function DUF580 ;from_BOGAS=1;length=5553;nb_exon=10 | | missense_variant | Qty: 6 |
| 46 | Query=Sopen07g031430.1<br>Length=1560 | >Solyc07g063240.2.1<br>Length=1560<br>Score = 2760 bits (1494), Expect = 0.0<br>Identities = 1538/1560 (99%), Gaps = 0/1560 (0%) | Note=Encodes a novel protein that is similar to PRL1 interacting factor and is involved in virus induced silencing. \| silencing defective 5 (SDE5) \| FUNCTIONS IN: molecular_function unknown \| INVOLVED IN: production of ta-siRNAs involved in RNA interference \| LOCATED IN: cellular_component unknown \| EXPRESSED IN: 21 plant structures \| EXPRESSED DURING: 12 growth stages \| CONTAINS InterPro DOMAIN/s: Domain of unknown function DUF1771 \| BEST Arabidopsis thaliana protein match is: smr (Small MutS Related) domain-containing protein . | Note=Genomic DNA chromosome 3 P1 clone MJK13 (AHRD V1 **-- Q9LUQ3_ARATH)%3B contains Interpro domain(s) IPR013899 Region of unknown function DUF1771 ;from_BOGAS=1;length=9442;nb_exon=12 | | missense_variant | Qty: 6 |
| 47 | Query=Sopen07g031440.1<br>Length=1689 | >Solyc07g063260.2.1<br>Length=1692<br>Score = 3033 bits (1642), Expect = 0.0<br>Identities = 1676/1692 (99%), Gaps = 3/1692 (0%) | Note=A member of a large family of seven-transmembrane domain proteins specific to plants, homologs of the barley mildew resistance locus o (MLO) protein. The Arabidopsis genome contains 15 genes encoding MLO proteins, with localization in plasma membrane. Phylogenetic analysis revealed four clades of closely-related AtMLO genes. ATMLO11 belongs to the clade I, with AtMLO4 and AtMLO14. The gene is expressed during early seedling growth (in primary root), in root tips and lateral root primordia, and in very young leaves, and in flowers and fruit abscission zone, as shown by GUS activity patterns. The expression of several phylogenetically closely-related AtMLO genes showed similar or overlapping tissue specificity and analogous responsiveness to external stimuli, suggesting functional redundancy, co-function, or antagonistic function(s). \| MILDEW RESISTANCE LOCUS O 11 (MLO11) \| FUNCTIONS IN: calmodulin binding \| INVOLVED IN: cell death, defense response \| LOCATED IN: integral to membrane, plasma membrane \| EXPRESSED IN: 25 plant structures \| EXPRESSED DURING: 10 growth stages \| CONTAINS InterPro DOMAIN/s: Mlo-related protein \| BEST Arabidopsis thaliana protein match is: Seven transmembrane MLO family protein | Note=MLO-like protein 14 (AHRD V1 ***- B6SWK8_MAIZE)%3B contains Interpro domain(s) IPR004326 Mlo-related protein ;Ontology_term=GO:0016021;from_BOGAS=1;interpro2go_term=GO:0016021;length=7556;nb_exon=15 | | missense_variant | Qty: 4 |
| 48 | Query=Sopen07g031450.1<br>Length=1551 | >Solyc07g063270.2.1<br>Length=1551<br>Score = 2798 bits (1515), Expect = 0.0<br>Identities = 1539/1551 (99%), Gaps = 0/1551 (0%) | Note=GTP-binding family protein \| FUNCTIONS IN: GTP binding \| INVOLVED IN: biological_process unknown \| LOCATED IN: intracellular \| EXPRESSED IN: 23 plant structures \| EXPRESSED DURING: 13 growth stages \| CONTAINS InterPro DOMAIN/s: GTP1/OBG , GTP-binding protein, HSR1-related , NGP1, N-terminal \| BEST Arabidopsis thaliana protein match is: GTP-binding family protein | Note=Nucleolar GTP-binding protein 2 (AHRD V1 **** B6TMG1_MAIZE)%3B contains Interpro domain(s) IPR012971 NGP1%2C N-terminal ;Ontology_term=GO:0005515,GO:0003924;from_BOGAS=1;interpro2go_term=GO:0005525,GO:0005622;length=5934;nb_exon=12;sifter_term=GO:0005515,GO:0003924 | | missense_variant | Qty: 1 |
| 49 | Query=Sopen07g031460.1<br>Length=417 | >Solyc07g063290.2.1<br>Length=405<br>Score = 673 bits (364), Expect = 0.0<br>Identities = 399/414 (96%), Gaps = 9/414 (2%) | Note=Transcription factor TFIIIC, tau55-related protein \| CONTAINS InterPro DOMAIN/s: Transcription factor TFIIIC, tau55-related \| BEST Arabidopsis thaliana protein match is: Transcription factor TFIIIC, tau55-related protein | Note=Unknown Protein (AHRD V1)%3B contains Interpro domain(s) IPR019481 Transcription factor TFIIIC%2C tau55-related ;from_BOGAS=1;length=4090;nb_exon=6 | | frameshift_variant<br>missense_variant | Glu78fs<br>Qty: 2 |
| 50 | Query=Sopen07g031470.1<br>Length=1446 | >Solyc07g063300.2.1<br>Length=1461<br>Score = 2599 bits (1407), Expect = 0.0<br>Identities = 1431/1443 (99%), Gaps = 0/1443 (0%) | Note=Regulator of chromosome condensation (RCC1) family protein \| CONTAINS InterPro DOMAIN/s: Regulator of chromosome condensation/beta-lactamase-inhibitor protein II , Regulator of chromosome condensation, RCC1 \| BEST Arabidopsis thaliana protein match is: Regulator of chromosome condensation (RCC1) family protein | Note=Regulator of chromosome condensation (RCC1)-like protein (AHRD V1 **-- D0NEM9_PHYIN)%3B contains Interpro domain(s) IPR009091 Regulator of chromosome condensation/beta-lactamase-inhibitor protein II ;Ontology_term=GO:0005087;from_BOGAS=1;length=4639;nb_exon=6;sifter_term=GO:0005087 | | missense_variant | Qty: 4 |
| 51 | Query=Sopen07g031480.1<br>Length=1227 | >Solyc07g063310.2.1<br>Length=1227<br>Score = 2172 bits (1176), Expect = 0.0<br>Identities = 1210/1227 (99%), Gaps = 0/1227 (0%) | Note=Encodes a plasma membraneÃ±localized ABA receptor, which interacts with the G, #945,, #946,, #947 \| complex. It has been postulated that the binding of ABA to GCR2 results in the release of the G protein and dissociation of the heterotrimeric complex into G, #945 \| and the G, #946,, #947 \| dimer to activate downstream ABA effectors and to trigger the ABA responses. \| G protein coupled receptor (GPCR) \| CONTAINS InterPro DOMAIN/s: LanC-like protein, eukaryotic , Six-hairpin glycosidase-like , Lanthionine synthetase C-like \| BEST Arabidopsis thaliana protein match is: GCR2-like 2 | Note=LanC-like protein 2 (AHRD V1 ***- B6TCV9_MAIZE)%3B contains Interpro domain(s) IPR020464 LanC-like protein%2C eukaryotic ;Ontology_term=GO:0010427;from_BOGAS=1;interpro2go_term=GO:0003824;length=2662;nb_exon=6;sifter_term=GO:0010427 | | missense_variant | Qty: 4 |
| 52 | Query=Sopen07g031490.1<br>Length=1224 | >Solyc07g063320.2.1<br>Length=1224<br>Score = 2134 bits (1155), Expect = 0.0<br>Identities = 1201/1224 (98%), Gaps = 0/1224 (0%) | Note=Encodes a plasma membraneÃ±localized ABA receptor, which interacts with the G, #945,, #946,, #947 \| complex. It has been postulated that the binding of ABA to GCR2 results in the release of the G protein and dissociation of the heterotrimeric complex into G, #945 \| and the G, #946,, #947 \| dimer to activate downstream ABA effectors and to trigger the ABA responses. \| G protein coupled receptor (GPCR) \| CONTAINS InterPro DOMAIN/s: LanC-like protein, eukaryotic , Six-hairpin glycosidase-like , Lanthionine synthetase C-like \| BEST Arabidopsis thaliana protein match is: GCR2-like 2 | Note=LanC-like protein 2 (AHRD V1 ***- B6TCV9_MAIZE)%3B contains Interpro domain(s) IPR020464 LanC-like protein%2C eukaryotic ;Ontology_term=GO:0010427;from_BOGAS=1;interpro2go_term=GO:0003824;length=3269;nb_exon=6;sifter_term=GO:0010427 | | splice_acceptor_variant&intron_variant<br>missense_variant | <br>Qty: 4 |
| 53 | Query=Sopen07g031500.1<br>Length=2613 | >Solyc07g063330.2.1<br>Length=2613<br>Score = 4682 bits (2535), Expect = 0.0<br>Identities = 2587/2613 (99%), Gaps = 0/2613 (0%) | Note=Transducin/WD40 repeat-like superfamily protein \| CONTAINS InterPro DOMAIN/s: WD40 repeat 2 , WD40 repeat, conserved site , WD40 repeat , G-protein beta WD-40 repeat, region , WD40 repeat-like-containing domain , WD40-repeat-containing domain , WD40/YVTN repeat-like-containing domain , WD40 repeat, subgroup \| BEST Arabidopsis thaliana protein match is: Transducin/WD40 repeat-like superfamily protein | Note=WD-40 repeat family protein (AHRD V1 ***- D7L4T9_ARALY)%3B contains Interpro domain(s) IPR020472 G-protein beta WD-40 repeat%2C region ;Ontology_term=GO:0004871;from_BOGAS=1;length=5755;nb_exon=8;sifter_term=GO:0004871 | | missense_variant | Qty: 6 |
| 54 | Query=Sopen07g031510.1<br>Length=561 | >Solyc07g063340.2.1<br>Length=561<br>Score = 992 bits (537), Expect = 0.0<br>Identities = 553/561 (99%), Gaps = 0/561 (0%) | Note=Protein of unknown function (DUF1218) \| FUNCTIONS IN: molecular_function unknown \| INVOLVED IN: biological_process unknown \| LOCATED IN: endomembrane system \| EXPRESSED IN: 20 plant structures \| EXPRESSED DURING: 13 growth stages \| CONTAINS InterPro DOMAIN/s: Protein of unknown function DUF1218 \| BEST Arabidopsis thaliana protein match is: Protein of unknown function (DUF1218) | Note=COSII_At1g13380 (Fragment) (AHRD V1 *-*- C0KEY3_SOLDU)%3B contains Interpro domain(s) IPR009606 Protein of unknown function DUF1218 ;from_BOGAS=1;length=2222;nb_exon=4 | | missense_variant | Qty: 3 |
| 55 | Query=Sopen07g031520.1<br>Length=3318 | >Solyc07g063350.2.1<br>Length=3318<br>Score = 5823 bits (3153), Expect = 0.0<br>Identities = 3263/3318 (98%), Gaps = 0/3318 (0%) | Note=Positive regulator of light-regulated genes. Novel nuclear protein which requires light for its high level expression. \| COP1-interacting protein 7 (CIP7) \| FUNCTIONS IN: transcription activator activity \| INVOLVED IN: chlorophyll biosynthetic process, response to light stimulus, anthocyanin biosynthetic process \| LOCATED IN: nucleus, plasma membrane \| EXPRESSED IN: 24 plant structures \| EXPRESSED DURING: 13 growth stages \| BEST Arabidopsis thaliana protein match is: disease resistance protein (TIR class), putative | Note=COP1-Interacting ProteinI 7 (CIP7) (Fragment) (AHRD V1 *-*- Q7GDB1_ARATH);Ontology_term=GO:0016563;from_BOGAS=1;length=4795;nb_exon=9;sifter_term=GO:0016563 | | missense_variant | Qty: 17 |
| 56 | Query=Sopen07g031530.1<br>Length=459<br><br><br>Query=Sopen07g031530.2<br>Length=405 | >Solyc07g063370.2.1<br>Length=1566<br>Score = 808 bits (437), Expect = 0.0<br>Identities = 446/450 (99%), Gaps = 2/450 (0%)<br>>Solyc07g063370.2.1<br>Length=1566<br>Score = 651 bits (352), Expect = 0.0<br>Identities = 359/362 (99%), Gaps = 2/362 (1%) | Note=beta glucosidase 46 (BGLU46) \| FUNCTIONS IN: cation binding, hydrolase activity, hydrolyzing O-glycosyl compounds, catalytic activity \| INVOLVED IN: lignin biosynthetic process \| EXPRESSED IN: 12 plant structures \| EXPRESSED DURING: 4 anthesis, F mature embryo stage, petal differentiation and expansion stage, E expanded cotyledon stage, D bilateral stage \| CONTAINS InterPro DOMAIN/s: Glycoside hydrolase, family 1 , Glycoside hydrolase, family 1, active site , Glycoside hydrolase, catalytic core , Glycoside hydrolase, subgroup, catalytic core \| BEST Arabidopsis thaliana protein match is: beta-glucosidase 45 | Note=Beta-glucosidase (AHRD V1 **** B6SYQ7_MAIZE)%3B contains Interpro domain(s) IPR001360 Glycoside hydrolase%2C family 1 ;Ontology_term=GO:0047782;from_BOGAS=1;interpro2go_term=GO:0005975;length=3804;nb_exon=12;sifter_term=GO:0047782 | | missense_variant | Qty: 6 |
| 57 | Query=Sopen07g031540.1<br>Length=783 | >Solyc07g063380.1.1<br>Length=297<br>Score = 403 bits (218), Expect = 4e-114<br>Identities = 269/294 (91%), Gaps = 1/294 (0%) | Note=hypothetical protein | Note=Unknown Protein (AHRD V1);eugene_evidence_code=10F0H0E0IEG;from_BOGAS=1;length=297;nb_exon=1 | | | |
| 58 | Query=Sopen07g031550.1<br>Length=1593 | >Solyc07g063390.2.1<br>Length=1617<br>Score = 2732 bits (1479), Expect = 0.0<br>Identities = 1553/1590 (98%), Gaps = 0/1590 (0%) | Note=beta glucosidase 46 (BGLU46) \| FUNCTIONS IN: cation binding, hydrolase activity, hydrolyzing O-glycosyl compounds, catalytic activity \| INVOLVED IN: lignin biosynthetic process \| EXPRESSED IN: 12 plant structures \| EXPRESSED DURING: 4 anthesis, F mature embryo stage, petal differentiation and expansion stage, E expanded cotyledon stage, D bilateral stage \| CONTAINS InterPro DOMAIN/s: Glycoside hydrolase, family 1 , Glycoside hydrolase, family 1, active site , Glycoside hydrolase, catalytic core , Glycoside hydrolase, subgroup, catalytic core \| BEST Arabidopsis thaliana protein match is: beta-glucosidase 45 | Note=Beta-glucosidase (AHRD V1 **** B6SYQ7_MAIZE)%3B contains Interpro domain(s) IPR001360 Glycoside hydrolase%2C family 1 ;Ontology_term=GO:0047782;from_BOGAS=1;interpro2go_term=GO:0005975;length=3533;nb_exon=12;sifter_term=GO:0047782 | | missense_variant | Qty: 9 |

**Supplemental Table S4. POLYMORPHISMS WITH A MODERATE EFFECT ON GENE FUNCTION FOR S.PENNELLII VS S.LYCOPERSICUM (SL2.50) WITHIN THE *ORG* REGION.** The variant information was obtained from the S.pennellii resequencing dataset (Aflitos et al. 2014).

| Chromosome | Position | Ref allele | Alt allele | Variant type | Impact | HGVS-nomenclatre | Gene | Protein Description |
|---|---|---|---|---|---|---|---|---|
| **SL2.50ch07** | **65444176** | **A** | **G** | | | | **LEFT BORDER** | |
| SL2.50ch07 | 65446115 | C | G | missense_variant | MODERATE | Gln192Glu | Solyc07g062750.2 | ERI1 exoribonuclease 3 (AHRD V1 **-- ERI3_BOVIN)%3B contains Interpro domain(s) IPR006055 Exonuclease |
| SL2.50ch07 | 65446691 | A | G | missense_variant | MODERATE | Thr224Ala | | |
| SL2.50ch07 | 65446945 | A | C | missense_variant | MODERATE | Lys278Asn | | |
| SL2.50ch07 | 65446986 | T | C | missense_variant | MODERATE | Val292Ala | | |
| SL2.50ch07 | 65449788 | T | G | missense_variant | MODERATE | Phe9Val | Solyc07g062760.1 | 26S protease regulatory subunit 4 homolog (AHRD V1 *--- PRS4_ORYSJ)%3B contains Interpro domain(s) IPR003959 ATPase%2C AAA-type%2C core |
| SL2.50ch07 | 65450033 | G | T | missense_variant | MODERATE | Arg90Ser | | |
| SL2.50ch07 | 65450194 | A | G | missense_variant | MODERATE | Asp144Gly | | |
| SL2.50ch07 | 65450269 | C | G | missense_variant | MODERATE | Ala169Gly | | |
| SL2.50ch07 | 65450388 | G | T | missense_variant | MODERATE | Gly209Cys | | |
| SL2.50ch07 | 65450569 | T | G | missense_variant | MODERATE | Phe269Cys | | |
| SL2.50ch07 | 65450634 | T | G | missense_variant | MODERATE | Trp291Gly | | |
| SL2.50ch07 | 65450877 | C | G | missense_variant | MODERATE | Pro372Ala | | |
| SL2.50ch07 | 65452364 | A | T | missense_variant | MODERATE | Glu568Asp | | |
| SL2.50ch07 | 65452553 | C | A | missense_variant | MODERATE | His631Gln | | |
| SL2.50ch07 | 65453368 | T | C | missense_variant | MODERATE | Val903Ala | | |
| SL2.50ch07 | 65453751 | A | G | missense_variant | MODERATE | Lys1031Glu | | |
| SL2.50ch07 | 65455295 | T | G | missense_variant | MODERATE | Asp11Glu | Solyc07g062770.2 | Unknown Protein (AHRD V1) |
| SL2.50ch07 | 65455521 | T | G | missense_variant | MODERATE | Leu87Val | | |
| SL2.50ch07 | 65456276 | G | T | missense_variant | MODERATE | Trp262Leu | | |
| SL2.50ch07 | 65456566 | T | G | missense_variant | MODERATE | Leu359Val | | |
| SL2.50ch07 | 65456817 | G | C | missense_variant | MODERATE | Glu412Asp | | |
| SL2.50ch07 | 65456843 | G | A | missense_variant | MODERATE | Arg421His | | |
| SL2.50ch07 | 65456879 | T | A | missense_variant | MODERATE | Ile433Asn | | |
| SL2.50ch07 | 65456993 | A | T | missense_variant | MODERATE | Glu471Val | | |
| SL2.50ch07 | 65457124 | T | G | missense_variant | MODERATE | Tyr515Asp | | |
| SL2.50ch07 | 65457277 | C | G | missense_variant | MODERATE | Leu566Val | | |
| SL2.50ch07 | 65457666 | T | G | missense_variant | MODERATE | Asp695Glu | | |
| SL2.50ch07 | 65457709 | A | G | missense_variant | MODERATE | Asn710Asp | | |
| SL2.50ch07 | 65458453 | AT | ATTATCTT | inframe_insertion | MODERATE | Asp255delinsGluAspAsn | Solyc07g062780.2 | E3 SUMO-protein ligase NSE2 (AHRD V1 **** C3KJP6_ANOFI)%3B contains Interpro domain(s) IPR004181 Zinc finger%2C MIZ-type |
| SL2.50ch07 | 65458493 | G | C | missense_variant | MODERATE | Ala242Gly | | |
| SL2.50ch07 | 65458739 | C | T | missense_variant | MODERATE | Arg186Lys | | |
| SL2.50ch07 | 65462391 | T | A | missense_variant | MODERATE | Arg144Trp | Solyc07g062790.1 | Homeobox leucine zipper protein (AHRD V1 ***- Q9XH73_PRUAR)%3B contains Interpro domain(s) IPR001356 Homeobox |
| SL2.50ch07 | 65462540 | A | T | missense_variant | MODERATE | Leu94Gln | | |
| SL2.50ch07 | 65465974 | AAACAACAA | AAACAA | inframe_deletion | MODERATE | Cys45del | Solyc07g062800.1 | Unknown Protein (AHRD V1) |
| SL2.50ch07 | 65468540 | C | G | missense_variant | MODERATE | Leu12Phe | | |
| SL2.50ch07 | 65469992 | T | C | missense_variant | MODERATE | Ile183Met | Solyc07g062810.2 | Branched-chain amino acid aminotransferase-like (AHRD V1 **** Q5NAM3_ORYSJ)%3B contains Interpro domain(s) IPR001544 Aminotransferase%2C class IV |
| SL2.50ch07 | 65470790 | T | A | missense_variant | MODERATE | Glu65Asp | | |
| SL2.50ch07 | 65470879 | C | G | missense_variant | MODERATE | Val36Leu | | |
| SL2.50ch07 | 65471864 | C | A | missense_variant | MODERATE | Val26Phe | | |
| SL2.50ch07 | 65473031 | G | A | missense_variant | MODERATE | Gly71Asp | Solyc07g062820.1 | Pentatricopeptide repeat protein PPR1106-17 (AHRD V1 ***- B6SWY6_MAIZE)%3B contains Interpro domain(s) IPR002885 Pentatricopeptide repeat |
| SL2.50ch07 | 65473116 | A | T | missense_variant | MODERATE | Glu99Asp | | |
| SL2.50ch07 | 65473458 | A | T | missense_variant | MODERATE | Leu213Phe | | |
| SL2.50ch07 | 65473517 | C | A | missense_variant | MODERATE | Ala233Glu | | |
| SL2.50ch07 | 65474125 | A | G | missense_variant | MODERATE | Thr436Ala | | |
| SL2.50ch07 | 65474148 | A | C | missense_variant | MODERATE | Glu443Asp | | |
| SL2.50ch07 | 65486951 | C | A | missense_variant | MODERATE | Thr246Asn | Solyc07g062840.2 | NAC domain protein IPR003441 (AHRD V1 ***- B9N1D6_POPTR)%3B contains Interpro domain(s) IPR003441 No apical meristem (NAM) protein |
| SL2.50ch07 | 65487055 | G | A | missense_variant | MODERATE | Ala281Thr | | |
| SL2.50ch07 | 65487073 | T | A | missense_variant | MODERATE | Ser287Thr | | |
| SL2.50ch07 | 65487160 | T | G | missense_variant | MODERATE | Ser316Ala | | |
| SL2.50ch07 | 65487206 | T | G | missense_variant | MODERATE | Val331Gly | | |
| SL2.50ch07 | 65493648 | C | T | missense_variant | MODERATE | Arg97Cys | Solyc07g062850.2 | Genomic DNA chromosome 3 BAC clone F4B12 (AHRD V1 ***- Q9LIL7_ARATH)%3B contains Interpro domain(s) IPR016024 Armadillo-type fold |
| SL2.50ch07 | 65494258 | C | G | missense_variant | MODERATE | Leu114Val | | |
| SL2.50ch07 | 65495573 | T | A | missense_variant | MODERATE | Leu172Met | | |
| SL2.50ch07 | 65495582 | G | A | missense_variant | MODERATE | Ala175Thr | | |
| SL2.50ch07 | 65495594 | G | T | missense_variant | MODERATE | Ala179Ser | | |
| SL2.50ch07 | 65496653 | C | T | missense_variant | MODERATE | Ser227Leu | | |
| SL2.50ch07 | 65498949 | T | A | missense_variant | MODERATE | Phe333Ile | | |
| SL2.50ch07 | 65498976 | A | G | missense_variant | MODERATE | Ile342Val | | |
| SL2.50ch07 | 65502383 | G | C | missense_variant | MODERATE | Ser69Thr | Solyc07g062860.2 | Genomic DNA chromosome 3 BAC clone F4B12 (AHRD V1 ***- Q9LIL7_ARATH) |
| SL2.50ch07 | 65523028 | T | G | missense_variant&splice_region_variant | MODERATE | Ser25Ala | Solyc07g062890.2 | Yippee zinc-binding-like protein (AHRD V1 *-** C1FDK9_9CHLO)%3B contains Interpro domain(s) IPR004910 Yippee-like protein |
| SL2.50ch07 | 65524686 | T | C | missense_variant | MODERATE | Met56Thr | | |
| SL2.50ch07 | 65538299 | G | A | missense_variant | MODERATE | Arg63Lys | Solyc07g062910.2 | Yippee family protein (AHRD V1 *-*- D7MTM3_ARALY)%3B contains Interpro domain(s) IPR004910 Yippee-like protein |
| SL2.50ch07 | 65538367 | G | T | missense_variant | MODERATE | Val86Leu | | |
| SL2.50ch07 | 65544918 | G | T | missense_variant | MODERATE | Arg141Leu | | |
| SL2.50ch07 | 65544926 | A | G | missense_variant | MODERATE | Lys144Glu | | |
| SL2.50ch07 | 65546756 | G | C | missense_variant | MODERATE | Glu186Asp | | |
| SL2.50ch07 | 65548265 | T | C | missense_variant | MODERATE | Leu7Ser | Solyc07g062920.2 | Genomic DNA chromosome 5 TAC clone K19P17 (AHRD V1 *-*- Q9FN33_ARATH) |
| SL2.50ch07 | 65549872 | A | G | missense_variant | MODERATE | Asn62Asp | | |
| SL2.50ch07 | 65549944 | T | C | missense_variant | MODERATE | Trp86Arg | | |
| SL2.50ch07 | 65550143 | G | A | missense_variant | MODERATE | Gly152Asp | | |
| SL2.50ch07 | 65550173 | G | A | missense_variant | MODERATE | Arg162His | | |
| SL2.50ch07 | 65550196 | A | G | missense_variant | MODERATE | Lys170Glu | | |
| SL2.50ch07 | 65550295 | T | A | missense_variant | MODERATE | Trp203Arg | | |
| SL2.50ch07 | 65550388 | C | T | missense_variant | MODERATE | Arg234Cys | | |
| SL2.50ch07 | 65550506 | C | G | missense_variant | MODERATE | Thr273Arg | | |
| SL2.50ch07 | 65550862 | C | T | missense_variant | MODERATE | Pro392Ser | | |
| SL2.50ch07 | 65550932 | A | C | missense_variant | MODERATE | Asp415Ala | | |
| SL2.50ch07 | 65551039 | T | G | missense_variant | MODERATE | Phe451Val | | |
| SL2.50ch07 | 65551085 | A | G | missense_variant | MODERATE | Gln466Arg | | |
| SL2.50ch07 | 65551117 | C | T | missense_variant | MODERATE | Pro477Ser | | |
| SL2.50ch07 | 65551312 | T | G | missense_variant | MODERATE | Leu542Val | | |
| SL2.50ch07 | 65551328 | C | T | missense_variant | MODERATE | Ala547Val | | |
| SL2.50ch07 | 65553293 | T | A | missense_variant | MODERATE | Thr373Ser | Solyc07g062930.2 | Ribosomal protein L11 methyltransferase-like protein (AHRD V1 **** Q9FN34_ARATH)%3B contains Interpro domain(s) IPR010456 Ribosomal L11 methyltransferase |
| SL2.50ch07 | 65553335 | T | A | missense_variant | MODERATE | Ile359Leu | | |
| SL2.50ch07 | 65554074 | C | G | missense_variant | MODERATE | Glu335Gln | | |
| SL2.50ch07 | 65554133 | C | T | missense_variant | MODERATE | Gly315Glu | | |
| SL2.50ch07 | 65555361 | C | T | missense_variant | MODERATE | Gly227Glu | | |
| SL2.50ch07 | 65557056 | TCT | TCTCTCACTGCT | inframe_insertion | MODERATE | Glu115_Glu117dup | | |
| SL2.50ch07 | 65557751 | C | T | missense_variant | MODERATE | Arg57His | | |
| SL2.50ch07 | 65557764 | T | G | missense_variant | MODERATE | Lys53Gln | | |
| SL2.50ch07 | 65559825 | T | A | missense_variant | MODERATE | Thr664Ser | Solyc07g062940.2 | Serine/threonine kinase 25 (AHRD V1 *-*- Q6V9V7_CHICK)%3B contains Interpro domain(s) IPR002290 Serine/threonine protein kinase |
| SL2.50ch07 | 65561601 | G | A | missense_variant | MODERATE | Pro540Ser | | |
| SL2.50ch07 | 65561669 | T | C | missense_variant | MODERATE | Glu517Gly | | |
| SL2.50ch07 | 65563239 | G | A | missense_variant | MODERATE | Thr405Ile | | |
| SL2.50ch07 | 65563323 | G | C | missense_variant | MODERATE | Ala377Gly | | |
| SL2.50ch07 | 65563594 | C | T | missense_variant | MODERATE | Arg310Lys | | |
| SL2.50ch07 | 65564248 | T | C | missense_variant | MODERATE | Asn269Ser | | |
| SL2.50ch07 | 65588315 | G | A | missense_variant | MODERATE | Ala285Thr | Solyc07g062950.2 | BHLH transcription factor (AHRD V1 *-*- Q9XIN0_ARATH) |
| SL2.50ch07 | 65592247 | A | G | missense_variant | MODERATE | Leu162Ser | Solyc07g062960.1 | Unknown Protein (AHRD V1) |
| SL2.50ch07 | 65592287 | A | T | missense_variant | MODERATE | Phe149Ile | | |
| SL2.50ch07 | 65592322 | G | A | missense_variant | MODERATE | Ala137Val | | |
| SL2.50ch07 | 65598058 | C | T | missense_variant | MODERATE | Ala98Val | Solyc07g062970.2 | Serine/threonine phosphatase family protein (AHRD V1 **** C1E0J1_9CHLO)%3B contains Interpro domain(s) IPR015655 Protein phosphatase 2C |
| SL2.50ch07 | 65599611 | G | T | missense_variant | MODERATE | Thr124Asn | Solyc07g062980.2 | Squamosa promoter binding-like protein (AHRD V1 ***- Q52UU1_9ROSI)%3B contains Interpro domain(s) IPR004333 Transcription factor%2C SBP-box |
| SL2.50ch07 | 65599653 | TCCCC | TC | inframe_deletion | MODERATE | Gly109del | | |
| SL2.50ch07 | 65603933 | C | T | missense_variant | MODERATE | Asp144Asn | Solyc07g062990.1 | Unknown Protein (AHRD V1) |
| SL2.50ch07 | 65604071 | C | T | missense_variant | MODERATE | Val98Ile | | |
| SL2.50ch07 | 65604149 | A | T | missense_variant | MODERATE | Cys72Ser | | |
| SL2.50ch07 | 65605759 | A | C | missense_variant | MODERATE | Val7Gly | | |
| SL2.50ch07 | 65608925 | G | C | missense_variant | MODERATE | Ala1012Gly | Solyc07g063000.2 | Receptor like kinase%2C RLK |
| SL2.50ch07 | 65609072 | G | A | missense_variant | MODERATE | Ala963Val | | |
| SL2.50ch07 | 65610189 | G | A | missense_variant | MODERATE | Leu591Phe | | |
| SL2.50ch07 | 65611890 | T | C | missense_variant | MODERATE | Asn51Asp | | |
| SL2.50ch07 | 65611971 | T | A | missense_variant | MODERATE | Thr24Ser | | |
| SL2.50ch07 | 65612007 | C | G | missense_variant | MODERATE | Val12Leu | | |
| SL2.50ch07 | 65621582 | T | A | missense_variant | MODERATE | Asp79Glu | Solyc07g063010.2 | Fatty acid oxidation complex subunit alpha (AHRD V1 **-- FADB_ACIBC)%3B contains Interpro domain(s) IPR006176 3-hydroxyacyl-CoA dehydrogenase%2C NAD binding |
| SL2.50ch07 | 65621591 | G | T | missense_variant | MODERATE | Gln82His | | |
| SL2.50ch07 | 65622853 | G | C | missense_variant | MODERATE | Cys166Ser | | |
| SL2.50ch07 | 65624370 | G | C | missense_variant | MODERATE | Val12Leu | Solyc07g063020.1 | CT099 (Fragment) (AHRD V1 *-*- Q4KR07_SOLPE)%3B contains Interpro domain(s) IPR008972 Cupredoxin |
| SL2.50ch07 | 65624924 | T | G | missense_variant | MODERATE | Val162Gly | | |
| SL2.50ch07 | 65625175 | A | G | missense_variant | MODERATE | Asn246Asp | | |
| SL2.50ch07 | 65625207 | T | G | missense_variant | MODERATE | Ile256Met | | |
| SL2.50ch07 | 65625277 | T | A | missense_variant | MODERATE | Ser280Thr | | |
| SL2.50ch07 | 65625430 | G | A | missense_variant | MODERATE | Ala331Thr | | |
| SL2.50ch07 | 65625514 | T | A | missense_variant | MODERATE | Ser359Thr | | |
| SL2.50ch07 | 65625569 | T | C | missense_variant | MODERATE | Leu377Pro | | |
| SL2.50ch07 | 65625577 | T | G | missense_variant | MODERATE | Ser380Ala | | |
| SL2.50ch07 | 65625865 | C | G | missense_variant | MODERATE | Ala407Gly | | |
| SL2.50ch07 | 65625979 | T | C | missense_variant | MODERATE | Ile445Thr | | |
| SL2.50ch07 | 65625993 | G | A | missense_variant | MODERATE | Val450Met | | |
| SL2.50ch07 | 65626314 | C | T | missense_variant | MODERATE | Pro557Ser | | |
| SL2.50ch07 | 65626338 | T | C | missense_variant | MODERATE | Tyr565His | | |
| SL2.50ch07 | 65626599 | A | G | missense_variant | MODERATE | Lys652Glu | | |
| SL2.50ch07 | 65626719 | C | T | missense_variant | MODERATE | His692Tyr | | |
| SL2.50ch07 | 65626761 | G | A | missense_variant | MODERATE | Asp706Asn | | |
| SL2.50ch07 | 65626813 | T | C | missense_variant | MODERATE | Leu723Pro | | |
| SL2.50ch07 | 65626875 | A | C | missense_variant | MODERATE | Ser744Arg | | |
| SL2.50ch07 | 65626884 | A | G | missense_variant | MODERATE | Asn747Asp | | |
| SL2.50ch07 | 65627013 | A | G | missense_variant | MODERATE | Ile790Val | | |
| SL2.50ch07 | 65627041 | A | G | missense_variant | MODERATE | Asn799Ser | | |
| SL2.50ch07 | 65627170 | G | A | missense_variant | MODERATE | Gly842Glu | | |
| SL2.50ch07 | 65627334 | G | A | missense_variant | MODERATE | Val897Ile | | |
| SL2.50ch07 | 65627372 | TCCAATTTCATCTTCCCCA | TCCAATTTCATCTTCCCCAATTTCATCTTCCCCA | inframe_insertion | MODERATE | Ile911_Pro915dup | | |
| SL2.50ch07 | 65627437 | A | G | missense_variant | MODERATE | Asn931Ser | | |
| SL2.50ch07 | 65627517 | G | T | missense_variant | MODERATE | Ala958Ser | | |
| SL2.50ch07 | 65627593 | A | T | missense_variant | MODERATE | Asn983Ile | | |
| SL2.50ch07 | 65630895 | A | G | missense_variant | MODERATE | Asn43Ser | Solyc07g063030.1 | Rapid alkalinization factor 2 (AHRD V1 **-- Q6TF28_SOLCH)%3B contains Interpro domain(s) IPR008801 Rapid ALkalinization Factor |
| SL2.50ch07 | 65630919 | G | T | missense_variant | MODERATE | Trp51Leu | | |
| SL2.50ch07 | 65631009 | A | G | missense_variant | MODERATE | Gln81Arg | | |
| SL2.50ch07 | 65631063 | A | G | missense_variant | MODERATE | His99Arg | | |

**Supplemental Table S4 (cont.). POLYMORPHISMS WITH A MODERATE EFFECT ON GENE FUNCTION FOR S.PENNELLII VS S.LYCOPERSICUM (SL2.50) WITHIN THE *ORG* REGION.** The variant information was obtained from the S.pennellii resequencing dataset (Aflitos et al. 2014).

| | | | | | | | | |
|---|---|---|---|---|---|---|---|---|
| SL2.50ch07 | 65633577 | T | G | missense_variant | MODERATE | Ser87Ala | Solyc07g063040.2 | Enolase-phosphatase E-1 (AHRD V1 *-*- A8UQN2_9AQUI)%3B contains Interpro domain(s) IPR017714 Methylthioribulose-1-phosphate dehydratase |
| SL2.50ch07 | 65637566 | T | G | missense_variant | MODERATE | Ser391Ala | | |
| SL2.50ch07 | 65637576 | A | G | missense_variant | MODERATE | Glu394Gly | | |
| SL2.50ch07 | 65639854 | A | C | missense_variant | MODERATE | Leu100Phe | Solyc07g063050.1 | Unknown Protein (AHRD V1) |
| SL2.50ch07 | 65644707 | G | A | missense_variant | MODERATE | Val40Ile | Solyc07g063060.1 | Os01g0101200 protein (Fragment) (AHRD V1 *-*- Q0JRH3_ORYSJ) |
| SL2.50ch07 | 65644761 | A | G | missense_variant | MODERATE | Met58Val | | |
| SL2.50ch07 | 65644797 | A | G | missense_variant | MODERATE | Thr70Ala | | |
| SL2.50ch07 | 65644803 | T | C | missense_variant | MODERATE | Cys72Arg | | |
| SL2.50ch07 | 65645349 | A | G | missense_variant | MODERATE | Lys114Glu | | |
| SL2.50ch07 | 65650033 | A | T | missense_variant | MODERATE | Ser95Thr | Solyc07g063070.1 | DNA-binding WRKY VQ (AHRD V1 ***- Q2HTS9_MEDTR)%3B contains Interpro domain(s) IPR008889 VQ |
| SL2.50ch07 | 65650060 | A | C | missense_variant | MODERATE | Ser86Ala | | |
| SL2.50ch07 | 65658471 | A | T | missense_variant | MODERATE | Ser705Thr | Solyc07g063080.2 | Dynamin family protein (AHRD V1 ***- D7KKE2_ARALY)%3B contains Interpro domain(s) IPR001401 Dynamin%2C GTPase region |
| SL2.50ch07 | 65658951 | G | A | missense_variant | MODERATE | Pro545Ser | | |
| SL2.50ch07 | 65659747 | T | A | missense_variant | MODERATE | Ile419Leu | | |
| SL2.50ch07 | 65661441 | T | G | missense_variant | MODERATE | Glu135Ala | | |
| SL2.50ch07 | 65661485 | A | T | missense_variant&splice_region_variant | MODERATE | Asp120Glu | | |
| SL2.50ch07 | 65661994 | A | C | missense_variant | MODERATE | Asp113Glu | | |
| SL2.50ch07 | 65670962 | G | A | missense_variant | MODERATE | Pro610Leu | Solyc07g063100.2 | Proton pump interactor 1 (AHRD V1 **-- D5L6G0_SOLTU) |
| SL2.50ch07 | 65671500 | C | T | missense_variant | MODERATE | Met488Ile | | |
| SL2.50ch07 | 65671549 | A | G | missense_variant | MODERATE | Ile472Thr | | |
| SL2.50ch07 | 65674291 | CCTT | CCTTCTT | inframe_insertion | MODERATE | Pro50dup | | |
| SL2.50ch07 | 65692406 | T | A | missense_variant | MODERATE | Met471Leu | Solyc07g063120.2 | WD-40 repeat protein (AHRD V1 *--- B7KCU4_CYAP7)%3B contains Interpro domain(s) IPR020472 G-protein beta WD-40 repeat%2C region |
| SL2.50ch07 | 65692475 | A | G | missense_variant | MODERATE | Ser448Pro | | |
| SL2.50ch07 | 65692654 | T | C | missense_variant | MODERATE | Lys388Arg | | |
| SL2.50ch07 | 65692729 | C | T | missense_variant | MODERATE | Arg363Lys | | |
| SL2.50ch07 | 65692749 | ACCCC | ACCCCCCC | inframe_insertion | MODERATE | Arg355dup | | |
| SL2.50ch07 | 65694927 | C | T | missense_variant | MODERATE | Ala238Thr | | |
| SL2.50ch07 | 65694942 | A | G | missense_variant | MODERATE | Ser233Pro | | |
| SL2.50ch07 | 65695100 | C | T | missense_variant | MODERATE | Val216Ile | | |
| SL2.50ch07 | 65695202 | TTTCTTC | TTTCTTCTTC | inframe_insertion | MODERATE | Leu179dup | | |
| SL2.50ch07 | 65695288 | A | G | missense_variant | MODERATE | Ile153Thr | | |
| SL2.50ch07 | 65695387 | G | T | missense_variant | MODERATE | Thr120Asn | | |
| SL2.50ch07 | 65695649 | A | C | missense_variant | MODERATE | Cys33Gly | | |
| SL2.50ch07 | 65695726 | A | C | missense_variant | MODERATE | Ile7Arg | | |
| SL2.50ch07 | 65699564 | G | T | missense_variant | MODERATE | Pro571Gln | Solyc07g063130.2 | Cell division protein kinase 10 (AHRD V1 *-*- D3TRH8_GLOMM)%3B contains Interpro domain(s) IPR002290 Serine/threonine protein kinase |
| SL2.50ch07 | 65699651 | A | G | missense_variant | MODERATE | Val542Ala | | |
| SL2.50ch07 | 65701377 | A | T | missense_variant | MODERATE | Cys423Ser | | |
| SL2.50ch07 | 65701392 | A | C | missense_variant | MODERATE | Cys418Gly | | |
| SL2.50ch07 | 65702650 | A | C | missense_variant | MODERATE | Ser338Ala | | |
| SL2.50ch07 | 65702923 | T | A | missense_variant | MODERATE | Met247Leu | | |
| SL2.50ch07 | 65706934 | T | C | missense_variant | MODERATE | Ile15Val | | |
| SL2.50ch07 | 65712553 | G | A | missense_variant | MODERATE | Arg31Gln | Solyc07g063140.1 | Aspartyl/glutamyl-tRNA(Asn/Gln) amidotransferase subunit C (AHRD V1 ***- A3IGR0_9CHRO)%3B contains Interpro domain(s) IPR003837 Glu-tRNAGln amidotransferase%2C C subunit |
| SL2.50ch07 | 65712595 | T | C | missense_variant | MODERATE | Val45Ala | | |
| SL2.50ch07 | 65712693 | G | A | missense_variant | MODERATE | Val78Ile | | |
| SL2.50ch07 | 65712756 | A | G | missense_variant | MODERATE | Asn99Asp | | |
| SL2.50ch07 | 65712803 | G | T | missense_variant | MODERATE | Glu114Asp | | |
| SL2.50ch07 | 65718720 | G | A | missense_variant | MODERATE | Asp12Asn | Solyc07g063150.2 | Os11g0198100 protein (Fragment) (AHRD V1 *--- Q0IU20_ORYSJ)%3B contains Interpro domain(s) IPR006852 Protein of unknown function DUF616 |
| SL2.50ch07 | 65718949 | T | G | missense_variant | MODERATE | Asp43Glu | | |
| SL2.50ch07 | 65719105 | A | C | missense_variant | MODERATE | Glu95Asp | | |
| SL2.50ch07 | 65721812 | G | T | missense_variant | MODERATE | Gln248His | | |
| SL2.50ch07 | 65722574 | T | G | missense_variant | MODERATE | Val430Gly | | |
| SL2.50ch07 | 65724283 | GAGAAGAAGAAGAAGAAGAA | GAGAAGAAGAAGAA | disruptive_inframe_deletion | MODERATE | Lys37_Lys38del | Solyc07g063160.2 | Cytochrome c oxidase copper chaperone family protein (AHRD V1 *-*- D7KKD3_ARALY)%3B contains Interpro domain(s) IPR007745 Cytochrome c oxidase copper chaperone |
| SL2.50ch07 | 65726273 | G | C | missense_variant | MODERATE | Val59Leu | | |
| SL2.50ch07 | 65726334 | T | C | missense_variant | MODERATE | Leu79Ser | | |
| SL2.50ch07 | 65727054 | T | C | missense_variant | MODERATE | Phe7Leu | Solyc07g063170.1 | Unknown Protein (AHRD V1) |
| SL2.50ch07 | 65727090 | A | C | missense_variant | MODERATE | Ile19Leu | | |
| SL2.50ch07 | 65727104 | G | T | missense_variant | MODERATE | Gln23His | | |
| SL2.50ch07 | 65727145 | G | A | missense_variant | MODERATE | Gly37Asp | | |
| SL2.50ch07 | 65727154 | T | A | missense_variant | MODERATE | Ile40Asn | | |
| SL2.50ch07 | 65727273 | G | T | missense_variant | MODERATE | Gly80Cys | | |
| SL2.50ch07 | 65727341 | G | A | missense_variant | MODERATE | Met102Ile | | |
| SL2.50ch07 | 65727426 | G | A | missense_variant | MODERATE | Gly131Arg | | |
| SL2.50ch07 | 65727450 | T | G | missense_variant | MODERATE | Tyr139Asp | | |
| SL2.50ch07 | 65729694 | A | G | missense_variant | MODERATE | Tyr63His | Solyc07g063180.2 | Cytoplasmic dynein light chain (AHRD V1 *-*- D3BU56_POLPA)%3B contains Interpro domain(s) IPR001372 Dynein light chain%2C type 1 and 2 |
| SL2.50ch07 | 65731436 | A | C | missense_variant | MODERATE | Gln23Pro | Solyc07g063190.2 | Thioredoxin (AHRD V1 ***- A7LNX7_9CARY)%3B contains Interpro domain(s) IPR005746 Thioredoxin |
| SL2.50ch07 | 65731469 | T | C | missense_variant | MODERATE | Phe34Ser | | |
| SL2.50ch07 | 65731479 | C | A | missense_variant | MODERATE | Ser37Arg | | |
| SL2.50ch07 | 65733205 | A | G | missense_variant | MODERATE | Thr81Ala | | |
| SL2.50ch07 | 65737800 | G | A | missense_variant | MODERATE | Glu13Lys | Solyc07g063200.1 | RING finger protein 5 (AHRD V1 *-*- B6TF06_MAIZE)%3B contains Interpro domain(s) IPR018957 Zinc finger%2C C3HC4 RING-type |
| SL2.50ch07 | 65738067 | C | A | missense_variant | MODERATE | Pro102Thr | | |
| SL2.50ch07 | 65738197 | G | A | missense_variant | MODERATE | Arg145Gln | | |
| SL2.50ch07 | 65738446 | G | T | missense_variant | MODERATE | Cys228Phe | | |
| SL2.50ch07 | 65743353 | T | C | missense_variant | MODERATE | Val242Ala | Solyc07g063210.2 | Expansin S2 (AHRD V1 ***- Q39626_CUCSA)%3B contains Interpro domain(s) IPR002963 Expansin |
| SL2.50ch07 | 65744928 | CGCT | C | inframe_deletion | MODERATE | Ala49del | Solyc07g063220.2 | Pentatricopeptide repeat-containing protein (AHRD V1 ***- D7MEQ2_ARALY)%3B contains Interpro domain(s) IPR002885 Pentatricopeptide repeat |
| SL2.50ch07 | 65746824 | A | G | missense_variant | MODERATE | Asn246Ser | | |
| SL2.50ch07 | 65747181 | A | G | missense_variant | MODERATE | His365Arg | | |
| SL2.50ch07 | 65747562 | T | C | missense_variant | MODERATE | Ile492Thr | | |
| SL2.50ch07 | 65747581 | C | A | missense_variant | MODERATE | Asn498Lys | | |
| SL2.50ch07 | 65750217 | A | G | missense_variant | MODERATE | Thr44Ala | Solyc07g063230.2 | Choline transporter like family (AHRD V1 ***- C1E2U6_9CHLO)%3B contains Interpro domain(s) IPR007603 Protein of unknown function DUF580 |
| SL2.50ch07 | 65751156 | C | A | missense_variant | MODERATE | Ser67Tyr | | |
| SL2.50ch07 | 65752401 | T | C | missense_variant | MODERATE | Phe267Leu | | |
| SL2.50ch07 | 65754313 | G | A | missense_variant | MODERATE | Ser539Asn | | |
| SL2.50ch07 | 65754334 | A | G | missense_variant | MODERATE | Gln546Arg | | |
| SL2.50ch07 | 65754349 | T | C | missense_variant | MODERATE | Ile551Thr | | |
| SL2.50ch07 | 65760195 | C | A | missense_variant | MODERATE | Arg309Leu | Solyc07g063240.2 | Genomic DNA chromosome 3 P1 clone MJK13 (AHRD V1 **-- Q9LUQ3_ARATH)%3B contains Interpro domain(s) IPR013899 Region of unknown function DUF1771 |
| SL2.50ch07 | 65760508 | A | G | missense_variant | MODERATE | Val248Ala | | |
| SL2.50ch07 | 65763097 | T | C | missense_variant | MODERATE | Ser206Gly | | |
| SL2.50ch07 | 65763595 | G | T | missense_variant | MODERATE | Ala170Asp | | |
| SL2.50ch07 | 65763797 | T | C | missense_variant | MODERATE | Lys103Glu | | |
| SL2.50ch07 | 65763884 | C | G | missense_variant | MODERATE | Glu74Gln | | |
| SL2.50ch07 | 65769266 | A | C,T | missense_variant | MODERATE | Thr38Pro; Thr38Ser | Solyc07g063250.2 | Unknown Protein (AHRD V1) |
| SL2.50ch07 | 65772102 | G | C | missense_variant | MODERATE | Thr540Ser | Solyc07g063260.2 | MLO-like protein 14 (AHRD V1 ***- B6SWK8_MAIZE)%3B contains Interpro domain(s) IPR004326 Mlo-related protein |
| SL2.50ch07 | 65772187 | AATGATGATGAT | AATGATGAT | inframe_deletion | MODERATE | His509del | | |
| SL2.50ch07 | 65777726 | T | A | missense_variant | MODERATE | Lys132Asn | | |
| SL2.50ch07 | 65777794 | T | C | missense_variant | MODERATE | Lys110Glu | | |
| SL2.50ch07 | 65785541 | T | C | missense_variant | MODERATE | Lys486Arg | Solyc07g063270.2 | Nucleolar GTP-binding protein 2 (AHRD V1 **** B6TMG1_MAIZE)%3B contains Interpro domain(s) IPR012971 NGP1%2C N-terminal |
| SL2.50ch07 | 65792828 | T | C | missense_variant | MODERATE | Asp74Gly | Solyc07g063280.1 | Nucleolar GTP-binding protein 2 (AHRD V1 ***- B6TMG1_MAIZE)%3B contains Interpro domain(s) IPR012971 NGP1%2C N-terminal |
| SL2.50ch07 | 65793947 | G | C | missense_variant | MODERATE | Ser22Cys | | |
| SL2.50ch07 | 65795069 | A | ATGGACCACC | missense_variant | MODERATE | Thr75_Pro77dup | Solyc07g063290.2 | Unknown Protein (AHRD V1)%3B contains Interpro domain(s) IPR019481 Transcription factor TFIIIC%2C tau55-related |
| SL2.50ch07 | 65795640 | T | A | missense_variant&splice_region_variant | MODERATE | Lys73Asn | | |
| SL2.50ch07 | 65800467 | G | A | missense_variant | MODERATE | Gly44Glu | Solyc07g063300.2 | Regulator of chromosome condensation (RCC1)-like protein (AHRD V1 **-- D0NEM9_PHYIN)%3B contains Interpro domain(s) IPR009091 Regulator of chromosome condensation/beta-lactamase-inhibitor protein II |
| SL2.50ch07 | 65800587 | C | A | missense_variant | MODERATE | Thr84Lys | | |
| SL2.50ch07 | 65801340 | G | A | missense_variant | MODERATE | Arg156Gln | | |
| SL2.50ch07 | 65802271 | T | A | missense_variant | MODERATE | Phe193Tyr | | |
| SL2.50ch07 | 65805342 | G | C | missense_variant | MODERATE | Glu55Gln | Solyc07g063310.2 | LanC-like protein 2 (AHRD V1 ***- B6TCV9_MAIZE)%3B contains Interpro domain(s) IPR020464 LanC-like protein%2C eukaryotic |
| SL2.50ch07 | 65805517 | T | C | missense_variant | MODERATE | Phe83Leu | | |
| SL2.50ch07 | 65806446 | C | A | missense_variant | MODERATE | Thr149Lys | | |
| SL2.50ch07 | 65806540 | T | G | missense_variant | MODERATE | Leu156Arg | | |
| SL2.50ch07 | 65808296 | T | G | missense_variant | MODERATE | Val19Gly | Solyc07g063320.2 | LanC-like protein 2 (AHRD V1 ***- B6TCV9_MAIZE)%3B contains Interpro domain(s) IPR020464 LanC-like protein%2C eukaryotic |
| SL2.50ch07 | 65808323 | T | C | missense_variant | MODERATE | Phe28Ser | | |
| SL2.50ch07 | 65809328 | C | G | missense_variant | MODERATE | Thr69Arg | | |
| SL2.50ch07 | 65811169 | G | A | missense_variant | MODERATE | Glu344Lys | | |
| SL2.50ch07 | 65816493 | C | T | missense_variant | MODERATE | Val490Ile | Solyc07g063330.2 | WD-40 repeat family protein (AHRD V1 ***- D7L4T9_ARALY)%3B contains Interpro domain(s) IPR020472 G-protein beta WD-40 repeat%2C region |
| SL2.50ch07 | 65817071 | G | A | missense_variant | MODERATE | Ala297Val | | |
| SL2.50ch07 | 65817250 | C | A | missense_variant | MODERATE | Lys237Asn | | |
| SL2.50ch07 | 65817339 | G | C | missense_variant | MODERATE | Gln208Glu | | |
| SL2.50ch07 | 65817366 | G | T | missense_variant | MODERATE | Pro199Thr | | |
| SL2.50ch07 | 65818094 | A | T | missense_variant | MODERATE | Phe14Tyr | | |
| SL2.50ch07 | 65831877 | T | C | missense_variant | MODERATE | Asp39Gly | Solyc07g063340.2 | COSII_At1g13380 (Fragment) (AHRD V1 *-*- C0KEY3_SOLDU)%3B contains Interpro domain(s) IPR009606 Protein of unknown function DUF1218 |
| SL2.50ch07 | 65831890 | T | G | missense_variant | MODERATE | Lys35Gln | | |
| SL2.50ch07 | 65831897 | A | T | missense_variant | MODERATE | Asn32Lys | | |
| SL2.50ch07 | 65839573 | T | G | missense_variant | MODERATE | His1039Pro | Solyc07g063350.2 | COP1-Interacting ProteinI 7 (CIP7) (Fragment) (AHRD V1 *-*- Q7GDB1_ARATH) |
| SL2.50ch07 | 65839732 | C | T | missense_variant | MODERATE | Arg986His | | |
| SL2.50ch07 | 65839768 | C | T | missense_variant | MODERATE | Ser974Asn | | |
| SL2.50ch07 | 65839952 | C | A | missense_variant | MODERATE | Val913Leu | | |
| SL2.50ch07 | 65840012 | C | A | missense_variant | MODERATE | Asp893Tyr | | |
| SL2.50ch07 | 65840436 | T | G | missense_variant | MODERATE | Thr785Pro | | |
| SL2.50ch07 | 65840480 | C | T | missense_variant | MODERATE | Arg770Lys | | |
| SL2.50ch07 | 65840527 | C | A | missense_variant | MODERATE | Glu754Asp | | |
| SL2.50ch07 | 65840592 | C | T | missense_variant | MODERATE | Glu733Lys | | |
| SL2.50ch07 | 65840701 | C | A | missense_variant | MODERATE | Glu696Asp | | |
| SL2.50ch07 | 65840882 | T | C | missense_variant | MODERATE | His636Arg | | |
| SL2.50ch07 | 65841014 | T | G | missense_variant | MODERATE | Asp592Ala | | |
| SL2.50ch07 | 65841789 | T | G | missense_variant | MODERATE | Thr334Pro | | |
| SL2.50ch07 | 65841870 | G | T | missense_variant | MODERATE | Leu307Met | | |
| SL2.50ch07 | 65842261 | T | C | missense_variant | MODERATE | Arg216Gly | | |
| SL2.50ch07 | 65843371 | C | G | missense_variant | MODERATE | Cys70Ser | | |
| SL2.50ch07 | 65843644 | T | A | missense_variant | MODERATE | Tyr9Phe | | |
| SL2.50ch07 | 65849958 | G | A | missense_variant | MODERATE | Asp90Asn | Solyc07g063360.1 | Tir%2C resistance protein fragment |
| SL2.50ch07 | 65850135 | C | G | missense_variant | MODERATE | Gln149Glu | | |
| SL2.50ch07 | 65851348 | A | G | missense_variant | MODERATE | Phe494Leu | Solyc07g063370.2 | Beta-glucosidase (AHRD V1 **** B6SYQ7_MAIZE)%3B contains Interpro domain(s) IPR001360 Glycoside hydrolase%2C family 1 |
| SL2.50ch07 | 65851634 | AT | ATATCCTCCTTCTTGGT | inframe_insertion | MODERATE | Gly431_Asn435dup | | |
| SL2.50ch07 | 65853212 | C | T | missense_variant | MODERATE | Val240Ile | | |
| SL2.50ch07 | 65853245 | C | A | missense_variant | MODERATE | Gly229Cys | | |
| SL2.50ch07 | 65853356 | C | T | missense_variant | MODERATE | Asp192Asn | | |
| SL2.50ch07 | 65854994 | T | C | missense_variant | MODERATE | Ser20Gly | | |
| SL2.50ch07 | 65862593 | G | C | missense_variant | MODERATE | His493Asp | Solyc07g063390.2 | Beta-glucosidase (AHRD V1 **** B6SYQ7_MAIZE)%3B contains Interpro domain(s) IPR001360 Glycoside hydrolase%2C family 1 |
| SL2.50ch07 | 65862637 | G | T | missense_variant | MODERATE | Thr478Asn | | |
| SL2.50ch07 | 65863263 | T | A | missense_variant | MODERATE | Asn351Tyr | | |
| SL2.50ch07 | 65863354 | C | A | missense_variant | MODERATE | Arg320Ser | | |
| SL2.50ch07 | 65863597 | T | A | missense_variant | MODERATE | Met263Leu | | |
| SL2.50ch07 | 65864051 | G | C | missense_variant | MODERATE | Ile171Met | | |
| SL2.50ch07 | 65864125 | C | G | missense_variant | MODERATE | Ala147Pro | | |
| SL2.50ch07 | 65864578 | G | C | missense_variant | MODERATE | Thr75Arg | | |
| SL2.50ch07 | 65865655 | A | T | missense_variant | MODERATE | Ser6Thr | | |
| **SL2.50ch07** | **65865655** | | | **RIGHT BORDER - end of introgression in IL7-3** | | | | |

**Supplemental Table S5. POLYMORPHISMS WITH A HIGH EFFECT ON GENE FUNCTION FOR S.PENNELLII VS S.LYCOPERSICUM (SL2.50) WITHIN THE *ORG* REGION.** The variant information was obtained from the S.pennellii resequencing dataset (Aflitos et al. 2014).

| Chromosome | Position | Ref allele | Alt allele | Variant type | Impact | HGVS-nomenclatre | Gene | Protein description |
|---|---|---|---|---|---|---|---|---|
| **SL2.50ch07** | **65444176** | **A** | **G** | **LEFT BORDER** | | | | |
| SL2.50ch07 | 65454205 | GTTTT | GTTT | frameshift_variant | HIGH | Phe1101fs | Solyc07g062760.1 | 26S protease regulatory subunit 4 homolog (AHRD V1 *--- PRS4_ORYSJ)%3B contains Interpro domain(s) IPR003959 ATPase%2C AAA-type%2C core |
| SL2.50ch07 | 65472098 | T | A | splice_acceptor_variant&intron_variant | HIGH | | Solyc07g062810.2 | Branched-chain amino acid aminotransferase-like (AHRD V1 **** Q5NAM3_ORYSJ)%3B contains Interpro domain(s) IPR001544 Aminotransferase%2C class IV |
| SL2.50ch07 | 65475767 | C | A | stop_gained | HIGH | Glu29* | Solyc07g062830.1 | Unknown Protein (AHRD V1) |
| SL2.50ch07 | 65476127 | AGGG | AGG | frameshift_variant | HIGH | Ser12fs | | |
| SL2.50ch07 | 65498996 | CA | CAGA | frameshift_variant | HIGH | Lys349fs | Solyc07g062850.2 | Genomic DNA chromosome 3 BAC clone F4B12 (AHRD V1 ***- Q9LIL7_ARATH)%3B contains Interpro domain(s) IPR016024 Armadillo-type fold |
| SL2.50ch07 | 65499036 | T | C | stop_lost | HIGH | Ter362Glnext*? | | |
| SL2.50ch07 | 65546998 | ATT | AT | frameshift_variant | HIGH | Tyr215fs | Solyc07g062910.2 | Yippee family protein (AHRD V1 *-*- D7MTM3_ARALY)%3B contains Interpro domain(s) IPR004910 Yippee-like protein |
| SL2.50ch07 | 65603906 | CTTTTTT | CTTTTT | frameshift_variant | HIGH | Lys151fs | Solyc07g062990.1 | Unknown Protein (AHRD V1) |
| SL2.50ch07 | 65604247 | C | T | splice_acceptor_variant&intron_variant | HIGH | | | |
| SL2.50ch07 | 65642125 | T | A | start_lost | HIGH | Met1? | Solyc07g063060.1 | Os01g0101200 protein (Fragment) (AHRD V1 *-*- Q0JRH3_ORYSJ) |
| SL2.50ch07 | 65645037 | C | T | stop_gained | HIGH | Gln89* | | |
| SL2.50ch07 | 65645392 | A | G | stop_lost&splice_region_variant | HIGH | Ter128Trpext*? | | |
| SL2.50ch07 | 65706927 | G | T | stop_gained&splice_region_variant | HIGH | Ser17* | Solyc07g063130.2 | Cell division protein kinase 10 (AHRD V1 *-*- D3TRH8_GLOMM)%3B contains Interpro domain(s) IPR002290 Serine/threonine protein kinase |
| SL2.50ch07 | 65724259 | CTTTT | CTTT | frameshift_variant | HIGH | Phe25fs | Solyc07g063160.2 | Cytochrome c oxidase copper chaperone family protein (AHRD V1 *-*- D7KKD3_ARALY)%3B contains Interpro domain(s) IPR007745 Cytochrome c oxidase copper chaperone |
| SL2.50ch07 | 65724314 | T | A | stop_gained | HIGH | Leu42* | | |
| SL2.50ch07 | 65747845 | C | T | stop_gained | HIGH | Arg531* | Solyc07g063220.2 | Pentatricopeptide repeat-containing protein (AHRD V1 ***- D7MEQ2_ARALY)%3B contains Interpro domain(s) IPR002885 Pentatricopeptide repeat |
| SL2.50ch07 | 65795069 | A | AT | frameshift_variant | HIGH | Glu78fs | Solyc07g063290.2 | Unknown Protein (AHRD V1)%3B contains Interpro domain(s) IPR019481 Transcription factor TFIIIC%2C tau55-related |
| SL2.50ch07 | 65810291 | T | TC | splice_acceptor_variant&intron_variant | HIGH | | Solyc07g063320.2 | LanC-like protein 2 (AHRD V1 ***- B6TCV9_MAIZE)%3B contains Interpro domain(s) IPR020464 LanC-like protein%2C eukaryotic |
| **SL2.50ch07** | **66374044** | | | **RIGHT BORDER - end of introgression in IL7-3** | | | | |

**Supplemental Table S6. Coding sequences of S. lycopersicum (SL2.50) not producing a match with S. pennellii CDS.** The coding sequences (CDS) of S.pennellii (Bolger et al. 2014) were queried against the CDS of S.lycopersicum SL2.50 using BLAST+ and the CDS of S.lycopersicum that did not match with S. pennellii are listed here.

| # | Slyc gene | Annotation |
|---|---|---|
| 1 | Solyc07g062800.1.1 | Note=Unknown Protein (AHRD V1);eugene_evidence_code=10F0H0E0IEG;from_BOGAS=1;length=2693;nb_exon=2 |
| 2 | Solyc07g062830.1.1 | Note=Unknown Protein (AHRD V1);eugene_evidence_code=10F0H0E0IEG;from_BOGAS=1;length=418;nb_exon=2 |
| 3 | Solyc07g062900.2.1 | Note=Yippee family protein (AHRD V1 ***- D7MTM3_ARALY)%3B contains Interpro domain(s) IPR004910 Yippee-like protein ;from_BOGAS=1;length=4569;nb_exon=3 |
| 4 | Solyc07g063250.2.1 | Note=Unknown Protein (AHRD V1);from_BOGAS=1;length=2537;nb_exon=3 |
| 5 | Solyc07g063280.1.1 | Note=Nucleolar GTP-binding protein 2 (AHRD V1 ***- B6TMG1_MAIZE)%3B contains Interpro domain(s) IPR012971 NGP1%2C N-terminal ;eugene_evidence_code=10F0H1E0IEG;from_BOGAS=1;length=1847;nb_exon=3 |
| 6 | Solyc07g063360.1.1 | Note=Tir%2C resistance protein fragment;Ontology_term=GO:0031224;eugene_evidence_code=10F0H1E0IEG;from_BOGAS=1;interpro2go_term=GO:0031224;length=1079;nb_exon=2 |
